# Identification of dynamic models of microbial communities

**DOI:** 10.1101/2025.06.09.658565

**Authors:** Ana Paredes-Vázquez, Eva Balsa-Canto, Julio R. Banga

**Affiliations:** Computational Biology Lab, MBG-CSIC (Spanish National Research Council), Pontevedra, Galicia, Spain; Biosystems and Bioprocess Engineering, IIM-CSIC (Spanish National Research Council), Vigo, Galicia, Spain; Applied Mathematics Dept., Universidade Santiago de Compostela, Santiago de Compostela, Galicia, Spain

## Abstract

Microbial communities, complex ecological networks crucial for human and planetary health, remain poorly understood in terms of the quantitative principles governing their composition, assembly, and function. Dynamic modeling using ordinary differential equations (ODEs) is a powerful framework for understanding and predicting microbiome behaviors. However, developing reliable ODE models is severely hampered by their nonlinear nature and the presence of significant challenges, particularly critical issues related to identifiability.

Here, we address the identification problem in dynamic microbial community models by proposing an integrated methodology to tackle key challenges. Focusing on nonlinear ODE-based models, we examine four critical pitfalls: identifiability issues (structural and practical), unstable dynamics (potentially leading to numerical blow-up), underfitting (convergence to suboptimal solutions), and overfitting (fitting noise rather than signal). These pitfalls yield unreliable parameter estimates, unrealistic model behavior, and poor generalization. Our study presents a comprehensive workflow incorporating structural and practical identifiability analysis, robust global optimization for calibration, stability checks, and rigorous predictive power assessment. The methodology’s effectiveness and versatility in mitigating these pitfalls are demonstrated through case studies of increasing complexity, paving the way for more reliable and mechanistically insightful models of microbial communities.

**Availability:** The code that implements the methodology and reproduces the results is available at https://doi.org/10.5281/zenodo.15309438

**Supplementary information:** Additional information supporting this manuscript is provided at https://doi.org/10.5281/zenodo.15309438.

**Author summary:** Microbial communities, vital for human and environmental health, are complex systems whose quantitative behaviors are not yet fully understood. Scientists employ mathematical models to study these communities, but developing accurate models of their dynamics is very challenging. Key difficulties include determining correct model parameters, ensuring model stability, and preventing the models from either under-learning from data or over-learning from noise. This paper presents a new, integrated methodology to overcome these obstacles. The approach provides a systematic workflow incorporating analyses of parameter identifiability, model stability, and predictive capabilities. By addressing these critical pitfalls, this research aims to facilitate the creation of more reliable and mechanistically insightful models, ultimately enhancing our understanding of microbial community dynamics.

## Introduction

A microbial community, often referred to as a microbiome, is a group of microorganisms that live together in a specific environment [1]. These microorganisms can include bacteria, archaea, fungi, and viruses. They interact with each other in complex ways, forming intricate ecosystems. These communities play crucial roles in various processes. For instance, the gut microbiota influences digestion and immunity, while the soil microbiome is essential for nutrient cycling and plant growth. Understanding microbial communities is vital for fields such as bio-medicine, microbiology, ecology, biotechnology, and agriculture, as they can impact human health and environmental sustainability [2].

Characterizing the nature of interactions within these systems helps reveal the roles of microbial species. Various qualitative and quantitative methods have been developed to analyze microbial community functions [3]. Quantitative approaches based on mathematical modeling are especially helpful. They provide valuable insights into the functioning of microbial communities, helping researchers to understand, predict, and potentially manipulate these complex systems [4–6]. The abstraction capabilities of these mathematical models are crucial for capturing underlying phenomena and linking the various scales at which these systems operate [7]. In this context, a number of network inference methods have been applied to characterize microbial interactions as static network models, sometimes mapping the inferred interactions with ecological motifs, such as cooperation, competition, commensalism, predation, and amensalism [8]. However, these network models offer a static view of microbial communities, capturing their status at a specific moment. In order to study changes over time, and time-dependent phenomena such as stability, response to disturbances, and succession, dynamic models are necessary.

Dynamic modeling is a particularly powerful framework that helps infer directionality and causality, predict time-dependent properties, and capture the dynamic behaviors of microbiomes in changing environments [9]. It can identify key microbes, molecules, and genetic determinants with significant causal effects on microbial community behaviors, predict their responses to perturbations, and guide the design of precise interventions to modify community functions [10]. Many existing frameworks for modeling the dynamics of microbial communities have been inspired by ecosystem modeling approaches, and include Lotka–Volterra, consumer–resource, trait-based, individual-based and genome-scale metabolic models, each one with its own scope, advantages and limitations [5, 11, 12]. In particular, Generalized Lotka-Volterra (GLV) and consumer-resource (CR) models are widely used [5, 13], although their suitability depends on the origin of the available experimental data and the theoretical assumptions [14].

While various dynamic modeling paradigms exist, including agent-based models (ABMs) and partial differential equations (PDEs) for spatially explicit scenarios [15, 16], ordinary differential equations (ODEs) are frequently used when spatial effects are averaged or not the primary focus [7, 17]. ODEs effectively capture time-dependent processes like the complex interactions between microbial species, such as competition, cooperation, and predation. Moreover, they can incorporate the influence of environmental factors (e.g., nutrient availability, temperature, and pH) on microbial growth and interactions, and can be used to make quantitative predictions about the behavior of microbial communities under different conditions. This can help understand the underlying mechanisms driving community dynamics and design time-dependent interventions to manipulate these communities. In the remainder, we consider dynamic modeling of microbial communities using ODE-based models. In any case, it should be noted that several of the methods described below can be extended to handle PDEs, and probably ABMs.

Building these ODE dynamic models is challenging due to several factors. To begin with, microbial communities involve complex networks of interactions among microbes and with their environment. These interactions can be direct (e.g., competition, predation), indirect (e.g., through metabolite exchange), or higher-order (beyond simple pairwise interactions). Capturing all these interactions accurately and mapping them to the structure of an ODE-based model is difficult [4, 5]. But, more importantly, even when the model structure is adequate, we still need to solve the identification problem, i.e., fitting the model to data and evaluating its predictive power.

The identification of dynamic nonlinear ODE-based models presents a complex set of challenges that stem from both the intrinsic properties of the models and the nature of available data. These challenges can significantly impact the accuracy, reliability, and predictive power of the resulting models. One of the primary challenges is the potential lack of identifiability. Given the model (as a set of ODEs) and the mathematical formulation of the measured quantities (e.g., microbial abundances, nutrient concentrations, etc.), lack of identifiability can manifest in two forms:

- Structural Identifiability: this arises from the structure of the model and its mapping with the measured variables. Some parameters or combinations of parameters may be inherently unidentifiable, regardless of the quality or quantity of data available.
- Practical Identifiability: even when a model is structurally identifiable, limitations in the available data (such as noise, sparsity, or limited range) can render certain parameters practically unidentifiable.

A second challenge is related to the nonlinear nature of these models, which introduces additional layers of complexity to the parameter estimation process, notably the non-convexity of the optimization problem used in the fitting process. Also, it should be noted that the solutions of some nonlinear ODEs may not exist for all integration times. A phenomenon known as finite-time blow-up, also called explosive instability, can occur, where solutions approach infinity in a finite amount of time, which is often biologically unrealistic but mathematically possible in some models. This behavior has been observed even in relatively simple ecological models [18, 19].

In this study, we address the identification problem within microbial communities and propose an integrated methodology to effectively tackle these challenges. Specifically, we examine four critical pitfalls that commonly undermine the identification of nonlinear ODE-based models:

- Identifiability issues, structural or practical: as mentioned above, can lead to unreliable parameter estimates or multiple equivalent solutions.
- Blow-up: if the model exhibits finite-time blow-up for certain parameter combinations, and the optimization algorithm explores these regions, this can lead to numerical issues that hinder optimization.
- Underfitting: this occurs when the estimation algorithm converges to a local optimum rather than the global optimum. It results in a model that fails to capture the full complexity of the underlying system dynamics.
- Overfitting: in this case, the model fits the noise in the data rather than the true underlying signal. Overfitted models often perform well on training data but fail to generalize to new scenarios, i.e., they lack predictive power.

These four issues are often intertwined and can exacerbate each other. For example, fitting a model with unidentifiable parameters might be more prone to overfitting and getting trapped in local optima. Blow-up dynamics can further complicate the optimization landscape and make it harder to find the true parameters.

We demonstrate that if these pitfalls are not properly detected and mitigated, they can lead to significant modeling artifacts. These artifacts may include biased or inconsistent parameter estimates, unrealistic model behavior in certain regimes, poor generalization to new data or scenarios, and misleading interpretations of system dynamics. Ultimately, models suffering from these issues lack robust predictive power and may lead to incorrect conclusions about the system being studied.

To the best of our knowledge, these issues have not been thoroughly examined in the microbial communities literature. Only recently have a few studies begun to explore identifiability analysis for some of these models. Two of these works address structural identifiability specifically [20, 21], while another considers both structural and practical identifiability [22]. Other critical challenges, however, remain largely unexamined.

Here, we present an integrated and systematic methodology to detect and address these issues simultaneously. This paper is structured as follows: first, we describe a comprehensive workflow to properly evaluate identifiability, perform robust model calibration, and assess the predictive power of the resulting fitted model. We also introduce a software implementation of this workflow that makes use of electronic notebooks, making it more accessible and user-friendly. Next, we apply our methodology to case studies of increasing complexity, including several examples of widely used canonical models, to demonstrate its effectiveness and versatility. Finally, we discuss the main findings, emphasizing the effectiveness of the proposed methodology in addressing the identified pitfalls.

## Materials and methods

We present a methodology for the proper identification and assessment of dynamic models of microbial communities described by sets of nonlinear Ordinary Differential Equations (ODEs).

Our integrated computational workflow consists of three sequential phases, each subdivided into modular subtasks:

- Phase 1, *pre-estimation preparation*, begins with structural identifiability analysis (SIA) to determine whether model parameters are theoretically distinguishable. If simplification is required, users can remove non-identifiable parameters, reparameterize equations to reduce complexity, or explore alternative input-output mappings by adjusting perturbed or measured variables.
- Phase 2, *robust parameter estimation*, employs global optimization techniques to avoid local minima, paired with efficient adaptive ODE solvers and mechanisms to address numerical instabilities and overfitting. After estimation, practical identifiability analysis (PIA) is used to assess whether model parameters can be reliably estimated from the available data, considering measurement noise and experimental constraints. When needed, it can be followed by dynamic stability analysis via Jacobian eigenvalue evaluations to assess system robustness.
- Phase 3, *predictive power analysis*, checks possible over-fitting and its consequences. It also evaluates predictive power and generalizability by comparing simulations against experimental data and additional cross-validation tests with held-out datasets. The results can help users to refine the model if needed. Refinement options include simplifying the model structure, redesigning experiments to improve data quality, or collecting additional data to resolve ambiguities.

This workflow, implemented via version-controlled scripts and interactive electronic notebooks, ensures reproducibility and transparency. By integrating theoretical rigor (e.g., identifiability checks, stability analysis) with computational robustness (e.g., global optimization, regularization), this pipeline minimizes risks such as over-fitting, non-identifiability, and unstable dynamics, yielding reliable models for real-world applications. The methodology is outlined in Figure 1. We describe the different phases in the following subsections, with further technical details in the Supporting Information file.

**Fig 1:**
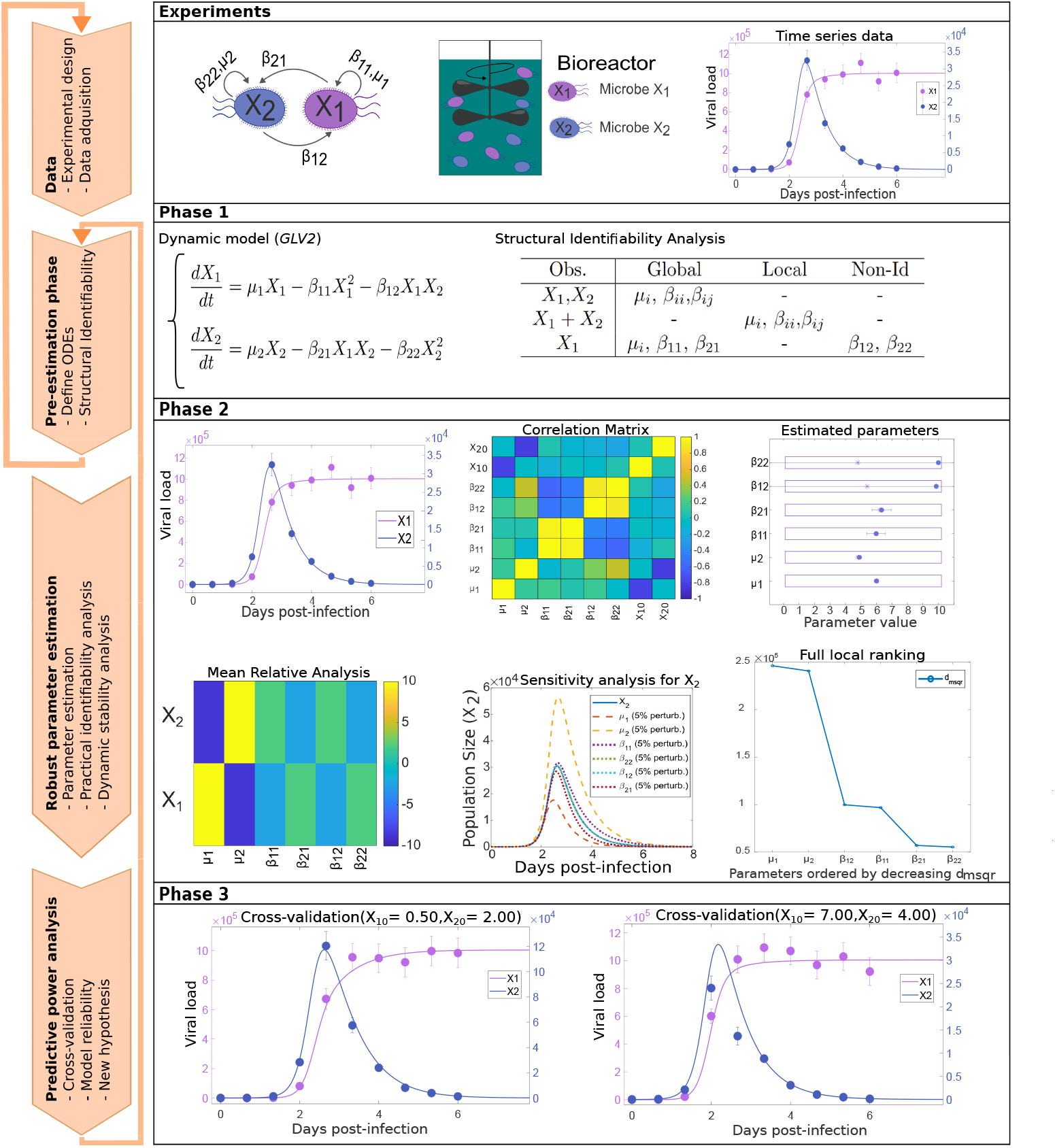
Diagram illustrating the workflow for model calibration. The figures correspond to a simple Generalized Lotka-Volterra model with 2 species in a competition scenario.

### Pre-estimation preparation

The microbial communities under consideration are described by sets of non-linear Ordinary Differential Equations (ODEs) and the observation function as follows:

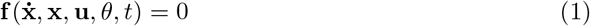

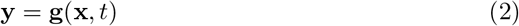

where 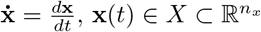 is the vector of state variables at time *t*, with initial conditions 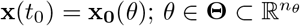 denotes the vector of model parameters within the feasible parameter space Θ, and 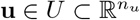 corresponds to the external factors or inputs. 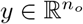 regards the vector of *n*_*o*_ observables. We will regard as fully observed (`FO`) systems those for which all the state variables are amenable to experimentation and partially observed (`PO`), otherwise.

Structural identifiability analysis (SIA) seeks to determine which model parameters can be uniquely estimated from a given perfect data set, i.e., continuous and noise-free. It should be noted that in reality we never have perfect, noise-free data and measurements over infinite time. The question is whether parameter estimation is even theoretically possible given the model structure in Eqns. (1)-(2). In general, we can distinguish:

- Global structural identifiability: parameters can be uniquely determined from the observations.
- Local structural identifiability: there is a finite set of parameter values that yield the same observations. In synthetic problems, parameters can be uniquely inferred near their nominal values, but multiple equivalent solutions may exist in the parameter space.
- Structural non-Identifiability: there is an infinite set of parameter values that yield the same observations.

Structural identifiability is crucial for meaningful parameter estimation, as non-identifiable models yield unreliable and non-unique parameter values that may not reflect the true system properties. Inaccurate estimates of mechanistically significant parameters compromise the usefulness of the model to provide biological insights [23–25]. It also impacts model validation and interpretation, making it difficult to draw robust conclusions when parameters cannot be uniquely determined. Furthermore, identifying non-identifiable parameters informs new experiment design, guiding data collection and experimental strategies to improve parameter identifiability.

Several methods exist for studying `SIA` in nonlinear models [23, 26], but they often rely on complex symbolic manipulations. While various software tools are available, they tend to be user-unfriendly and require significant expertise. Moreover, no single method applies universally to all models. These challenges create barriers that often lead to the neglect of this crucial step in model calibration.

After an initial screening based on our requirement to analyze structural global identifiability, as well as considerations of computational scalability, flexibility (including multi-experiment identifiability of both parameters and initial conditions), and previously reported performance [26], we selected the following tools for further evaluation:

- `GenSSI2` [27] is a MATLAB-based tool that uses a generating series approach. It transforms the model equations into a system of polynomial equations on the parameters, analyzing their rank conditions to determine local identifiability and solving the system of equations to analyze global identifiability. However, its reliance on pure symbolic computations can make it computationally demanding for partially observed, highly non-linear systems.
- `SIAN` [28, 29] is a Julia package that uses a randomized Monte-Carlo algorithm and an improved version of the Taylor series approach. It employs a differential algebra method that eliminates non-identifiable parameters through Gröbner basis computations.
- `Structural Identifiability` [30] is another Julia-based package that also uses differential algebra approach based on input-output equations. The method combines randomization and a differential elimination algorithm, and presents good scalability.

In principle, all these tools can handle both local and global identifiability, and can be applied to polynomial and rational ODE models. This selection offers a well-balanced combination of capabilities, performance, and suitability for the models under analysis. Each tool has unique strengths: `GenSSI2` is effective for smaller to medium-sized models but can struggle with scalability due to its symbolic computation overhead, while `SIAN` and `Structural Identifiability` employ symbolic-numeric randomized algorithms and leverage Julia’s modern computational advantages for improved scalability and efficiency.

### Robust parameter estimation

Dynamic model calibration is the process of adjusting the parameters to ensure accurate representation of the biological system’s behavior based on available data [31]. This process is typically framed as an optimization problem aimed at minimizing discrepancies between model predictions and observed data. Here we adopt a single-shooting approach, where the initial value problem described by the ODEs is solved for each evaluation of the cost function [32]. Other options are discussed elsewhere [33, 34].

The choice of objective function plays a crucial role in quantifying this mismatch, as it directly impacts the accuracy and reliability of the calibration. The objective function encapsulates characteristics of the measurement process and incorporates prior knowledge. In frequentist approaches, it typically takes the form of the maximum (log)-likelihood function (details in the Supporting Information file). When standard deviations are known, this function follows a weighted least-squares form:

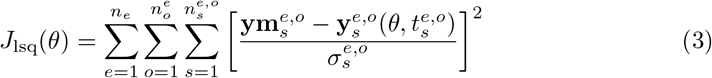

Where *n*_*e*_, 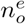 and 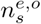 regard the number of experiments, the number of observables in a given experiment *e* and the number of sampling times for the specific observables and experiment, respectively. 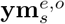 represents the measured data at a given sampling time *s* for a specific observable *o* in the experiment *e*; 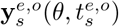 denotes the model prediction for the sampling time 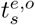; and 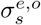 is the standard deviation of noise. The estimation problem is formulated as the minimization of one of these objective functions subject to differential and algebraic constraints (i.e., the ODEs describing the dynamics plus the observation function and bounds on the parameters, *θ*_min_ ≤ *θ* ≤ *θ*_max_). This optimization problem is non-convex, so in our workflow, we use global optimization solvers. In particular, we employed a metaheuristic approach known as *enhanced Scatter Search* (eSS), which combines global search heuristics with local optimization steps to refine potential solutions and is recognized for its reliable convergence properties [35].

### Stability analysis

In the above formulation, the estimation relies on repeatedly solving the ODE system for different parameter values within an optimization loop. If the ODE system is unstable (e.g., exhibits blow-up) for even a small region of parameter space, the simulations will become unreliable or impossible to complete. For certain parameter values, the numerical solver might fail, produce nonsensical results (NaN, Inf), or take excessively long to simulate. This disrupts the optimization process, making it difficult to evaluate the objective function and navigate the parameter space. Even when convergence to a good fit is achieved, it is important to check if the calibrated model is stable.

Here we use a strategy for stability analysis for nonlinear dynamical systems which begins by identifying equilibrium points and then describing the system’s behavior around these points. This is achieved by linearizing the system using the Jacobian matrix at each equilibrium point, which captures the local dynamics. The stability is then determined by analyzing the real parts of the eigenvalues of this Jacobian matrix. If all real parts are negative, the equilibrium is asymptotically stable; if at least one is positive, it is unstable; and if all are non-positive with some being zero, it is neutrally stable. Based on the patterns of these eigenvalues, equilibrium points can be further classified as stable or unstable nodes, saddle points, spirals, or centers. To understand the broader system behavior beyond local stability, numerical simulations are used to explore global dynamics, including periodic orbits or chaotic behavior. For complex systems, more advanced numerical methods can be employed to reveal aspects like basins of attraction, limit cycles, and chaotic regimes. This approach is applicable to a wide range of ODE systems and is described in more detail in the Supporting Information file.

### Practical Identifiability Analysis (PIA)

To assess the practical identifiability of parameters, our workflow uses the Fisher Information Matrix (FIM), computing parameter confidence intervals derived from the Cramèr-Rao inequality, and examining the correlation matrix. The process starts by constructing the FIM (eqn. 4), which quantifies the amount of information the data is expected to provide about the parameters:

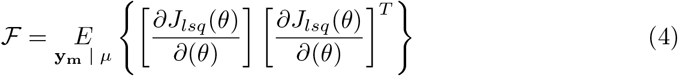

where *E* represents expected values and *µ* a value of the parameters, hopefully close to their real value. Computing the FIM involves calculating sensitivity matrices, which represent how parameter changes affect the model output, and incorporating assumptions about the measurement noise. The local parametric sensitivities for a specific experiment (*e*), observable (*o*), and sampling time (*t*_*s*_) read as follows:

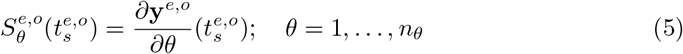

A key step is to evaluate the FIM’s properties: a singular or ill-conditioned FIM suggests practical non-identifiability, indicating that parameters or combinations of parameters cannot be reliably estimated from the data. The condition number and eigenvalues of the FIM can provide further insights into the degree of identifiability, with small eigenvalues or large condition numbers signaling potential issues.

Subsequently, confidence intervals for the parameters can be approximated using the Cramèr-Rao inequality, which states that the inverse of the FIM provides a lower bound on the variance of unbiased estimators:

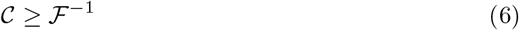

where 𝒞 is the covariance matrix. By taking the square root of the diagonal elements of the inverse FIM, one obtains estimates of the standard errors for each parameter, which are then used to construct confidence intervals. Wide confidence intervals indicate poor practical identifiability, meaning that even with the assumed data quality, the parameters cannot be estimated with high precision. These wide intervals suggest that different parameter values, spanning a considerable range, could plausibly explain the observed data, thus hindering accurate parameter determination.

Finally, the correlation matrix, derived from the inverse of the FIM, offers insights into the interdependencies between parameter estimates. Off-diagonal elements of this matrix reveal the correlation coefficients between pairs of parameters. High correlation values, close to 1 or -1, suggest that these parameters are not independently identifiable, i.e., changes in one parameter must be compensated by changes in the correlated parameter to maintain a similar model fit to the data. This correlation implies redundancy in the parameterization and can guide model simplification or experimental redesign to improve identifiability. By collectively analyzing the FIM’s condition, the Cramèr-Rao based confidence intervals, and the parameter correlation matrix, one can gain a comprehensive understanding of the practical identifiability of parameters in ODE models, highlighting parameters that are poorly identifiable and suggesting potential remedies such as model refinement or improved experimental design.

### Predictive power analysis

To assess the predictive power of a calibrated ODE model, a crucial next step is to investigate potential **overfitting**. This can be achieved by examining the residuals, which are the differences between the experimental data and the model predictions at the time points where data was collected. If the model is appropriately calibrated and not overfitting the training data, the residuals should exhibit a random distribution around zero, with no discernible patterns or trends. Visual inspection of residual plots, such as plotting residuals against time or predicted values, is helpful. Ideally, the residuals should resemble white noise, indicating that the model has captured the underlying signal and the remaining variation is attributable to random error rather than systematic model deficiencies.

Following residual analysis, a robust evaluation of predictive power requires **cross-validation**. This involves partitioning the available data into training and validation sets. The model is then calibrated using only the training data to obtain parameter estimates. Subsequently, the predictive capability of the calibrated model is assessed by comparing its predictions against the validation data, which was not used during calibration. If possible, this process should be repeated multiple times for several sets of validation data. The predictive accuracy can be quantified using metrics like the root mean squared error (RMSE) or normalized root mean squared error (NRMSE) between the model predictions and the validation data across all folds. Consistently good predictive performance on the validation sets, comparable to the fit on the training data, indicates robust predictive power and absence of overfitting.

If the model demonstrates good predictive power and no signs of overfitting, the final step involves evaluating the **mechanistic plausibility** of the model fit. This is crucial, especially when dealing with models built upon mechanistic principles. The estimated parameter values should be examined for their magnitudes and signs in the context of their mechanistic interpretation. For instance, parameters representing rates of biological processes should have positive values and magnitudes that are biologically reasonable. Similarly, the signs of parameters governing interactions should align with the expected mechanistic relationships (e.g., a parameter representing inhibition should have a negative effect). Deviations from expected ranges or signs for mechanistically meaningful parameters can suggest issues with the model structure, identifiability problems, or inconsistencies between the model and the underlying mechanisms, even if the model exhibits good statistical fit and predictive power. This mechanistic evaluation provides a crucial layer of validation beyond purely statistical measures, ensuring that the model not only fits the data but also offers a plausible and interpretable representation of the system under study.

### Software implementation

We developed our workflow as a unified, Matlab-based pipeline that integrates the previously mentioned three tools for structural identifiability analysis (SIA) alongside estimation and practical identifiability analysis (PIA) methods using `AMIGO2` [36]. `AMIGO2` (*Advanced Model Identification using Global Optimization*) is a powerful toolbox for dynamic modeling and optimization, offering a broad selection of nonlinear optimization solvers. These include direct and indirect local methods, multi-start local approaches, global stochastic algorithms, and hybrid optimization techniques. Additionally, `AMIGO2` facilitates PIA.

To enhance computational efficiency, particularly for demanding tasks such as parameter estimation, using `AMIGO2` in this framework enables automatic generation of C-compiled code, significantly improving performance. Furthermore, we extended the pipeline with additional code to support other key steps, such as stability analysis. Several case studies, based on commonly used models of microbial communities, were implemented and tested within this workflow.

The resulting integrated software is available in both electronic notebook format (live scripts) and as standard MATLAB scripts. It features the following key components:

- **Pre-estimation analysis:** the workflow simplifies model definition, data integration, and identifiability analysis, ensuring a solid foundation for parameter estimation. This module efficiently handles multi-experiment datasets, incorporates data visualization tools, and facilitates structural identifiability analysis using several methods, so non-expert users can determine whether model parameters can be uniquely inferred.
- **Robust parameter estimation:** a diverse set of global and local optimization algorithms enhances the reliability and accuracy of parameter estimation. Sensitivity analysis, practical identifiability assessment, and stability analysis help refine parameter estimates. The framework also supports easy cross-validation with additional datasets and leverages parallel computing for improved efficiency and scalability.
- **Predictive power analysis:** to ensure the model produces reliable and meaningful predictions, the workflow includes sensitivity analysis, statistical goodness-of-fit tests and cross-validation to assess model performance. These methods enable model comparison, interactive plotting, time series visualization, and statistical summarization, aiding in result interpretation. Furthermore, automated report generation and documentation via electronic notebooks streamline reproducibility and facilitate effective communication of findings.

## Results

We provide here a summary of the key findings, with further details available in the Supporting Information. We evaluated our integrated computational workflow using a set of canonical models representing common frameworks for modeling microbial communities (Table 1). These models range from Generalized Lotka-Volterra (GLV) systems of varying complexity to more intricate models involving resource competition, phage dynamics, and synthetic gene circuits. To rigorously test the workflow and assess its robustness against experimental variability, we generated synthetic pseudo-experimental datasets for each model, incorporating different types and levels of Gaussian noise. This use of synthetic data provides a ground truth for evaluating parameter recovery and model predictive capabilities.

**Table 1:**
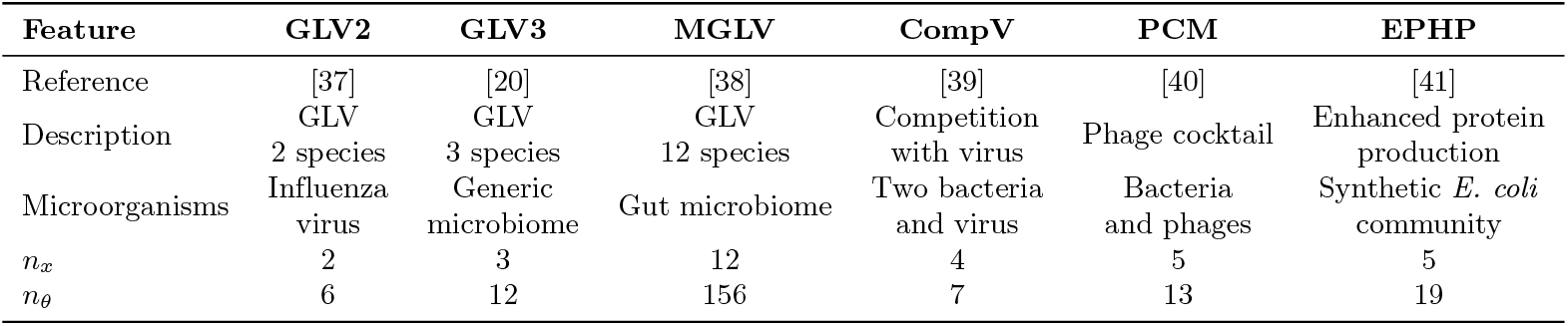
Overview of case studies. The first row shows abbreviated model names. *n*_*x*_ denotes the number of state variables, and *n*_*θ*_ the number of parameters. GLV denotes generalized Lotka-Volterra. For the **GLV2** model, two subcases were analyzed: competition and coexistence. Further details are provided in the Supporting Information.

### Phase 1: Structural Identifiability Analysis

We initiated the workflow with structural identifiability analysis (SIA) using `GenSSI2`, `SIAN`, and `Structural Identifiability`. The analysis covered scenarios ranging from ideal (fully observed states, known initial conditions) to realistic (partially observed states, unknown initial conditions). As summarized in Table 2, all models were found to be structurally identifiable under full observation. However, limitations arose under partial observation; for instance, in the GLV2 case, measuring only total biomass or a single species abundance resulted in non-identifiability, consistent with [22] and highlighting how experimental constraints impact parameter determination.

**Table 2:** Summary of SIA results. FO: Fully Observed, PO: Partially Observed, GI: Globally Identifiable, NGI: Non-Globally Identifiable (i.e. only a subset of parameters are identifiable). Last column indicates which methods were successful. Key: 1 = `GenSSI2`, 2 = `SIAN`, 3 = `Structural Identifiability`. Full details, including the partially observed schemes considered, can be found in the Supporting Information.

| Model | Measurements | SIA Result | Method |
| --- | --- | --- | --- |
| GLV2 | FO | GI | 1, 2, 3 |
|  | PO | NGI | 2, 3 |
| GLV3 | FO | GI | 1, 2, 3 |
|  | PO | NGI | 2, 3 |
| MGLV | FO | GI | 3 |
| CompV | FO | GI | 2, 3 |
| PCM | FO and PO | GI | 2, 3 |

The performance of SIA tools varied. `GenSSI2`’s symbolic approach was effective for simpler models but struggled with larger partially observed ones due to computational demands. `SIAN` and `Structural Identifiability`, combining symbolic and numerical techniques, handled more complex models efficiently. The EPHP model could not be analyzed due to its discontinuous formulation, a limitation of current SIA tools assuming continuous dynamics.

### Phase 2: Parameter Estimation and Practical Identifiability

This phase addresses the challenges of finding reliable parameter estimates from noisy data, considering potential non-convexity and numerical issues.

### Optimization Strategies and Performance

We first explored multistart local optimization using least-squares optimizers (`lsqnonlin, nl2sol`) [36]. This revealed significant challenges across most models, particularly non-convexity (multiple local optima) and model instability (numerical blow-ups). For example, in the GLV3 case with 10% noise (Figure 2), only 15% of runs converged, and only 6% reached objective function values near the nominal (true) value. Many runs terminated due to blow-ups or converged to poor local optima (LO) or potentially overfitting solutions (OF).

**Fig 2:**
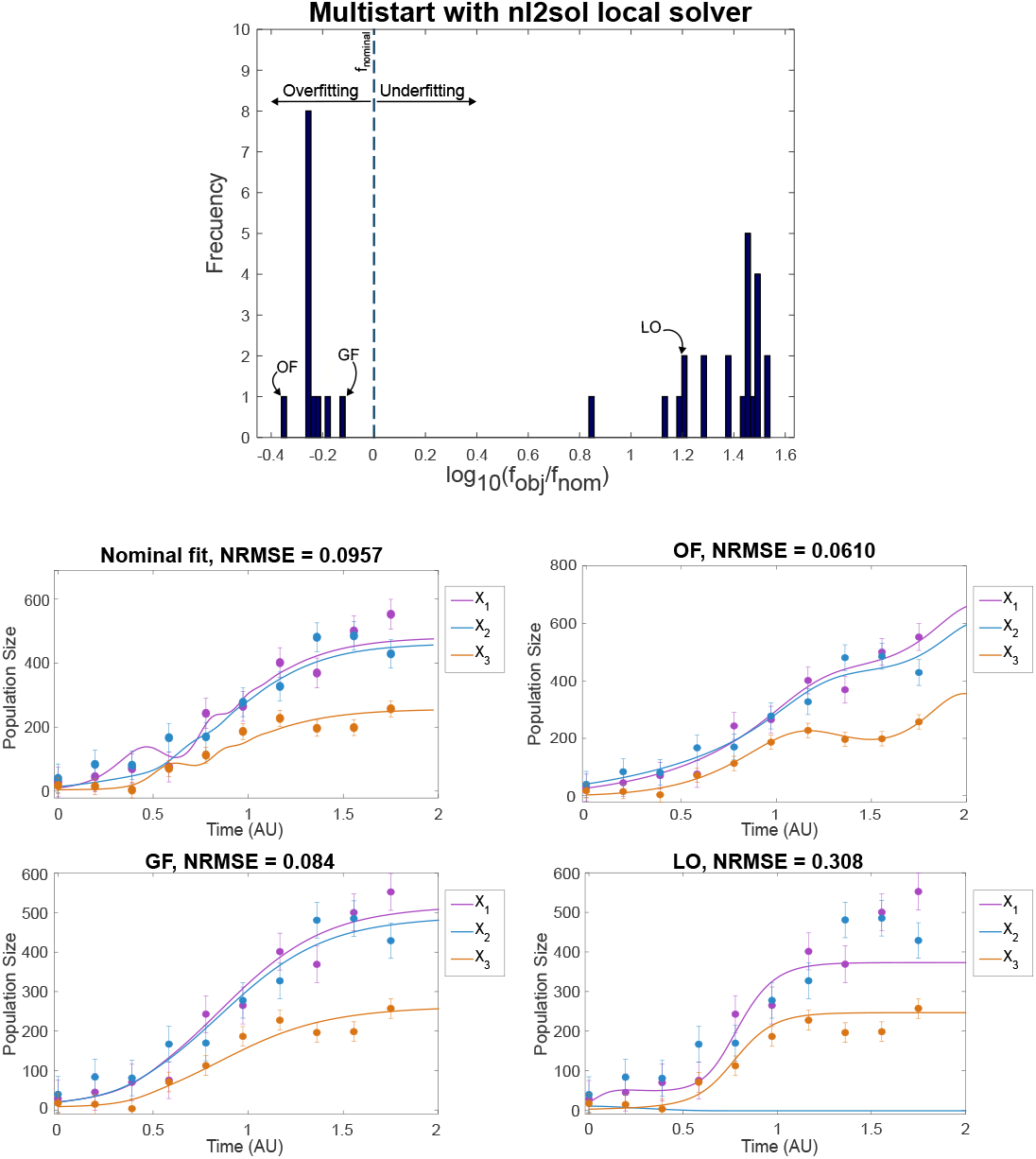
Case study ***GLV3*** : multistart optimization using the `nl2sol` local solver. Top figure shows a histogram of the 35 optima obtained in 200 runs, highlighting the presence of overfitting (OF), underfitting (local optimum, LO), and good fit (GF) solutions. The *x*-axis shows the log_10_ of the ratio between the objective function achieved and that of the nominal vector of parameters. Figures below present the nominal system behavior and show examples for the OF, GF, and LO cases, respectively.

Overall, multistart local methods showed acceptable performance only for the simplest cases (GLV2) and moderate effectiveness for PCM, achieving 70% good fits. For more complex cases, such as GLV3 and particularly MGLV models, convergence rates were significantly lower. Blow-ups occurred frequently (over 80%, even with tight bounds), and there was a high risk of converging to suboptimal solutions or overfitting. The high-dimensional MGLV model was especially challenging (good fits in only 12% or runs), often failing to accurately reproduce system dynamics, even when using noise-free data. In the CompV and EPHP cases, while most of the multistarts converged, good fits were scarce: only 13% of runs for EPHP and 30% for PCM achieved satisfactory results.

In contrast, the enhanced Scatter Search (`eSS`) global optimization algorithm exhibited significantly greater robustness. Convergence rates were consistently high (exceeding 95% overall and 85% for MGLV), effectively navigating complex optimization landscapes. While `eSS` did not completely eliminate challenges such as potential overfitting or local optima traps in highly complex models (e.g., MGLV, GLV3), it consistently delivered much more reliable results than multistart local methods. Additionally, the computational cost using `eSS` was very reasonable, ranging between 1 and 2 minutes for all cases on a standard PC with Intel i5-13500 CPU (further details available in the Supporting Information).

### Practical Identifiability and Mechanistic Interpretation

After optimization with `eSS`, we evaluated practical identifiability using the FIM to derive parameter correlations and confidence intervals (CIs), alongside sensitivity analysis. Results indicated a consistent decline in identifiability with increasing model complexity and noise. For simpler models like GLV2, parameters were reasonably well-defined with narrow CIs under full observation, although some correlations emerged (e.g., between *β*_12_ and *β*_22_, and between growth rates *µ*_*i*_ and initial conditions). Sensitivity analysis confirmed that parameters with larger CIs had less influence on system states (see Figure F in the Supporting Information).

However, for more complex models (GLV3, CompV, PCM, MGLV), PIA revealed significant limitations. Even when structurally identifiable, parameters often had wide CIs and strong correlations, especially with noisy data. This indicates that, despite achieving a good fit, the available data may not be adequate for accurately estimating all parameters. In other words, the insufficient information content of the data rendered it inadequate for reliable model calibration.

Crucially, poor practical identifiability can compromise the mechanistic interpretation of the model. This was evident in GLV models where estimated interaction coefficients (*β*_*ij*_) frequently had signs opposite to the ground truth values, even for models achieving a good fit (low RMSE/NRMSE). Such sign errors imply a misrepresentation of the underlying biological interactions (e.g., predicting competition instead of facilitation). Figure 4 illustrates this for the MGLV model, showing sign disagreements increasing with noise but present even in the noise-free case. This highlights inherent limitations in extracting mechanistic details when practical identifiability is poor, often linked to insufficient information content in the experimental data. Stability analysis was also performed on the best parameter sets to ensure they did not lead to unstable dynamics under the calibration conditions.

### Phase 3: Predictive Power Analysis

The final phase assessed model generalizability and overfitting risk using cross-validation with held-out data, also incorporating datasets from novel experimental conditions. Our analysis confirmed that overfitting is a significant risk across all models. Solutions achieving excellent fits to the training data often failed to predict system behavior accurately under new conditions. The GLV3 case study provides a clear example (Figure 3): an overfitted solution (OF) matches the training data well but exhibits unstable dynamics long-term (left panel) and performs poorly in cross-validation with different initial conditions (center panel). In contrast, a better-calibrated, though not perfectly identifiable, solution (GF) captures the trends under new conditions more reliably (right panel).

**Fig 3:**
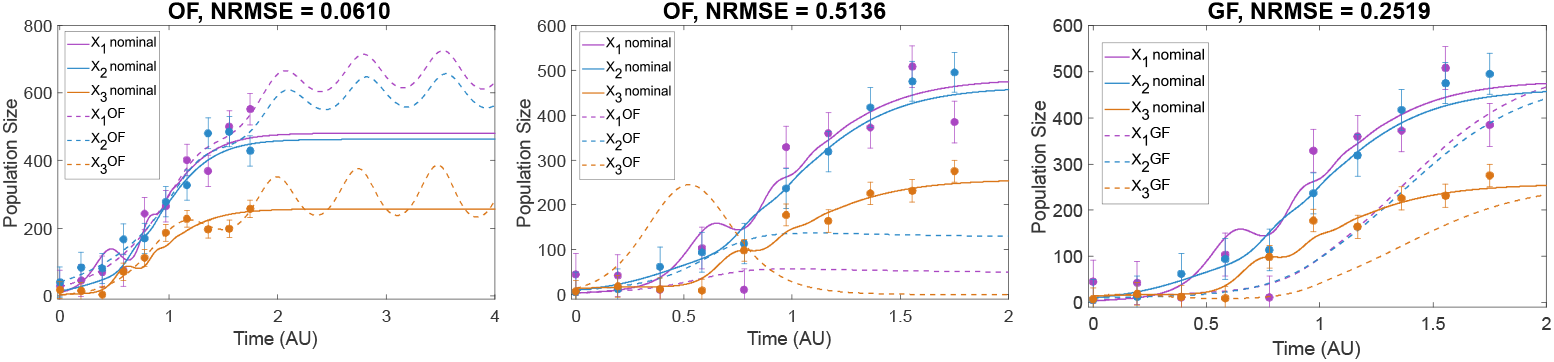
Case study ***GLV3***. Left figure: an overfitted solution OF that agrees very well with the data but shows oscillatory behavior (and eventually blow-up) after *t* = 2.0. Center figure: cross-validation of the same OF for different initial conditions, showing very poor predictive value. Right figure: cross-validation of a good fit (GF), showing that due to practical identifiability issues, the agreement with the data is not very good, although ultimately predicts well steady state values at final time.

**Fig 4:**
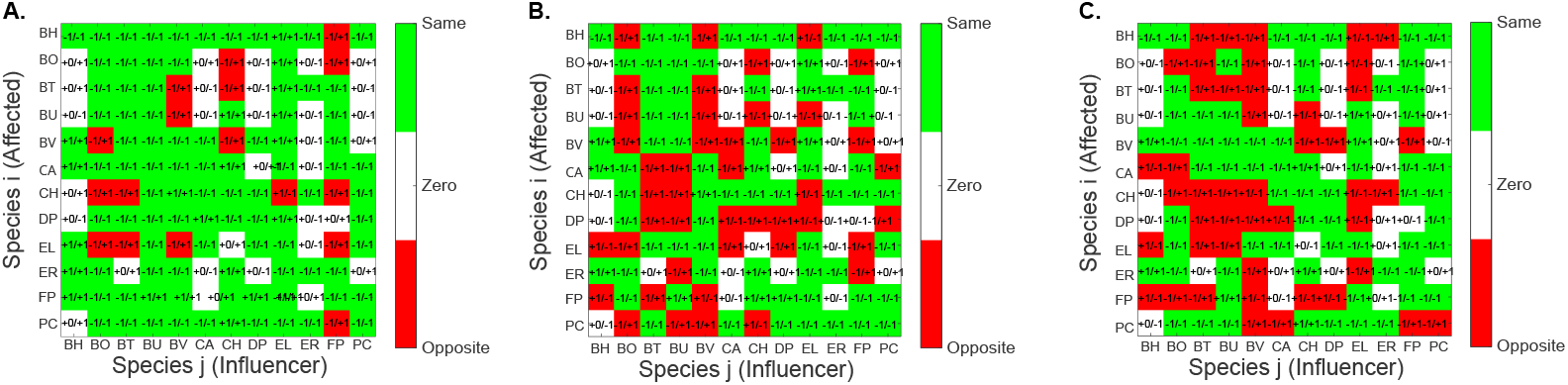
Case study ***MGLV*** : Sign agreement between estimated and nominal values of the interaction coefficient *β*_*ii*_. The figures correspond to successful calibrations to data with 0, 5 and 10% noise. Details provided in the Supporting Information.

This underscores the insufficiency of relying solely on goodness-of-fit to training data and highlights the necessity of cross-validation for evaluating true predictive power. Predictive capability generally decreased with increasing model complexity and noise. While simpler models could maintain some predictive power if carefully calibrated, complex models like MGLV showed severely compromised prediction, even when calibrated on noise-free data, linking back to the practical identifiability limitations discussed in Phase 2.

Recognizing that obtaining additional experimental data for cross-validation can be challenging, we also explored several techniques for residual analysis and concluded that Quantile-Quantile (Q-Q) plots can be used as a potential indicator of overfitting based solely on training data (details in Supporting Information).

Overall, this phase emphasizes the critical need for rigorous validation beyond the initial calibration dataset to ensure models are not merely fitting noise but capture the underlying system dynamics robustly, especially for complex biological systems like microbial communities.

## Discussion

Modeling the complex dynamics of microbial communities using nonlinear ODEs presents significant challenges, often hindering the development of reliable and predictive models. In this study, we proposed and evaluated an integrated computational workflow designed to systematically address four critical pitfalls inherent in the identification process: structural and practical identifiability issues, numerical instabilities leading to finite-time blow-up during estimation, convergence to suboptimal solutions (underfitting), and fitting experimental noise rather than the underlying signal (overfitting). Our results across a range of case studies, from simple GLV models to complex representations of gut microbiomes and synthetic systems (Table 1), demonstrate the prevalence of these issues and the necessity of a structured approach to mitigate them.

Identifiability is a cornerstone of reliable parameter estimation. Our workflow begins with structural identifiability analysis (SIA) in Phase 1, which, as shown in our results (Table 2), effectively flags theoretical limitations based on model structure and observation schemes before estimation commences. This step can guide experimental design by highlighting necessary measurements (e.g., demonstrating insufficiency of total biomass measurement in GLV2). However, even structurally identifiable models often suffer from practical non-identifiability, especially with increasing complexity and noise, as revealed by our practical identifiability analysis (PIA) in Phase 2. This was manifested through wide confidence intervals, parameter correlations, and, critically, incorrect sign estimations for interaction parameters (e.g., Figure 4), leading to mechanistically unsound models despite potentially good data fits. Addressing both SIA and PIA is crucial for obtaining meaningful parameter estimates.

The parameter estimation phase itself is fraught with difficulties arising from non-convex optimization landscapes exacerbated by potential model instabilities. Our findings highlight the limitations of standard multistart local optimization methods, which frequently failed due to encountering finite-time blow-up regions or converged to poor local optima (underfitting), as exemplified by the GLV3 results (Figure 2). The use of robust global optimization techniques, such as the enhanced Scatter Search (eSS) employed in Phase 2, proved significantly more effective in navigating these complex landscapes, reducing the incidence of underfitting and managing numerical integration failures caused by blow-ups, thereby increasing the likelihood of finding globally competitive solutions, even for challenging high-dimensional models like MGLV.

Achieving a good fit to the training data is insufficient proof of a model’s validity. Overfitting, where the model captures noise, poses a major threat to predictive power. Our results clearly demonstrated instances where models fit the training data well but failed dramatically when tested on unseen data via cross-validation (Phase 3, Figure 3). This underscores the indispensable role of cross-validation in assessing generalization capability. While cross-validation using additional datasets is preferred, residual analysis (e.g., Q-Q plots, see Supporting Information) offers a potential alternative check when obtaining such data is prohibitive. Detecting and avoiding overfitting is paramount for developing models that provide reliable predictions beyond the calibration conditions.

Crucially, our results demonstrate that the four pitfalls (identifiability, blow-up, underfitting, and overfitting) are often interconnected and can exacerbate one another. For instance, poor identifiability increases the risk of overfitting or converging to local minima. Blow-up dynamics can completely derail the search for meaningful parameters. Failing to address these intertwined problems systematically leads to significant modeling artifacts, as seen across our case studies. For example, in GLV models such artifacts manifest as unreliable parameters, often with incorrect signs that misrepresent biological mechanisms. Consequently, these models predict poorly under new conditions and can lead to flawed biological interpretations. Therefore, considerable caution and rigorous validation, extending beyond simple goodness-of-fit checks, are essential when developing and applying dynamic models. The integrated, multi-phase workflow presented here provides a structured methodology. It allows researchers to diagnose and mitigate these common pitfalls sequentially. This approach fosters the development of more robust, mechanistically plausible, and predictively powerful models, vital for advancing our understanding of microbial communities.

## Supporting information

S1 Text. Additional information supplementing this manuscript is provided at https://doi.org/10.5281/zenodo.15309438.

## Acknowledgments

JRB acknowledges support from grant PID2020-117271RB-C22 (BIODYNAMICS) funded by MCIN/AEI/10.13039/501100011033, from grant PID2023-146275NB-C22 (DYNAMO-bio) funded by MICIU/AEI/ 10.13039/501100011033 and ERDF/EU, and from grant CSIC PIE 202470E108 (LARGO). APV acknowledges support from grant PREP2023-002147, funded by MICIU/AEI/10.13039/501100011033/ and the European Social Fund Plus (ESF+), as part of the project PID2023-146275NB-C22.

The funding bodies played no role in the design of the study, the collection and analysis of data, or in the writing of the manuscript.

## References

1. Berg G, Rybakova D, Fischer D, Cernava T, Vergés MCC, Charles T, et al. Microbiome definition re-visited: old concepts and new challenges. Microbiome. 2020;8(1):103.

2. Eren AM, Banfield JF. Modern microbiology: Embracing complexity through integration across scales. Cell. 2024;187(19):5151–5170.

3. Srinivasan S, Jnana A, Murali TS. Modeling microbial community networks: Methods and tools for studying microbial interactions. Microb Ecol. 2024;87(1):56.

4. Widder Sea. Challenges in microbial ecology: building predictive understanding of community function and dynamics. ISME J. 2016;10(11):2557–2568.

5. van den Berg NI, Machado D, Santos S, Rocha I, Chacón J, Harcombe W, et al. Ecological modelling approaches for predicting emergent properties in microbial communities. Nat Ecol Evol. 2022;6(7):855–865.

6. Lange E, Kranert L, Krüger J, Benndorf D, Heyer R. Microbiome modeling: a beginner’s guide. Frontiers in microbiology. 2024;15:1368377.

7. Succurro A, Ebenhöh O. Review and perspective on mathematical modeling of microbial ecosystems. Biochem Soc Trans. 2018;46(2):403–412.

8. Faust K, Raes J. Microbial interactions: from networks to models. Nat Rev Microbiol. 2012;10(8):538–550.

9. Song HS, Cannon W, Beliaev A, Konopka A. Mathematical modeling of microbial community dynamics: A methodological review. Processes (Basel). 2014;2(4):711–752.

10. Qian Y, Lan F, Venturelli OS. Towards a deeper understanding of microbial communities: integrating experimental data with dynamic models. Curr Opin Microbiol. 2021;62:84–92.

11. Wade MJ, Oakley J, Harbisher S, Parker NG, Dolfing J. MI-Sim: A MATLAB package for the numerical analysis of microbial ecological interactions. PLOS ONE. 2017;12(3):e0173249. doi:10.1371/journal.pone.0173249.

12. Gao Y, Şimşek Y, Gheysen E, Borman T, Li Y, Lahti L, et al. miaSim: an R/Bioconductor package to easily simulate microbial community dynamics. Methods in Ecology and Evolution. 2023;14(8):1967–1980. doi:10.1111/2041-210x.14129.

13. Gonze D, Coyte KZ, Lahti L, Faust K. Microbial communities as dynamical systems. Curr Opin Microbiol. 2018;44:41–49.

14. Picot A, Shibasaki S, Meacock OJ, Mitri S. Microbial interactions in theory and practice: when are measurements compatible with models? Curr Opin Microbiol. 2023;75(102354):102354.

15. Lardon LA, Merkey BV, Martins S, Dötsch A, Picioreanu C, Kreft JU, et al. iDynoMiCS: next-generation individual-based modelling of biofilms. Environ Microbiol. 2011;13(9):2416–2434.

16. Zomorrodi AR, Segré D. Synthetic ecology of microbes: Mathematical models and applications. J Mol Biol. 2016;428(5):837–861.

17. Yip A, Smith-Roberge J, Khorasani SH, Aucoin MG, Ingalls BP. Calibrating spatiotemporal models of microbial communities to microscopy data: A review. PLoS Comput Biol. 2022;18(10):e1010533.

18. Parshad RD, Upadhyay RK, Mishra S, Tiwari SK, Sharma S. On the explosive instability in a three-species food chain model with modified Holling type IV functional response. Mathematical Methods in the Applied Sciences. 2017;40(16):5707–5726. doi:10.1002/mma.4419.

19. Batabyal S, Jana D, Lyu J, Parshad RD. Explosive predator and mutualistic preys: A comparative study. Physica A: Statistical Mechanics and its Applications. 2020;541:123348. doi:10.1016/j.physa.2019.123348.

20. Remien CH, Eckwright MJ, Ridenhour BJ. Structural identifiability of the generalized Lotka–Volterra model for microbiome studies. Royal Society Open Science. 2021;8(7):201378.

21. Díaz-Seoane S, Sellán E, Villaverde AF. Structural identifiability and observability of microbial community models. Bioengineering (Basel). 2023;10(4).

22. Balsa-Canto E, Alonso-Del-Real J, Querol A. Mixed growth curve data do not suffice to fully characterize the dynamics of mixed cultures. Proc Natl Acad Sci U S A. 2020;117(2):811–813.

23. Chis OT, Banga JR, Balsa-Canto E. Structural identifiability of systems biology models: a critical comparison of methods. PloS one. 2011;6(11):e27755.

24. Wieland FG, Hauber AL, Rosenblatt M, Tönsing C, Timmer J. On structural and practical identifiability. Current Opinion in Systems Biology. 2021;25:60–69. doi:10.1016/j.coisb.2021.03.005.

25. Villaverde AF. Observability and structural identifiability of nonlinear biological systems. Complexity. 2019;2019(1):8497093.

26. Rey Barreiro X, Villaverde AF. Benchmarking tools for a priori identifiability analysis. Bioinformatics. 2023;39(2):btad065.

27. Ligon TS, Fröhlich F, Chiş OT, Banga JR, Balsa-Canto E, Hasenauer J. GenSSI 2.0: multi-experiment structural identifiability analysis of SBML models. Bioinformatics. 2018;34(8):1421–1423.

28. Hong H, Ovchinnikov A, Pogudin G, Yap C. SIAN: software for structural identifiability analysis of ODE models. Bioinformatics. 2019;35(16):2873–2874. doi:10.1093/bioinformatics/bty1069.

29. Hong H, Ovchinnikov A, Pogudin G, Yap C. Global identifiability of differential models. Communications on Pure and Applied Mathematics. 2020;73(9):1831–1879.

30. Dong R, Goodbrake C, Harrington HA, Pogudin G. Differential elimination for dynamical models via projections with applications to structural identifiability. SIAM Journal on Applied Algebra and Geometry. 2023;7(1):194–235.

31. Villaverde AF, Pathirana D, Fröhlich F, Hasenauer J, Banga JR. A protocol for dynamic model calibration. Briefings in bioinformatics. 2022;23(1):bbab387.

32. Schittkowski K. Numerical data fitting in dynamical systems: a practical introduction with applications and software. vol. 77. Springer Science & Business Media; 2002.

33. Chung M, Krueger J, Pop M. Identification of microbiota dynamics using robust parameter estimation methods. Mathematical Biosciences. 2017;294:71–84. doi:10.1016/j.mbs.2017.09.009.

34. Shin S, Venturelli OS, Zavala VM. Scalable nonlinear programming framework for parameter estimation in dynamic biological system models. PLoS computational biology. 2019;15(3):e1006828.

35. Egea JA, Balsa-Canto E, García MSG, Banga JR. Dynamic Optimization of Nonlinear Processes with an Enhanced Scatter Search Method. Industrial & Engineering Chemistry Research. 2009;48(9):4388–4401. doi:10.1021/ie801717t.

36. Balsa-Canto E, Henriques D, Gábor A, Banga JR. AMIGO2, a toolbox for dynamic modeling, optimization and control in systems biology. Bioinformatics. 2016;32(21):3357–3359. doi:10.1093/bioinformatics/btw411.

37. Dimas Martins A, Gjini E. Modeling Competitive Mixtures With the Lotka-Volterra Framework for More Complex Fitness Assessment Between Strains. Frontiers in Microbiology. 2020;11. doi:10.3389/fmicb.2020.572487.

38. Venturelli OS, Carr AV, Fisher G, Hsu RH, Lau R, Bowen BP, et al. Deciphering microbial interactions in synthetic human gut microbiome communities. Molecular systems biology. 2018;14(6):e8157.

39. Alsolami AA, El Hajji M. Mathematical Analysis of a Bacterial Competition in a Continuous Reactor in the Presence of a Virus. Mathematics. 2023;11(4). doi:10.3390/math11040883.

40. Li G, Leung CY, Wardi Y, Debarbieux L, Weitz JS. Optimizing the Timing and Composition of Therapeutic Phage Cocktails: A Control-Theoretic Approach. Bulletin of Mathematical Biology. 2020;82.

41. Mauri M, Gouzé JL, de Jong H, Cinquemani E. Enhanced production of heterologous proteins by a synthetic microbial community: Conditions and trade-offs. PLOS Computational Biology. 2020;16(4):1–30. doi:10.1371/journal.pcbi.1007795.

