## SUPPORTING INFORMATION for "Identification of dynamic models of microbial communities"

### SUPPORTING INFORMATION OF THE PAPER
### Identification of dynamic models of microbial communities

Ana Paredes-Vázquez[1,3], Eva Balsa-Canto[2] and Julio R. Banga[1]
[1] Computational Biology Lab, MBG-CSIC (Spanish National Research Council), Pontevedra, Galicia, Spain,
[2] Biosystems and Bioprocess Engineering, IIM-CSIC (Spanish National Research Council), Vigo, Galicia, Spain,
[3] Applied Mathematics Dept., Universidade Santiago de Compostela, Santiago de Compostela, Galicia, Spain

12th June 2025

### 1 Overview

This supplementary information document provides detailed methodological descriptions and comprehensive results, supporting the findings presented in the main manuscript. Our primary focus is the challenge of identifying reliable dynamic models, typically based on Ordinary Differential Equations (ODEs), for complex microbial communities.

Here, we describe the integrated and systematic computational workflow proposed in the main text. This workflow sequentially addresses critical aspects of model development. This document presents the mathematical formulation of the problems addressed, elaborates on the specific algorithms and metrics employed within each phase of the workflow, and provides practical guidance on software installation for reproducibility.

Furthermore, we provide exhaustive results from applying this workflow to a diverse set of case studies, ranging from simple Generalized Lotka-Volterra (GLV) systems to more complex microbial interaction models (e.g., competition with virus, phage-cocktail dynamics, consumer-resource models). For each case study, we present detailed structural identifiability analysis results comparing different tools, comprehensive practical identifiability analysis results under varying noise levels (including parameter estimates, error metrics, model fits, convergence plots, correlation matrices, and sensitivity analyzes), and specific examples illustrating the common pitfalls encountered: structural and practical identifiability limitations (particularly with increasing complexity and noise), numerical instabilities, convergence to local optima (underfitting), and the dangers of overfitting leading to poor predictive power.

Collectively, this supplementary information offers a thorough technical resource and extensive empirical evidence demonstrating the challenges in microbial community modeling and the utility of our proposed workflow in navigating these complexities to build more reliable and predictive dynamic models.

### 2 Problem definition

We assume that the microbial community is described by a vector of state variables $\mathbf{x}(t) \in \boldsymbol{X} \subset \mathbb{R}^{n_x}$, with initial conditions $\mathbf{x}_0(\boldsymbol{\theta})$. This vector is the unique solution of the following nonlinear ordinary differential equations:

$$\mathbf{f}(\dot{\mathbf{x}}, \mathbf{x}, \mathbf{u}, \boldsymbol{\theta}, t) = 0 \tag{1}$$

where $\dot{\mathbf{x}} = \frac{d\mathbf{x}}{dt}, \mathbf{u} \in \boldsymbol{U} \subset \mathbb{R}^{n_u}$ corresponds to the external factor, $\boldsymbol{\theta} \in \boldsymbol{\Theta} \subset \mathbb{R}^{n_\theta}$ is the vector of model parameters, where $\boldsymbol{\Theta}$ is the feasible parameter space. Given an experimental scheme with $n_e$ experiments, $n_o^e$ observables per experiment $e$, and $n_s^{e,o}$ sampling times per experiment $e$ and observable $o$, $\mathbf{y}^{e,o} \in \mathbf{Y} \subset \mathbb{R}^{n_s^{e,o}}$ will regard the vector of $n_s^{e,o}$ discrete time measurements as follows:

$$\mathbf{y}^{e,o}(t_s^{e,o}, \mathbf{u}, \boldsymbol{\theta}) = \mathbf{g}^{e,o}(\mathbf{x}(\mathbf{u}, \boldsymbol{\theta}, t_s^{e,o}), \boldsymbol{\theta}, t) \tag{2}$$

$$s = 1, ..., n_s^{e,o}$$

where $t_s^{e,o}$ regards the $s^{th}$ sampling time for observable $o$ in experiment $e$; $\mathbf{ym}_s^{e,o}$ represents the measured data; $\mathbf{y}_s^{e,o}(\boldsymbol{\theta})$ denotes the simulation result for the sampling time $t_s^{e,o}$.

### 3 Methodology and workflow

The subsequent sections outline a detailed methodology for achieving effective model calibration. Dynamic model calibration, also known as parameter estimation, is the process of adjusting model parameters to ensure that the model accurately represents the behavior of the biological system under study (i.e., the model represents well the available data). This process is fundamental in systems biology, as it ensures that models are reliable, predictive, and capable of providing meaningful insights into complex biological phenomena. By bridging the gap between theoretical frameworks and real-world systems, model calibration empowers researchers to draw actionable conclusions based on model predictions.

Nonlinear dynamic model identification is often a challenging task due to the frequent lack of identifiability, which can be categorized as practical or, in the worst case, structural. Several authors have reported difficulties in determining unique and meaningful parameters from given experimental datasets, as broad parameter ranges can result in similar model predictions (Balsa-Canto *et al.*, 2010).

In order to surmount these challenges, here we propose and utilize an integrated computational workflow. This workflow is designed to sequentially tackle critical issues, such as identifiability, numerical stability, underfitting, and overfitting, thereby enhancing the reliability and predictive validity of the resulting models. It is structured into three distinct phases, each comprising specific modular subtasks aimed at ensuring rigorous model development and validation.

Our integrated computational workflow consists of the following three sequential phases:

- **Phase 1: Pre-estimation Preparation**
  This initial phase focuses on preparing the model structure and assessing its theoretical suitability before attempting parameter estimation from data. Key tasks include:
  - *Structural Identifiability Analysis* (SIA): Determining whether the model parameters are theoretically distinguishable given the model structure and the planned experimental outputs (measured variables). This analysis identifies parameters or parameter combinations that cannot be uniquely determined, even with perfect, noise-free data.
  - *Model Refinement (if needed):* Based on SIA results, if structural non-identifiability is detected, this task involves modifying the model or experimental plan. Options include simplifying the model by removing non-identifiable parameters, reparameterizing equations to reduce complexity, or exploring alternative input-output mappings by adjusting the experimental design (e.g., changing measured variables or perturbations).
- **Phase 2: Robust Parameter Estimation**
  This phase involves fitting the model parameters to experimental data while addressing numerical challenges and assessing the practical reliability of the estimates. Key tasks include:
  - *Parameter Estimation:* Employing robust numerical techniques to find parameter values that minimize the mismatch between model predictions and experimental data. This typically involves using global optimization algorithms (such as enhanced Scatter Search, `eSS`) to mitigate the risk of converging to local optima (underfitting), paired with efficient and stable adaptive ODE solvers to handle potential numerical instabilities (such as blow-up).
  - *Practical Identifiability Analysis* (PIA): Assessing whether model parameters can be reliably estimated from the available experimental data, considering measurement noise and data limitations. This involves analyzing the Fisher Information Matrix (FIM), calculating parameter confidence intervals (e.g., via Cramèr-Rao inequality), and examining parameter correlation matrices to identify poorly determined parameters or strong interdependencies.

– *Dynamic Stability Analysis:* Evaluating the stability properties of the calibrated model (using the estimated parameters) around relevant equilibrium points, typically by analyzing the eigenvalues of the system's Jacobian matrix. This helps ensure the model behaves realistically and avoids biologically implausible, unstable dynamics within the domain of interest.

- **Phase 3: Predictive Power Analysis**
  The final phase focuses on evaluating the model's ability to generalize beyond the training data and assessing the potential impact of overfitting. Key tasks include:

  – *Overfitting Assessment:* Checking for signs that the model has fit the noise in the training data rather than the underlying biological signal. This involves analyzing the distribution and patterns of residuals (differences between predictions and data).

  – *Generalizability Evaluation:* Quantifying the model's predictive performance on data not used during calibration. This is typically achieved through cross-validation, where the model calibrated on a training subset is tested against a separate validation subset, often involving different experimental conditions or time points.

  – *Mechanistic Plausibility Check:* Evaluating whether the estimated parameter values and the model's simulated behavior are consistent with known biological mechanisms and constraints (e.g., signs of interaction terms, magnitudes of rates).

  – *Guidance for Model Refinement:* Using the insights gained from all phases to inform decisions about further model improvement. This might involve simplifying the model structure if overfitting or identifiability issues persist, redesigning experiments to generate more informative data, or collecting additional data to resolve ambiguities.

#### 3.1 Optimization problem

The parameter estimation problem is typically formulated as a nonlinear programming task. The objective is to determine the values of unknown parameters that minimize a specified cost function subject to the constraints imposed by the ODE system and potentially additional algebraic constraints. The cost function represents the model-data mismatch. This function encodes the characteristics of the measurement process and potential prior knowledge. In this study, two cost functions are considered that represent different measures of error, such as least-squares residuals (Eqn. 3) or maximum likelihood estimates (Eqn. 4), depending on the context and available data.

Weighted least squares minimizes the sum of squared residuals, scaled by weights $q_s^{e,o}$ selected by the user:.

$$J_{lsq}(\boldsymbol{\theta}) = \sum_{e=1}^{n_e} \sum_{o=1}^{n_o^e} \sum_{s=1}^{n_s^{e,o}} \left[ \frac{\mathbf{y}_s^{e,o}(\boldsymbol{\theta}, t_s^{e,o}) - \mathbf{ym}_s^{e,o}}{q_s^{e,o}} \right]^2 \tag{3}$$

In frequentist approaches, and under the assumption of independent measurements and normally distributed noise, the log-likelihood function is defined as follows:

$$J_{llk}(\boldsymbol{\theta}) = \sum_{e=1}^{n_e} \sum_{o=1}^{n_o^e} \sum_{s=1}^{n_s^{e,o}} \left( \log(\sigma_s^{e,o}\sqrt{2\pi}) + \frac{1}{2} \frac{(\mathbf{ym}_s^{e,o} - \mathbf{y}_s^{e,o}(\boldsymbol{\theta}, t_s^{e,o}))^2}{(\sigma_s^{e,o})^2} \right) \tag{4}$$

where $\sigma_s^{e,o}$ is the standard deviation of the experimental noise. In the case of known standard deviations, the terms $\log(\sigma_s^{e,o}\sqrt{2\pi})$ are independent of the parameter vector and can be disregarded for parameter optimization and uncertainty analysis, leading to a weighted least squares function in which the weights correspond to the measurements standard deviation:

$$J_{llk}(\boldsymbol{\theta}) = \sum_{e=1}^{n_e} \sum_{o=1}^{n_o^e} \sum_{s=1}^{n_s^{e,o}} \left[ \frac{\mathbf{y}_s^{e,o}(\boldsymbol{\theta}, t_s^{e,o}) - \mathbf{ym}_s^{e,o}}{\sigma_s^{e,o}} \right]^2 \tag{5}$$

#### 3.2 Structural Identifiability Analysis (SIA)

Structural identifiability analysis (SIA) aims to identify model parameters that cannot be uniquely estimated from an ideal dataset, that is, continuous and noise-free (Chis *et al.*, 2011).

An unknown parameter $\theta$ is said to be **Structurally Globally Identifiable** if for almost any $\boldsymbol{\theta}^* \in \boldsymbol{\Theta}$:

$$\sum(\boldsymbol{\theta}) = \sum(\boldsymbol{\theta}^*) \rightarrow \theta_i = \theta_i^* \tag{6}$$

A combination of model and observation states is considered *globally structurally identifiable* if all its unknown parameters can be uniquely estimated from both input and output data, assuming perfect measurements. However, verifying global identifiability is a complex task.

A parameter $\theta$ is **Structurally Locally Identifiable** if, for almost any $\boldsymbol{\theta}^* \in \boldsymbol{\Theta}$, there exists a neighborhood $\mathbf{V}(\boldsymbol{\theta}^*)$ such that:

$$\boldsymbol{\theta} \in \mathbf{V}(\boldsymbol{\theta}^*) \quad and \quad \sum(\boldsymbol{\theta}) = \sum(\boldsymbol{\theta}^*) \rightarrow \theta_i = \theta_i^* \tag{7}$$

A *locally identifiable* parameter can be uniquely inferred within a neighborhood of its nominal value, but a finite number of indistinguishable solutions may exist in the parameter space.

On the other hand, unidentifiability can result in inaccurate values for mechanistically meaningful parameters, as numerical approaches are unable to provide reliable estimates for *non-identifiable* parameters (Rey Barreiro and Villaverde, 2023). An unknown parameter is **Structurally Non-Identifiable** if, for almost any $\boldsymbol{\theta}^* \in \boldsymbol{\Theta}$, there exists no neighborhood $\mathbf{V}(\theta^*)$ such that:

$$\boldsymbol{\theta} \in \mathbf{V}(\boldsymbol{\theta}^*) \quad and \quad \sum(\boldsymbol{\theta}) = \sum(\boldsymbol{\theta}^*) \rightarrow \theta_i = \theta_i^* \tag{8}$$

Structural non-identifiability often arises from the over-parameterization of the model (including its observation function), while practical non-identifiability is typically caused by noise or lack of information in the data (Chiş *et al.*, 2011). Potential approaches to handle structural non-identifiability include (Chis *et al.*, 2011):

- Redefining the model by reducing the number of states and parameters
- Assigning fixed values to certain parameters (especially those less relevant to model predictions)
- Devising new experiments by incorporating measured quantities

In general, dealing with practical identifiability issues is feasible as long as the experimental constraints allow sufficiently comprehensive experiments to be designed.

##### 3.2.1 Tools for Structural Identifiability Analysis

Several software tools are available to study structural identifiability analysis (SIA) of non-linear models, such as: `DAISY` (`Reduce`) (Bellu *et al.*, 2007); `GenSSI` (MATLAB) (Chiş *et al.*, 2011); `EAR` (`Mathematica`) (Karlsson *et al.*, 2012); `COMBOS` (`Maxima`) (Meshkat *et al.*, 2014); `STRIKE-GOLDD` (MATLAB) (Villaverde *et al.*, 2016); `GenSSI2` (MATLAB) (Ligon *et al.*, 2018); `Observability Test` (`Maple`), based on (Sedoglavic, 2001); `SIAN` (`Maple` or Julia) (Hong *et al.*, 2019); `ORC-DF` (MATLAB) (Maes *et al.*, 2019), `rational ORC-DF` or `RORC-DF` (MATLAB) (Shi and Chatzis, 2022); `StrikePy` (`Python`) (Rey Rostro and Villaverde, 2022); `Structural Identifiability` (Julia) (Dong *et al.*, 2023).

Our study required tools capable of assessing both global and local structural identifiability, as well as handling scenarios with unknown initial conditions. Initial screening considered several candidates, including `DAISY` 2007, `COMBOS` 2014, `GenSSI2` 2018, `SIAN`2019, and `Structural Identifiability` 2023. Based on criteria such as reported success rates, computational performance, implementation features, and scope, we selected the following three tools for detailed evaluation in this study: `SIAN`, `Structural Identifiability`, and `GenSSI2`.

`GenSSI2` employs the *Generating Series* approach and *Identifiability Tableaus.* `SIAN` could be seen as a combination of *Differential Algebra* and *Taylor Series* approaches with a correct termination criterion. Both provide information regarding the number of global and local identifiability.

Specifically, we selected the Julia implementation of `SIAN` (`SIAN`, 2019) due to its high success rate, as documented in benchmarking studies (Rey Barreiro and Villaverde, 2023). Although the `Maple` implementation of `SIAN` may be faster for simpler models, it may be less efficient for larger systems. In contrast, Julia's Just-in-Time (JIT) compilation enhances performance for the complex models considered here by compiling functions efficiently upon their first execution. `SIAN` could be seen as a combination of *Differential Algebra* 1994; 2007 and *Taylor Series* 1978 approaches with a correct termination criterion.

We also included `Structural Identifiability` (2023), another Julia package, which has similarly demonstrated strong performance in recent benchmarks (Rey Barreiro and Villaverde, 2023).

Finally, we incorporated `GenSSI2` (2018). While benchmark comparisons indicate that `GenSSI2` often exhibits longer computation times than the modern Julia-based tools, it was historically capable of analyzing a broader range of model types compared to older software such as `DAISY` (2007) or `COMBOS` (2014). It is also the only tool which is fully symbolic, thus no conditions on the parameter feasibility space are required. Therefore, we included `GenSSI2` in our study partly to explore its practical performance limits when applied to scenarios of increasing complexity. It combines the *Generating Series* approach 1982 with the so-called *Identifiability Tableaus* 2010 to assess model identifiability. By computing Lie derivatives of the ODE system, `GenSSI2` generates a nonlinear system of equations on the parameters whose solvability properties provide information about local and global structural identifiability, as well as non-identifiability (Ligon *et al.*, 2018).

#### 3.3 Estimation and Practical Identifiability Analysis (PIA)

Practical identifiability analysis (PIA) attempts to assign unique values to unknown parameters from a given set of experimental data subject to experimental noise. It is worth noting that in the presence of experimental errors, there may be several equivalent solutions that define the confidence region of the parameters. The shape and size of this region will determine whether practical identifiability is guaranteed.

Parameter estimation can be challenging, resulting in poor parameter identifiability (i.e. high inaccuracy in the parameter estimates) due to a lack of sufficiently informative experimental data and the existence of local minima in the objective function. These challenges worsen as the complexity of the system increases, with higher computational costs and more unidentifiable parameters. Incorrect parameter values can significantly compromise model accuracy, leading to erroneous predictions and potentially misleading conclusions (Villaverde *et al.*, 2022).

##### 3.3.1 Global optimization methods

In the case of non-linear ODEs as those describing microbial communities, the corresponding optimization problems are often non-convex. Therefore, global optimization methods are necessary (Banga and Balsa-Canto, 2008). The use of global optimization methods is necessary to guarantee somehow that the best possible solution is located. Usually, several suboptimal solutions are possible.

In this work, the use of a metaheuristic that combines global search heuristics with local optimizers, the so-called `enhanced Scatter Search` (`eSS`), is used as it has shown to have good convergence properties (Egea *et al.*, 2009). Among the local solvers included in the `eSS` implementation, a *Nonlinear Least Squares method* (`nl2sol`) based on the Levenberg-Marquardt algorithm was selected.

##### 3.3.2 Practical Identifiability Analysis (PIA)

Practical identifiability analysis (PIA) does not have a universally agreed-upon definition or a standardized metric, resulting in a variety of methodologies for its assessment. Several approaches are used for uncertainty analysis, including the Fisher Information Matrix approximation ($FIM$), the Bootstrapping method ($BS$), the Profile Likelihood approach ($PL$), and the Multistart technique ($MS$) (Fröhlich *et al.*, 2014).

The Fisher Information Matrix ($FIM$) is defined as follows:

$$\mathcal{F} = \underset{\mathbf{y_m} \mid \mu}{E} \left\{ \left[ \frac{\partial J(\boldsymbol{\theta})}{\partial(\boldsymbol{\theta})} \right] \left[ \frac{\partial J(\boldsymbol{\theta})}{\partial(\boldsymbol{\theta})} \right]^T \right\} \tag{9}$$

$E$ represents expected values and $\mu$ a value of the parameters, hopefully close to their real value. The diagonal of the covariance matrix approximates the confidence intervals for the parameters. The correlation between parameters can be calculated with $C_{r_{ij}} = \frac{C_{ij}}{\sqrt{C_{ii}C_{jj}}}$, the parameters $ij$ are highly correlated if $C_{r_{ij}} = 1$, if $C_{r_{ij}} = 0$ are fully uncorrelated. The confidence intervals obtained will help to decide whether further experiments are needed to improve the parameter estimates.

##### 3.3.3 Goodness of fit. Underfitting and overfitting.

Goodness-of-fit analyzes include metrics like the root mean square error ($RMSE$), Akaike Information Criterion ($AIC$), Bayesian Information Criterion ($BIC$), and $R^2$, these metrics offer complementary perspectives on model validity.

The chosen metric for evaluating the quality of the ensemble predictions is the normalized root mean square error ($NRMSE$) (Massonis *et al.*, 2022):

$$NRMSE(\mathbf{y}) = \frac{RMSE(\mathbf{y_s^{e,o}})}{max(\mathbf{ym_s^{e,o}}) - min(\mathbf{ym_s^{e,o}})} \tag{10}$$

where:

$$RMSE(\mathbf{y_s^{e,o}}) = \sqrt{\frac{\sum_{i=1}^{n_s^{e,o}} (\mathbf{y}_s^{e,o} - \mathbf{ym}_s^{e,o})^2}{n_s^{e,o}}} \tag{11}$$

##### 3.3.4 Sensitivity analysis

It may be useful to explore how different parameters affect the predictions made by the model. We conducted sensitivity analyzes to determine which parameters have a greater impact on each state variable. By ranking the parameters, we can decide which ones need to be fixed (if necessary). The significance of the parameters can be measured using parametric sensitivities. The local parametric sensitivities for a specific experiment ($e$), observable ($o$), and sampling time ($t_s$), correspond to the following equation:

$$S_\theta^{e,o}(t_s^{e,o}) = \frac{\partial \mathbf{y}^{e,o}}{\partial \theta}(t_s^{e,o}); \quad \theta = 1, \ldots, n_\theta \tag{12}$$

To take into account differences in orders of magnitude in the observables and parameters, it is advised to use full relative sensitivities:

$$S_\theta^{e,o}(t_s^{e,o}) = \frac{\Delta\theta}{\Delta y^{e,o}} \frac{\partial \mathbf{y}^{e,o}}{\partial \theta}(t_s^{e,o}); \quad \theta = 1, \ldots, n_\theta \tag{13}$$

where $\Delta\theta$ and $\Delta y^{e,o}$ have the same order of magnitude than $\theta$ and $y^{e,o}$ at the optimum.

In our workflow, we use $\delta^{mean}$ to quantify how sensitive a model is to a given parameter:

$$\delta_p^{mean} = mean(s_p^{e,o}(t_s^{e,o})) \qquad \forall \quad e = 1 \ldots n_e; \; o = 1 \ldots n_o; s = 1 \ldots n_s \tag{14}$$

##### 3.3.5 Tools for estimation and practical identifiability analysis

These problems can be stated and efficiently solved using `AMIGO2` (*Advanced Model Identification using Global optimization*) toolbox for MATLAB, which is specifically oriented to dynamic modeling, dynamic optimization and inverse optimal control. In the context of model calibration, the tool allows for global and local sensitivity analyzes, ranking of parameters, practical identifiability tests, parameter estimation using global and local optimization methods and model-based optimal experimental design (Balsa-Canto *et al.*, 2010, 2016).

`AMIGO2` is equipped with advanced initial value problem solvers ($IVPs$) and non-linear optimization methods ($NLPs$). It can handle both parameter estimation and model-based experimental design problems. For $IVPs$, explicit and implicit Runge-Kutta, Adams-Bashforth, and $BDF$ methods are included along with sensitivity computation techniques. As far as $NLP$ solvers are concerned, `AMIGO2` offers a range of direct and indirect local methods, multistart of local methods, global stochastic, and hybrid optimization methods. For computationally demanding tasks, `C` compiled code can be used.

In the proposed framework we used `AMIGO2_SData` to generate synthetic experimental data, `AMIGO2_PE` for parameter estimation. Note that the output of `AMIGO2_PE` includes the FIM plus the confidence intervals for the optimal parameters. `AMIGO2_LRank` was used to compute local parametric sensitivities. In the examples we will show the mean local sensitivities but other measures of the parametric sensitivities are automatically provided. Goodness-of-fit was evaluated using `AMIGO2_postAnalysis`. All the steps are illustrated with the live scripts accompanying the manuscript.

#### 3.4 Predictive power of calibrated models

While the parameter estimation with global methods aims to identify the optimal set of parameters that results in low RMSE values or $R^2$ close to 1, the quality of the resulting fit must still be critically evaluated.

To ensure reliable predictions, it is crucial to identify and mitigate both underfitting and overfitting. Underfitting occurs when the model is too simple to capture the underlying dynamics of the data, leading to poor performance on both the training and validation sets, often due to insufficient model complexity or inadequate feature representation. Note that in synthetic examples as the once considered here, the model does indeed explain the data. Thus underfitting arises from local convergence of the optimizer.

Overfitting, on the other hand, arises when the model is overly complex and captures noise or specific variations in the training dataset rather than the generalizable patterns. While an overfit model may perform well on the training data, its predictive power on unseen data is significantly diminished.

The predictive power of a model can be assessed using techniques like cross-validation, which partitions the dataset into subsets, one used for validation while the others serve as training data. Examples of this approach can be found in sections 6.6.6 and 6.10.6. To diagnose underfitting and overfitting, we employ methods like cross-validation and residual analysis. Key performance metrics, including root mean squared error ($RMSE$), mean absolute error ($MAE$), and $R^2$, provide valuable insights into the model's bias-variance trade-off: high bias often indicates underfitting, while high variance suggests overfitting.

#### 3.5 Stability analysis

In specific cases, a stability analysis is conducted to identify equilibrium points and characterize the system's behavior in their vicinity. This process involves linearizing the model via the Jacobian matrix, which captures the system's local dynamics around equilibrium points. The stability of these points is determined by analyzing the eigenvalues of the Jacobian. Additionally, numerical simulations can provide deeper insights into global dynamics, including periodic orbits and potential chaotic behavior.

We illustrate the stability analysis using a *Generalized Lotka-Volterra* model with three interacting species. The system is described by the following equations:

$$f_1 = \frac{dX_1}{dt} = \mu_1 X_1 + \beta_{11} X_1^2 + \beta_{12} X_1 X_2 + \beta_{13} X_1 X_3$$

$$f_2 = \frac{dX_2}{dt} = \mu_2 X_2 + \beta_{21} X_1 X_2 + \beta_{22} X_2^2 + \beta_{23} X_2 X_3$$

$$f_3 = \frac{dX_3}{dt} = \mu_3 X_3 + \beta_{31} X_1 X_3 + \beta_{32} X_2 X_3 + \beta_{33} X_3^2$$

where:

- $X_1, X_2, X_3$ represent the populations of the three species.
- $\mu_i$ are the growth rates.
- $\beta_{ij}$ are the interaction coefficients.

The stability analysis is carried out following these steps:

- **1. Find equilibrium points**
  Equilibrium points are found by solving:

  $$\frac{dX_1}{dt} = 0, \quad \frac{dX_2}{dt} = 0, \quad \frac{dX_3}{dt} = 0$$

  This results in equilibrium points $(X_1^*, X_2^*, X_3^*)$, where the system remains at rest. Solving these equations often involves handling nonlinear systems.

- **2. Linearize the system around equilibrium points**
  To analyze stability, the system is linearized using the Jacobian matrix:

  $$J = \begin{bmatrix} \frac{\partial f_1}{\partial X_1} & \frac{\partial f_1}{\partial X_2} & \frac{\partial f_1}{\partial X_3} \\ \frac{\partial f_2}{\partial X_1} & \frac{\partial f_2}{\partial X_2} & \frac{\partial f_2}{\partial X_3} \\ \frac{\partial f_3}{\partial X_1} & \frac{\partial f_3}{\partial X_2} & \frac{\partial f_3}{\partial X_3} \end{bmatrix}$$

  Here, $f_i$ represents the right-hand side of the ODEs. The Jacobian is evaluated at each equilibrium point.

- **3. analyze the eigenvalues of the Jacobian**
  The stability of the equilibrium depends on the eigenvalues of the Jacobian:

  - If **all eigenvalues have negative real parts**, the equilibrium is **asymptotically stable** (locally attracting).
  - If **any eigenvalue has a positive real part**, the equilibrium is **unstable**.
  - If **eigenvalues have zero real parts**, higher-order terms or nonlinear techniques (e.g., Lyapunov methods) are needed to determine stability.

- **4. Classify the equilibrium**
  Based on the eigenvalues, the equilibrium can be classified as:

  - **Node**: Real eigenvalues, all negative (stable) or all positive (unstable).

- **Spiral/Sink/Source**: Complex eigenvalues with negative real parts (stable spiral) or positive real parts (unstable spiral).
- **Saddle point**: Mixed signs in real eigenvalues (unstable).

- **5. (Optional) Perform numerical simulations**
  For complex systems, numerical simulations might be useful to explore global dynamics, periodic orbits, or chaotic behavior.

This structured approach provides a clear and concise method for analyzing the stability of dynamical systems, particularly in ecological models like the *Generalized Lotka-Volterra* system. We provide a MATLAB implementation of this procedure (*GLV3_Stability* in the folder *Practical_Identifibility_Analysis*).

### 4 Software implementation

We developed our workflow as a unified, Matlab-based pipeline that integrates the previously mentioned three tools for structural identifiability analysis (SIA) alongside estimation and practical identifiability analysis (PIA) methods using `AMIGO2` Balsa-Canto *et al.* (2016). `AMIGO2` (*Advanced Model Identification using Global optimization*) is a powerful toolbox for dynamic modeling and optimization, offering a broad selection of nonlinear optimization solvers. These include direct and indirect local methods, multi-start local approaches, global stochastic algorithms, and hybrid optimization techniques. Additionally, `AMIGO2` facilitates PIA.

To enhance computational efficiency, particularly for demanding tasks such as parameter estimation, using `AMIGO2` in this framework enables automatic generation of `C`-compiled code, significantly improving performance. Furthermore, we extended the pipeline with additional code to support other key steps, such as stability analysis. Several case studies, based on commonly used models of microbial communities, were implemented and tested within this workflow.

The resulting integrated software is available in both electronic notebook format (live scripts) and as standard MATLAB scripts. It features the following key components:

- **Pre-estimation analysis:** the workflow simplifies model definition, data integration, and identifiability analysis, ensuring a solid foundation for parameter estimation. This module efficiently handles multi-experiment datasets, incorporates data visualization tools, and facilitates structural identifiability analysis using several methods, so non-expert users can determine whether model parameters can be uniquely inferred.

- **Robust parameter estimation:** a diverse set of global and local optimization algorithms enhances the reliability and accuracy of parameter estimation. Sensitivity analysis, practical identifiability assessment, and stability analysis help refine parameter estimates. The framework also supports easy cross-validation with additional datasets and leverages parallel computing for improved efficiency and scalability.

- **Predictive power analysis:** to ensure the model produces reliable and meaningful predictions, the workflow includes sensitivity analysis, statistical goodness-of-fit tests and cross-validation to assess model performance. These methods enable model comparison, interactive plotting, time series visualization, and statistical summarization, aiding in result interpretation. Furthermore, automated report generation and documentation via electronic notebooks streamline reproducibility and facilitate effective communication of findings.

### 5 Software Installation

This section provides instructions for installing the software packages required to perform the structural and practical identifiability analyzes described in our workflow.

#### 5.1 Structural Identifiability Analysis (SIA)

To facilitate structural identifiability analysis (SIA) across different tools, we developed helper scripts. These scripts translate model definitions from the `AMIGO2` syntax into the specific input formats required by `GenSSI2` (MATLAB), `SIAN` (Julia), and `Structural Identifiability` (Julia).

Taking an existing `AMIGO2` model definition as input, these functions automatically generate the necessary files and execution commands for each of the three SIA tools. This automation streamlines the analysis process, enabling straightforward application of SIA using any of these platforms. For further details, please consult the `Readme` file located in the `Structural_Identifiability_Analysis` folder.

##### 5.1.1 GenSSI2

To install GenSSI2:

1. Download the .zip archive containing the MATLAB package from the official GitHub repository: https://github.com/genssi-developer/GenSSI.git.
2. Unzip the archive.
3. Add the extracted GenSSI2 folder to your MATLAB path.
4. Ensure you run the genssiStartup.m script within MATLAB before using the toolbox functions or running examples.

##### 5.1.2 SIAN

The stable version of SIAN can be installed directly within a Julia environment using the package manager:

```
# Enter Pkg mode by typing ']'
pkg> add SIAN

# Exit Pkg mode (Backspace)
julia> using SIAN
```

For comprehensive installation guidance, refer to the official documentation:
https://github.com/alexeyovchinnikov/SIAN-Julia.

##### 5.1.3 Structural Identifiability

To install the Structural Identifiability package for Julia, follow the instructions provided in the official documentation: https://docs.sciml.ai/StructuralIdentifiability/stable/. We recommend using version 0.5.8 or later for compatibility with our workflow.

Alternatively, install a specific version directly using the Julia package manager:

```
using Pkg
Pkg.add(name="StructuralIdentifiability", version="0.5.8")
# Or Pkg.add("StructuralIdentifiability") for the latest stable version

julia> using StructuralIdentifiability
```

#### 5.2 Estimation and Practical Identifiability Analysis: AMIGO2

The AMIGO2 toolbox, used here for parameter estimation and practical identifiability analysis, can be installed as follows:

1. Download the toolbox archive (.zip file) from the official website: https://sites.google.com/site/amigo2toolbox/download.
2. Extract the contents of the .zip file.
3. Before running any examples or using the toolbox, execute the startup script AMIGO_Startup.m from within MATLAB to ensure all necessary paths are configured correctly.

### 6 Results

#### 6.1 Computational resources

Structural identifiability analyzes were performed on a PC workstation equipped with dual Intel Xeon Silver 4210R CPUs operating at 2.40 GHz, 256 GB of RAM, and running Windows 11 Pro 64-bit alongside MATLAB R2024. Practical identifiability analyzes utilized both this workstation and an additional PC featuring a 13th Gen Intel Core i5-13500 CPU running at 2.50 GHz, 64 GB of RAM, also operating on Windows 11 Pro 64-bit with MATLAB R2024. Based on published benchmarks (PassMark), the PC with the i5 CPU offers roughly twice the performance.

#### 6.2 Overview of case studies

This section provides a comprehensive analysis of a set of canonical models of increasing complexity as representatives of the most common frameworks: from the most classical and simple ecological models (such as *Lotka-Volterra*), to more complex models accounting for nutrients dynamics (such as food web models), and coarse-grained models incorporating different regulatory mechanisms. The results include structural and practical identifiability assessments, model performance evaluations, and insights into parameter stability and predictive power. Each subsection focuses on a specific model, summarized in Table A.

| | ***GLV2_Comp***, ***GLV2_Coex*** | ***GLV3*** | ***MGLV*** | ***CompV*** | ***PCM*** | ***EPHP*** |
|---|---|---|---|---|---|---|
| Original ref. | Dimas Martins and Gjini (2020) | Remien *et al.* (2021) | Venturelli *et al.* (2018) | Alsolami and El Hajji (2023) | Li *et al.* (2020) | Mauri *et al.* (2020) |
| Description | *Generalized Lotka-Volterra* 2 species | *Generalized Lotka-Volterra* 3 species | *Generalized Lotka-Volterra* 12 species | *Competition with a virus* | *Phage Cocktail Model* | *Enhanced Production of Heterologous Proteins* |
| Microbial community | Influenza virus | Synthetic community | Gut Microbiome | 2 bacteria | Bacteria and phages | E.coli |
| $n_x$ | 2 | 3 | 12 | 4 | 5 | 5 |
| $n_p$ | 6 | 12 | 156 | 7 | 13 | 19 |
| Data type | Simulated | Simulated | Simulated | Simulated | Simulated | Simulated |
| Noise Level | 0%, 5%, 10% | 0%, 5%, 10% | 0%, 5%, 10% | 0%, 5%, 10% | 0%, 5%, 10% | 0%, 5% |
| Data points | ***GLV2_Comp***: 60, 10, 10<br>***GLV2_Coex***: 60, 10, 10 | 300, 10, 10 | 100, 20, 20 | 30, 10, 10 | 150, 10, 10 | 150, 10 ,10 |
| Noise type | ***GLV2_Comp***: Heteroscedastic<br>***GLV2_Coex***: Homoscedastic | Homoscedastic | Homoscedastic | Homoscedastic | Heteroscedastic | Homoscedastic |

Table A: Selected cases for model calibration under fully observed scenarios, considering various noise levels (0%, 5%, and 10%). ***GLV*** refers to the *Generalized Lotka-Volterra* model, $n_x$ denotes the number of variables, $n_p$ the number of parameters, and the data points represent the number of observations per variable.

#### 6.3 Data

We created a set of identification problems using the selected case studies described above. For each model, we generated synthetic (pseudo-experimental) data, incorporating different types and levels of noise to mimic realistic experimental conditions. This approach allows us to assess the robustness of our methodology and identify potential challenges arising from data variability. Synthetic data plays a crucial role in testing and validating identification workflows for dynamic biological models by providing a controlled environment where the true parameter values and system behavior are known. Working with synthetic data, we can generate observations from a model using predetermined (nominal) parameter values, potentially adding realistic measurement noise to simulate experimental conditions. This ground truth knowledge allows for a direct assessment of how well parameter estimation methods can recover the nominal parameters, making it possible to evaluate the accuracy, precision, and reliability of the identification approach.

Three sets of *pseudo*-experimental data were generated for each model (refers to Table A) with varying Gaussian noise levels (0%, 5%, 10%). First, we consider an ideal measurement method that provides a large number of data points with no Gaussian noise (0%). Then, we examine more realistic scenarios, with fewer data points with 5% and 10% Gaussian noise. The **log-likelihood function** was employed for noisy data, while the **weighted least squares function** was used for noise-free data. Depending on the problem characteristics the noise was *homoscedastic* or *heteroscedastic*. Homoscedasticity implies that the spread of the residuals is the same across the entire range of the observed variable(s), while heteroscedasticity implies that the spread of the residuals changes across different levels of the observed variable(s).

#### 6.4 *Generalized Lotka-Volterra* Models

The *Lotka-Volterra* model is one of the simplest ecological models and was originally used to describe macrofaunal predator–prey dynamics. *Generalized Lotka-Volterra* models extend this framework, encompassing a broader class of equations beyond the classic competitive or predator-prey cases. They consist of a set of nonlinear, coupled, first-order differential equations.

The growth rate of a species' population is determined by its intrinsic growth rate and the linear interactions with other species. *Lotka–Volterra* (***LV***) models have been widely used to describe the temporal evolution of ecosystems. Back in 1987, Hofbauer *et al.* (1987) expanded the Lotka-Volterra equations to encompass an arbitrary number of coexisting species, providing deeper insights into the mathematical properties of ***GLV*** models. Thanks to the advancements in sequencing techniques, ***GLV*** models have recently been employed to investigate the temporal dynamics of diverse bacterial communities (Succurro and Ebenhöh, 2018).

##### 6.4.1 Model

The *Generalized Lotka-Volterra* (***GLV***) model predicts the dynamic behaviors of microbial communities. The model assumes the microbial population is homogeneous (i.e. microorganisms grow simultaneously) and that interactions happen within the same species (intraspecific competition) and with other species (interspecific competition). The model is given by the following equation:

$$\frac{dX_i}{dt} = X_i \left( \mu_i + \sum_{j=1}^{n} \beta_{ij} X_j \right) \tag{15}$$

where $n$ represents the number of species, $\mu$ positive growth rates, $\beta_{ii}$ and $\beta_{ij}$, the intraspecies and interspecies interaction coefficients respectively, $\beta_{ij}$ describes how microbe $j$ affects the growth rate of microbe $i$. Typically, the parameters $\beta_{ii}$ are constrained to be negative so that the carrying capacity ($k_i = -\mu_i/\beta_{i,i}$) is positive, but this is not required for the structural identifiability analysis (SIA).

The ***GLV*** equations can accommodate all possible combinations of interaction signs between species pairs ($-/-$, $-/+$, $+/-$, $+/+$). It should be noted that in this model the spatial distribution of the population is disregarded. This is a suitable approximation when, for example, a microbial community grows in a well-stirred bioreactor.

In this study, we will focus on ***GLV*** models with two (section 6.5), three (section 6.6) and twelve species (section 6.10).

#### 6.5 *GLV2*: *Generalized Lotka-Volterra* 2 species

##### 6.5.1 Model

Assuming a two-species population with only two-way interactions between microbes (as in Figure A), the change in density over time of the species populations can be described by the following system:

$$\frac{dX_1}{dt} = \mu_1 X_1 - \beta_{11} \cdot 10^{-6} X_1^2 - \beta_{12} \cdot 10^{-6} X_1 X_2 \tag{16}$$

$$\frac{dX_2}{dt} = \mu_2 X_2 - \beta_{21} \cdot 10^{-6} X_1 X_2 - \beta_{22} \cdot 10^{-6} X_2^2 \tag{17}$$

$X_1$ and $X_2$ represent two strains of the *Influenza* virus. The parameters were obtained from Dimas Martins and Gjini (2020), who derived them using experimental data from McCaw *et al.* (2011). To maintain a consistent order of magnitude across all parameters, all $\beta$ values are scaled by $10^{-6}$. To be faithful to the original paper, we consider all $\beta$ positive, preceded by the sign - in the equations.

Each strain grows bounded by its carrying capacity due to limiting factors such as finite resources or space, according to logistic growth in the absence of the other strain.

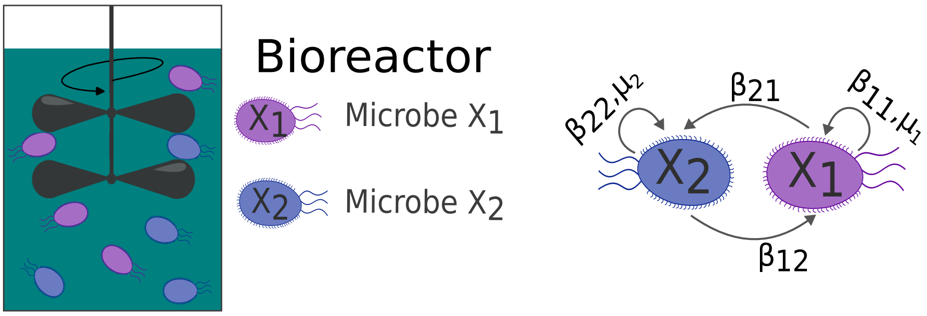

Figure A: A bioreactor with two microbe populations ($X_1$, $X_2$) interacting with each other (**Left**). Interactions between two microorganisms (**Right**).

Two scenarios were considered, a scenario of competitive exclusion (case study ***GLV2_Comp***), and a scenario of coexistence (case study ***GLV2_Coex***). Parameters and initial conditions (IC) for both cases are summarized in Table B (coexistence) and Table C (competition).

**Case study: Coexistence (*GLV2_Coex*)**

| **Parameter** | **Value [AU]** | **Parameter** | **Value [AU]** | **Parameter** | **Value [AU]** |
|---|---|---|---|---|---|
| $\mu_1$ | 6.00 | $\beta_{1,1}$ | 6.00 | $\beta_{2,1}$ | 2.40 |
| $\mu_2$ | 4.80 | $\beta_{1,2}$ | 1.20 | $\beta_{2,2}$ | 4.80 |
| **Initial conditions (IC):** $X_{10} = 0.50$, $X_{20} = 0.50$ | | | | | |

Table B: Case study ***GLV2_Coex***: Parameter values and initial conditions (IC) for the case study of the *Generalized Lotka-Volterra* two-species model in the coexistence scenario. The parameters ($\mu_i$, $\beta_{i,j}$) represent growth and interaction rates, respectively, while the initial conditions ($X_{i0}$) denote the starting population densities. Values are taken from Dimas Martins and Gjini (2020).

**Case study: Competition (*GLV2_Comp*)**

| **Parameter** | **Value [AU]** | **Parameter** | **Value [AU]** | **Parameter** | **Value [AU]** |
|---|---|---|---|---|---|
| $\mu_1$ | 6.00 | $\beta_{1,1}$ | 6.00 | $\beta_{2,1}$ | 6.24 |
| $\mu_2$ | 4.80 | $\beta_{1,2}$ | 5.40 | $\beta_{2,2}$ | 4.80 |
| **Initial conditions (IC):** $X_{10} = 0.50$, $X_{20} = 0.50$ | | | | | |

Table C: Case study ***GLV2_Comp***: Parameter values and initial conditions (IC) for the case study of the *Generalized Lotka-Volterra* two-species model in the competition scenario. The parameters ($\mu_i$, $\beta_{i,j}$) represent growth and interaction rates, respectively, while the initial conditions ($X_{i0}$) denote the starting population densities. Values are taken from Dimas Martins and Gjini (2020).

##### 6.5.2 Structural Identifiability Analysis (SIA)

We analyzed three observation scenarios: one fully observed ($X_1$,$X_2$) and two partially observed ($X_1 + X_2$; $X_1$). Since `GenSSI2` is a fully symbolic tool, its execution times are typically longer and may fail to solve the system of equations. `Structural Identifiability` proved to be the most efficient tool, delivering the shortest execution times and consistent results in all cases.

In the scenario where only one population ($X_1$) was observed, `SIAN` and `Structural Identifiability` determined that parameters $X_2$, $\beta_{22}$, $\beta_{12}$ were non-identifiable, while parameters $X_1$, $\beta_{11}$, $\beta_{21}$, $\mu_i$ were found to be globally identifiable (Table E). Meanwhile, `GenSSI2` was unable to provide a solution. Even after 24 hours, no solution was reached (Table D).

| | `GenSSI2` (**IC Unknown**) | | | | `GenSSI2` (**IC Known**) | | | |
|---|---|---|---|---|---|---|---|---|
| Obs. | Global | Local | Non-Id | Time[s] | Global | Local | Non-Id | Time[s] |
| $X_1$,$X_2$ | $X_{i0}$,$\mu_i$, $\beta_{ii}$,$\beta_{ij}$ | - | - | 7.38 | $\mu_i$, $\beta_{ii}$,$\beta_{ij}$ | - | - | 7.67 |
| $X_1 + X_2$ | - | $X_{i0}$,$\mu_i$, $\beta_{ii}$,$\beta_{ij}$ | - | 105.46 | - | $\mu_i$,$\beta_{ii}$,$\beta_{ij}$ | - | 551.65 |
| $X_1$ | - | - | - | *OOM* | - | - | - | |

Table D: Case study ***GLV2***: Structural identifiability analysis (SIA) results obtained with `GenSSI2` for the *Generalized Lotka-Volterra* 2 species model. The analysis was performed considering three observation scenarios ($X_1$, $X_2$; $X_1 + X_2$; $X_1$) and two cases for initial conditions: unknown (IC Unknown) and known (IC Known). The parameters are classified as globally identifiable ("Global"), locally identifiable ("Local"), or non-identifiable ("Non-Id"). A " - " indicates that no parameters are found in that category. The table summarizes the computational time and the identifiability results for the model parameters ($\mu_i$, $\beta_{ii}$, $\beta_{ij}$).

| **IC Unknown** | `SIAN` | | | | `Structural Identifiability` | | | |
|---|---|---|---|---|---|---|---|---|
| Obs. | Global | Local | Non-Id | Time[s] | Global | Local | Non-Id | Time[s] |
| $X_1$,$X_2$ | $X_i$,$\mu_i$, $\beta_{ii}$,$\beta_{ij}$ | - | - | 57.86 | $X_i(t)$,$\mu_i$, $\beta_{ii}$,$\beta_{ij}$ | - | - | 12.84 |
| $X_1 + X_2$ | - | $X_i$,$\mu_i$, $\beta_{ii}$,$\beta_{ij}$ | - | 160.46 | - | $X_i(t)$,$\mu_i$, $\beta_{ii}$,$\beta_{ij}$ | - | 12.82 |
| $X_1$ | $X_1$,$\mu_i$, $\beta_{11}$,$\beta_{21}$ | - | $X_2$, $\beta_{12}$,$\beta_{22}$ | 56.70 | $X_1(t)$,$\mu_i$, $\beta_{11}$,$\beta_{21}$ | - | $X_2(t)$, $\beta_{12}$,$\beta_{22}$ | 12.43 |

Table E: Case study ***GLV2***: Structural identifiability analysis (SIA) results obtained with `SIAN` and `Structural Identifiability` for the *Generalized Lotka-Volterra* 2 species model. The analysis was performed considering three observation scenarios ($X_1$, $X_2$; $X_1 + X_2$; $X_1$) and unknown initial conditions (IC Unknown). The parameters are classified as globally identifiable ("Global"), locally identifiable ("Local"), or non-identifiable ("Non-Id"). A " - " indicates that no parameters are found in that category. The table summarizes the computational time and the identifiability results for the model parameters ($\mu_i$, $\beta_{ii}$, $\beta_{ij}$).

| **IC Known** | `SIAN` | | | | `Structural Identifiability` | | | |
|---|---|---|---|---|---|---|---|---|
| Obs. | Global | Local | Non-Id | Time[s] | Global | Local | Non-Id | Time[s] |
| $X_1$,$X_2$ | $\mu_i$,$\beta_{ii}$, $\beta_{ij}$ | - | - | 53.52 | $X_{i0}$, $\mu_i$, $\beta_{ii}$,$\beta_{ij}$ | - | - | 15.42 |
| $X_1 + X_2$ | - | $\mu_i$,$\beta_{ii}$ $\beta_{ij}$ | - | 154.24 | - | $X_{i0}$,$\mu_i$, $\beta_{ii}$,$\beta_{ij}$ | - | 16.66 |
| $X_1$ | $\mu_i$, $\beta_{11}$, $\beta_{21}$ | - | $\beta_{12}$, $\beta_{22}$ | 55.76 | $X_{10}$, $\mu_i$, $\beta_{11}$, $\beta_{21}$ | - | $X_{20}$, $\beta_{12}$, $\beta_{22}$ | 14.11 |

Table F: Case study ***GLV2***: Structural identifiability analysis (SIA) results obtained with `SIAN` and `Structural Identifiability` for the *Generalized Lotka-Volterra* 2 species model. The analysis was performed considering three observation scenarios ($X_1$, $X_2$; $X_1 + X_2$; $X_1$) and known initial conditions (IC Known). The parameters are classified as globally identifiable ("Global"), locally identifiable ("Local"), or non-identifiable ("Non-Id"). A " - " indicates that no parameters are found in that category. The table summarizes the computational time and the identifiability results for the model parameters ($\mu_i$, $\beta_{ii}$, $\beta_{ij}$).

##### 6.5.3 Practical Identifiability Analysis (PIA)

We addressed a parameter estimation problem involving two observed populations, $X_1$ and $X_2$, with unknown initial conditions ($X_{10}$, $X_{20}$) (Figure B, Figure D). To assess the robustness of our estimation methods, we generated three sets of pseudo-experimental data, each with a different Gaussian noise level: 0%, 5%, 10%.

In the coexistence case study, the generated data was homoscedastic and positive, reflecting the nature of population measurements. In contrast, the data for the competition case study was heteroscedastic and positive. For the dataset with 0% noise, we employed a weighted least squares approach as the cost function for parameter estimation. For datasets with 5% and 10% noise levels, we used a cost function based on the negative log-likelihood, which is better suited for handling data with higher noise levels.

**Case study: Competition**

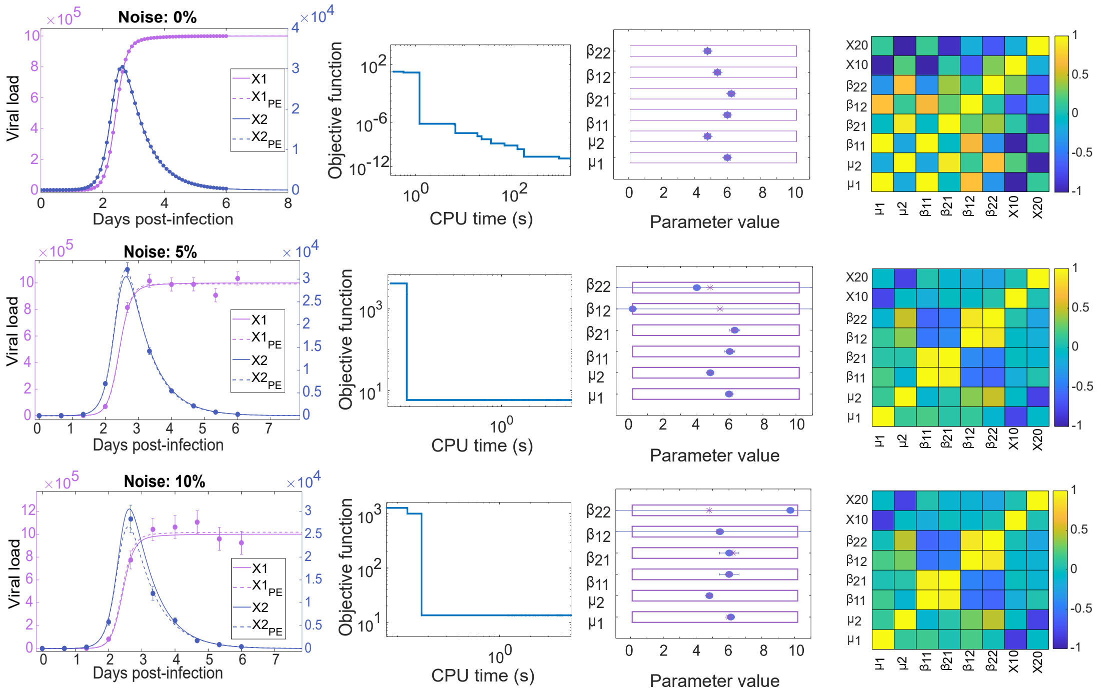

Figure B: Case study ***GLV2_Comp***: Model calibration results for the competition scenario of the *Generalized Lotka-Volterra* two species model (Dimas Martins and Gjini, 2020). The first row displays results for the pseudo-experimental data with 0% noise, the second row shows results for 5% noise, and the third row corresponds to 10% noise. In each case, the simulation using the estimated parameters is compared to the simulation with nominal parameters, along with the convergence curve, correlation matrix, and a comparison of nominal and estimated parameters. These results demonstrate the model's ability to accurately fit the data under varying noise levels.

After estimating the parameters, we conducted a sensitivity analysis to evaluate their influence on the population dynamics. This involved calculating the mean relative sensitivity and perturbing individually each parameter by +5% to identify which ones have the greatest impact on species population sizes (Figure C).

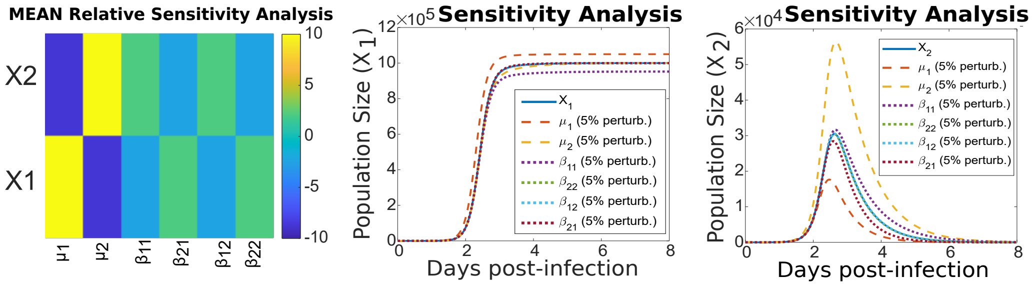

Figure C: Case study ***GLV2_Comp***: Sensitivity analysis for the competition scenario of a two species *Generalized Lotka-Volterra*. **Left**: Mean relative sensitivity analysis. **Center**: Simulations of species $X_1$ with a positive 5% perturbation of each parameter. **Right**: Simulations of species $X_2$ with a positive 5% perturbation of each parameter.

| | **LB** | **UB** | **Nominal** | **Estimated** | **EA** | **ER (%)** |
|---|---|---|---|---|---|---|
| $\mu_1$ | $1.00 \times 10^{-1}$ | $1.00 \times 10^{1}$ | 6.00 | 6.00 | $1.66 \times 10^{-6}$ | $5.07 \times 10^{-6}$ |
| $\mu_2$ | $1.00 \times 10^{-1}$ | $1.00 \times 10^{1}$ | 4.80 | 4.80 | $1.15 \times 10^{-6}$ | $2.33 \times 10^{-5}$ |
| $\beta_{11}$ | $1.00 \times 10^{-1}$ | $1.00 \times 10^{1}$ | 6.00 | 6.00 | $1.63 \times 10^{-6}$ | $3.11 \times 10^{-6}$ |
| $\beta_{21}$ | $1.00 \times 10^{-1}$ | $1.00 \times 10^{1}$ | 6.24 | 6.24 | $1.03 \times 10^{-6}$ | $1.92 \times 10^{-5}$ |
| $\beta_{12}$ | $1.00 \times 10^{-1}$ | $1.00 \times 10^{1}$ | 5.40 | 5.40 | $1.71 \times 10^{-5}$ | $1.86 \times 10^{-4}$ |
| $\beta_{22}$ | $1.00 \times 10^{-1}$ | $1.00 \times 10^{1}$ | 4.80 | 4.80 | $2.62 \times 10^{-5}$ | $2.96 \times 10^{-4}$ |
| $X_{10}$ | $1.00 \times 10^{-2}$ | 1.00 | $5.00 \times 10^{-1}$ | $5.00 \times 10^{-1}$ | $1.94 \times 10^{-6}$ | $9.25 \times 10^{-4}$ |
| $X_{20}$ | $1.00 \times 10^{-2}$ | 1.00 | $5.00 \times 10^{-1}$ | $5.00 \times 10^{-1}$ | $1.22 \times 10^{-6}$ | $4.37 \times 10^{-5}$ |

Table G: Case study ***GLV2_Comp***: Results for the *Generalized Lotka-Volterra 2 species* model with 0% heteroscedastic Gaussian noise. The table presents the lower and upper bounds (**LB**, **UB**), nominal and estimated values, as well as the absolute and relative errors (**EA**, **ER**), for each estimated parameter.

| | **LB** | **UB** | **Nominal** | **Estimated** | **EA** | **ER (%)** |
|---|---|---|---|---|---|---|
| $\mu_1$ | $1.00 \times 10^{-1}$ | $1.00 \times 10^{1}$ | 6.00 | 5.94 | $6.52 \times 10^{-2}$ | 1.1 |
| $\mu_2$ | $1.00 \times 10^{-1}$ | $1.00 \times 10^{1}$ | 4.80 | 4.81 | $6.74 \times 10^{-2}$ | 1.4 |
| $\beta_{11}$ | $1.00 \times 10^{-1}$ | $1.00 \times 10^{1}$ | 6.00 | 5.99 | $3.02 \times 10^{-1}$ | 5.05 |
| $\beta_{21}$ | $1.00 \times 10^{-1}$ | $1.00 \times 10^{1}$ | 6.24 | 6.30 | $3.07 \times 10^{-1}$ | 4.88 |
| $\beta_{12}$ | $1.00 \times 10^{-1}$ | $1.00 \times 10^{1}$ | 5.40 | $1.00 \times 10^{-1}$ | $1.28 \times 10^{1}$ | $1.28 \times 10^{4}$ |
| $\beta_{22}$ | $1.00 \times 10^{-1}$ | $1.00 \times 10^{1}$ | 4.80 | 3.99 | $1.47 \times 10^{1}$ | $3.68 \times 10^{2}$ |
| $X_{10}$ | $1.00 \times 10^{-2}$ | 1.00 | $5.00 \times 10^{-1}$ | $5.23 \times 10^{-1}$ | $4.13 \times 10^{-2}$ | 7.9 |
| $X_{20}$ | $1.00 \times 10^{-2}$ | 1.00 | $5.00 \times 10^{-1}$ | $4.93 \times 10^{-1}$ | $4.09 \times 10^{-2}$ | 8.29 |

Table H: Case study ***GLV2_Comp***: Results for the *Generalized Lotka-Volterra 2 species* model with 5% heteroscedastic Gaussian noise. The table presents the lower and upper bounds (**LB**, **UB**), nominal and estimated values, as well as the absolute and relative errors (**EA**, **ER**), for each estimated parameter.

| | LB | UB | Nominal | Estimated | EA | ER (%) |
|---|---|---|---|---|---|---|
| $\mu_1$ | $1.00 \times 10^{-1}$ | $1.00 \times 10^{1}$ | 6.00 | 6.12 | $1.38 \times 10^{-1}$ | 2.26 |
| $\mu_2$ | $1.00 \times 10^{-1}$ | $1.00 \times 10^{1}$ | 4.80 | 4.81 | $1.41 \times 10^{-1}$ | 2.92 |
| $\beta_{11}$ | $1.00 \times 10^{-1}$ | $1.00 \times 10^{1}$ | 6.00 | 6.01 | $6.02 \times 10^{-1}$ | $1.0 \times 10^{1}$ |
| $\beta_{21}$ | $1.00 \times 10^{-1}$ | $1.00 \times 10^{1}$ | 6.24 | 6.02 | $5.86 \times 10^{-1}$ | 9.74 |
| $\beta_{12}$ | $1.00 \times 10^{-1}$ | $1.00 \times 10^{1}$ | 5.40 | 5.45 | $3.30 \times 10^{1}$ | $6.06 \times 10^{2}$ |
| $\beta_{22}$ | $1.00 \times 10^{-1}$ | $1.00 \times 10^{1}$ | 4.80 | 9.74 | $3.52 \times 10^{1}$ | $3.61 \times 10^{2}$ |
| $X_{10}$ | $1.00 \times 10^{-2}$ | 1.00 | $5.00 \times 10^{-1}$ | $4.31 \times 10^{-1}$ | $7.89 \times 10^{-2}$ | $1.83 \times 10^{1}$ |
| $X_{20}$ | $1.00 \times 10^{-2}$ | 1.00 | $5.00 \times 10^{-1}$ | $4.80 \times 10^{-1}$ | $8.20 \times 10^{-2}$ | $1.71 \times 10^{1}$ |

Table I: Case study: ***GLV2_Comp***. Results for the *Generalized Lotka-Volterra 2 species* model with 10% heteroscedastic Gaussian noise. The table presents the lower and upper bounds (**LB**, **UB**), nominal and estimated values, as well as the absolute and relative errors (**EA**, **ER**), for each estimated parameter.

##### Case study: Coexistence

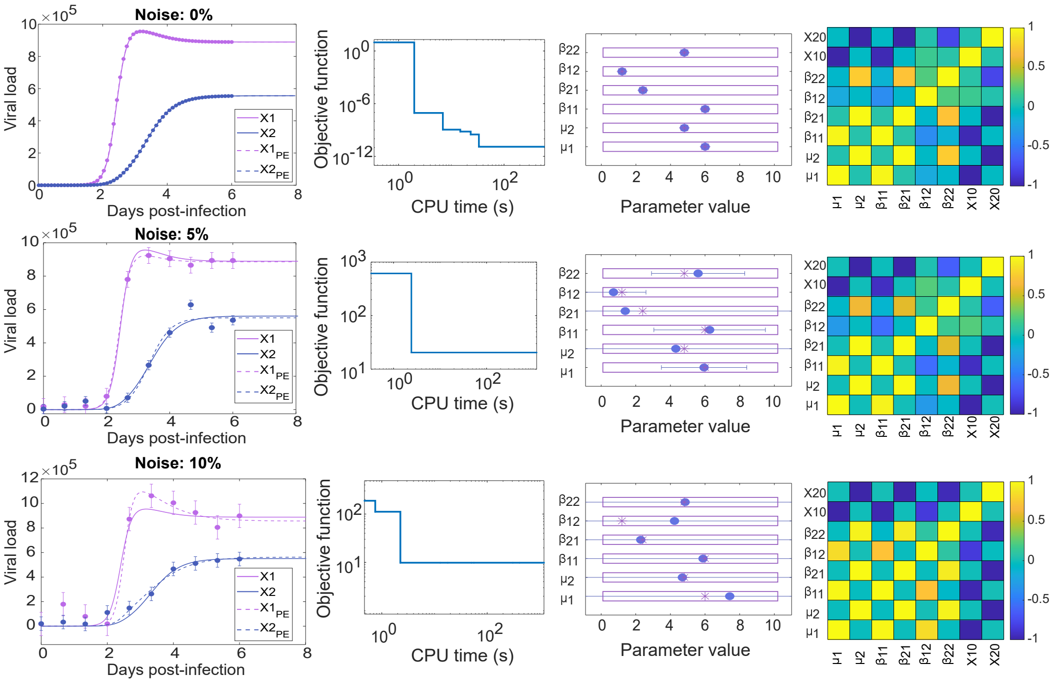

Figure D: Case study ***GLV2_Coex***: Model calibration results for the coexistence scenario of the *Generalized Lotka-Volterra* two species model (Dimas Martins and Gjini, 2020). The first row displays results for the pseudo-experimental data with 0% noise, the second row shows results for 5% noise, and the third row corresponds to 10% noise. In each case, the simulation using the estimated parameters is compared to the simulation with nominal parameters, along with the convergence curve, correlation matrix, and a comparison of nominal and estimated parameters. These results demonstrate the model's ability to accurately fit the data under varying noise levels.

After estimating the parameters, we conducted a sensitivity analysis to evaluate their influence on the population dynamics (Figure E). This involved calculating the mean relative sensitivity and perturbing individually each parameter by +5% to identify which ones have the greatest impact on species population sizes.

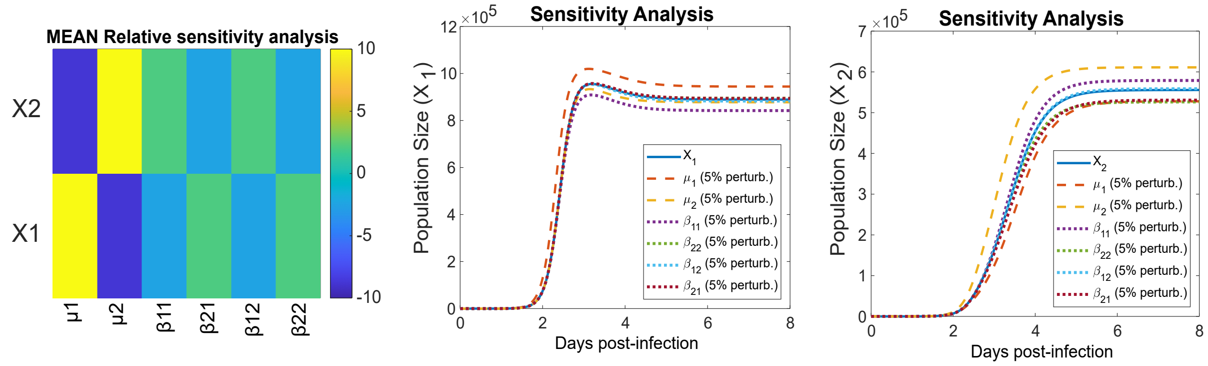

Figure E: Case study ***GLV2_Coex***: Sensitivity analysis for the coexistence scenario of a two species *Generalized Lotka-Volterra*. **Left**: Mean relative sensitivity analysis. **Center**: Simulations of species $X_1$ with a positive 5% perturbation of each parameter. **Right**: Simulations of species $X_2$ with a positive 5% perturbation of each parameter.

| | **LB** | **UB** | **Nominal** | **Estimated** | **EA** | **ER (%)** |
|---|---|---|---|---|---|---|
| $\mu_1$ | $1.00 \times 10^{-1}$ | $1.00 \times 10^{1}$ | 6.00 | 6.00 | $6.30 \times 10^{-6}$ | $2.48 \times 10^{-5}$ |
| $\mu_2$ | $1.00 \times 10^{-1}$ | $1.00 \times 10^{1}$ | 4.80 | 4.80 | $2.75 \times 10^{-5}$ | $4.78 \times 10^{-4}$ |
| $\beta_{11}$ | $1.00 \times 10^{-1}$ | $1.00 \times 10^{1}$ | 6.00 | 6.00 | $7.51 \times 10^{-6}$ | $3.12 \times 10^{-5}$ |
| $\beta_{21}$ | $1.00 \times 10^{-1}$ | $1.00 \times 10^{1}$ | 2.40 | 2.40 | $2.80 \times 10^{-5}$ | $1.05 \times 10^{-3}$ |
| $\beta_{12}$ | $1.00 \times 10^{-1}$ | $1.00 \times 10^{1}$ | 1.20 | 1.20 | $3.57 \times 10^{-6}$ | $5.52 \times 10^{-4}$ |
| $\beta_{22}$ | $1.00 \times 10^{-1}$ | $1.00 \times 10^{1}$ | 4.80 | 4.80 | $5.92 \times 10^{-6}$ | $5.04 \times 10^{-5}$ |
| $X_{10}$ | $1.00 \times 10^{-2}$ | 1.00 | $5.00 \times 10^{-1}$ | $5.00 \times 10^{-1}$ | $7.49 \times 10^{-6}$ | $3.54 \times 10^{-4}$ |
| $X_{20}$ | $1.00 \times 10^{-2}$ | 1.00 | $5.00 \times 10^{-1}$ | $5.00 \times 10^{-1}$ | $3.31 \times 10^{-5}$ | $5.63 \times 10^{-3}$ |

Table J: Case study ***GLV2_Coex***: Results for the *Generalized Lotka-Volterra 2 species* model with 0% homoscedastic Gaussian noise. The table represents the lower and upper bounds (**LB**, **UB**), nominal and estimated values, as well as the absolute and relative errors (**EA**, **ER**), for each estimated parameter.

| | **LB** | **UB** | **Nominal** | **Estimated** | **EA** | **ER (%)** |
|---|---|---|---|---|---|---|
| $\mu_1$ | $1.00 \times 10^{-1}$ | $1.00 \times 10^{1}$ | 6.00 | 5.94 | 2.46 | $9.25 \times 10^{-1}$ |
| $\mu_2$ | $1.00 \times 10^{-1}$ | $1.00 \times 10^{1}$ | 4.80 | 4.31 | $1.94 \times 10^{1}$ | $1.01 \times 10^{1}$ |
| $\beta_{11}$ | $1.00 \times 10^{-1}$ | $1.00 \times 10^{1}$ | 6.00 | 6.27 | 3.22 | 4.45 |
| $\beta_{21}$ | $1.00 \times 10^{-1}$ | $1.00 \times 10^{1}$ | 2.40 | 1.39 | $2.08 \times 10^{1}$ | $4.20 \times 10^{1}$ |
| $\beta_{12}$ | $1.00 \times 10^{-1}$ | $1.00 \times 10^{1}$ | 1.20 | $7.14 \times 10^{-1}$ | 1.88 | $4.05 \times 10^{1}$ |
| $\beta_{22}$ | $1.00 \times 10^{-1}$ | $1.00 \times 10^{1}$ | 4.80 | 5.60 | 2.69 | $1.67 \times 10^{1}$ |
| $X_{10}$ | $1.00 \times 10^{-2}$ | 1.00 | $5.0 \times 10^{-1}$ | $5.99 \times 10^{-1}$ | 3.58 | $1.98 \times 10^{1}$ |
| $X_{20}$ | $1.00 \times 10^{-2}$ | 1.00 | $5.0 \times 10^{-1}$ | 1.00 | $4.64 \times 10^{1}$ | $1.00 \times 10^{2}$ |

Table K: Case study ***GLV2_Coex***: Results for the *Generalized Lotka-Volterra 2 species* model with 5% homoscedastic Gaussian noise. The table represents the lower and upper bounds (**LB**, **UB**), nominal and estimated values, as well as the absolute and relative errors (**EA**, **ER**), for each estimated parameter.

| | **LB** | **UB** | **Nominal** | **Estimated** | **EA** | **ER (%)** |
|---|---|---|---|---|---|---|
| $\mu_1$ | $1.00 \times 10^{-1}$ | $1.00 \times 10^{1}$ | 6.00 | 7.43 | $1.07 \times 10^{1}$ | $-2.38 \times 10^{1}$ |
| $\mu_2$ | $1.00 \times 10^{-1}$ | $1.00 \times 10^{1}$ | 4.80 | 4.69 | $1.52 \times 10^{1}$ | $2.28 \times 10^{-2}$ |
| $\beta_{11}$ | $1.00 \times 10^{-1}$ | $1.00 \times 10^{1}$ | 6.00 | 5.88 | 8.76 | $1.92 \times 10^{-2}$ |
| $\beta_{21}$ | $1.00 \times 10^{-1}$ | $1.00 \times 10^{1}$ | 2.40 | 2.29 | $1.22 \times 10^{1}$ | $4.69 \times 10^{-2}$ |
| $\beta_{12}$ | $1.00 \times 10^{-1}$ | $1.00 \times 10^{1}$ | 1.20 | 4.24 | 6.93 | $-2.53$ |
| $\beta_{22}$ | $1.00 \times 10^{-1}$ | $1.00 \times 10^{1}$ | 4.80 | 4.85 | 8.76 | $-1.05 \times 10^{-2}$ |
| $X_{10}$ | $1.00 \times 10^{-2}$ | 1.00 | $5.0 \times 10^{-1}$ | $1.00 \times 10^{-2}$ | $2.82 \times 10^{-1}$ | $-9.80 \times 10^{1}$ |
| $X_{20}$ | $1.00 \times 10^{-2}$ | 1.00 | $5.0 \times 10^{-1}$ | 1.00 | $3.83 \times 10^{1}$ | $1.00 \times 10^{2}$ |

Table L: Case study: ***GLV2_Coex***. Results for the *Generalized Lotka-Volterra 2 species* model with 10% homoscedastic Gaussian noise. The table represents the lower and upper bounds (**LB**, **UB**), nominal and estimated values, as well as the absolute and relative errors (**EA**, **ER**), for each estimated parameter.

##### 6.5.4 Estimation using multistart of local methods

In order to illustrate the advantages of using a global optimizer (`eSS`) versus a multistart with local methods, we present below the results obtained for this case study. The histograms illustrate the distribution of objective function values obtained from multiple runs, allowing for a comparison between the robustness and consistency of both approaches.

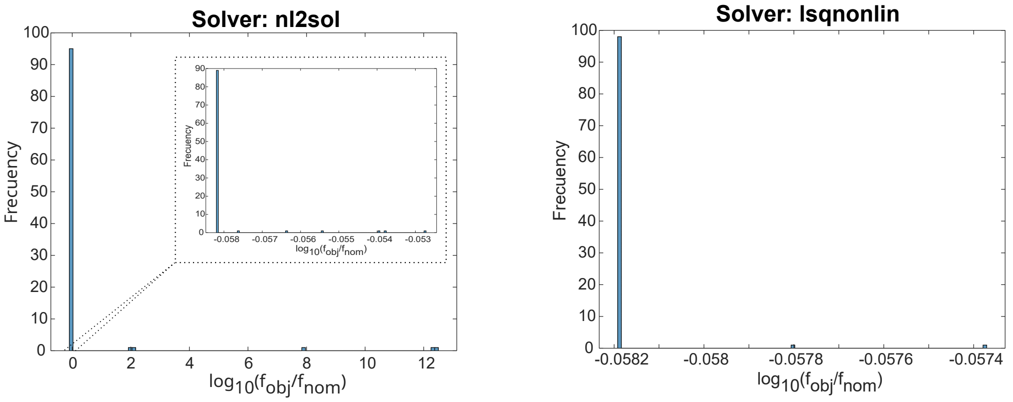

Figure F: Case study ***GLV2_Comp***: Histogram of objective function values obtained from 100 optimization runs using the local solvers `nl2sol` (**Left**) and `lsqnonlin` (**Right**). All runs successfully converged. For `nl2sol`, 95 out of 100 runs resulted in a logarithm of the normalized objective function value ($log_{10}(f_{obj}/f_{nom})$) below 0, whereas 5 runs produced higher values. For `lsqnonlin`, 100 out of 100 runs achieved values below 0.

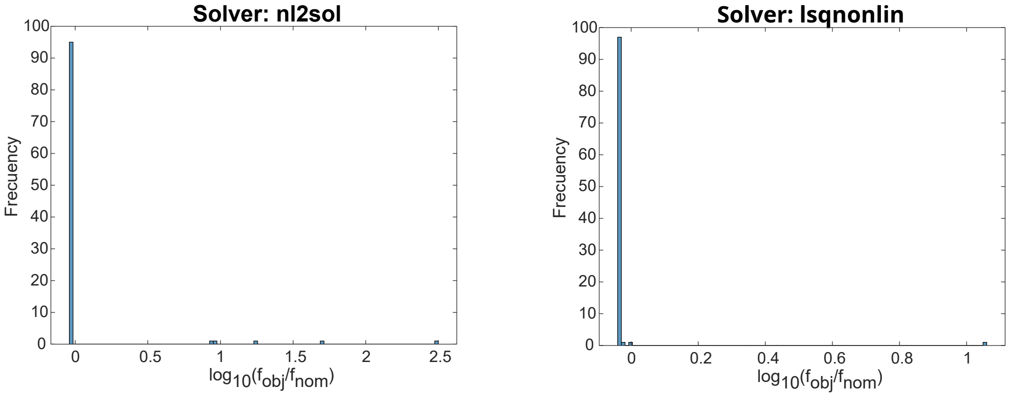

Figure G: Case study ***GLV2_Coex***: Histogram of objective function values obtained from 100 optimization runs using the local solvers `nl2sol` (**Left**) and `lsqnonlin` (**Right**). For `nl2sol`, 95 out of 100 runs resulted in a logarithm of the normalized objective function value ($log_{10}(f_{obj}/f_{nom})$) below 0, whereas 5 runs produced higher values. For `lsqnonlin`, 98 out of 100 runs achieved values below 0, with 2 runs yielding higher outcomes.

#### 6.6 *GLV3*: *Generalized Lotka-Volterra* 3 species

##### 6.6.1 Model

Assuming a three-species population with only two-way interactions between microbes (as in Figure H), the change in density over time of the species populations can be described by the following system:

$$\frac{dX_1}{dt} = \mu_1 X_1 + \beta_{11} X_1^2 + \beta_{12} X_1 X_2 + \beta_{13} X_1 X_3 \quad (18)$$

$$\frac{dX_2}{dt} = \mu_2 X_2 + \beta_{21} X_1 X_2 + \beta_{22} X_2^2 + \beta_{23} X_2 X_3 \quad (19)$$

$$\frac{dX_3}{dt} = \mu_3 X_3 + \beta_{31} X_1 X_3 + \beta_{32} X_2 X_3 + \beta_{33} X_3^2 \quad (20)$$

$N$ represents the total abundance of microbes in the community: $\frac{dN}{dt} = \sum_{i=1}^{N} \frac{dX_i}{dt}$.

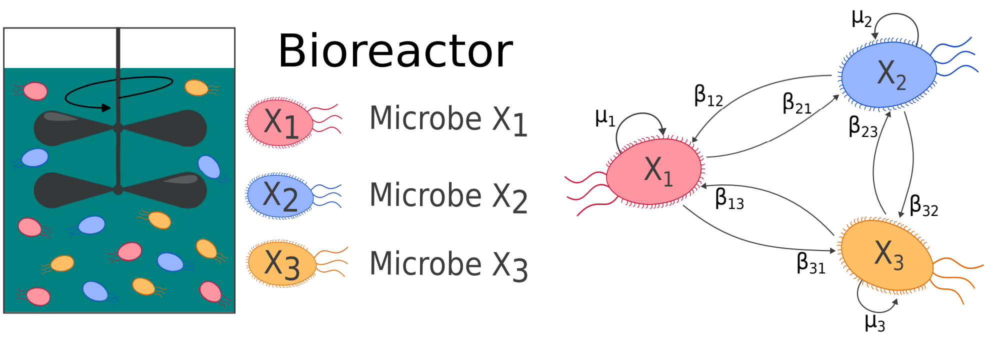

Figure H: A bioreactor with three microbe populations ($X_1$, $X_2$, $X_3$) (**Left**). Interactions among three microorganisms (**Right**).

The parameter values, as presented in Table M, were obtained from Remien *et al.* (2021).

| Parameter | Value [AU] | Parameter | Value [AU] | Parameter | Value [AU] |
|---|---|---|---|---|---|
| $\mu_1$ | 6.00 | $\beta_{1,1}$ | $-5.00 \times 10^{-2}$ | $\beta_{2,1}$ | $-1.00 \times 10^{-2}$ |
| $\mu_2$ | 4.00 | $\beta_{1,2}$ | $1.50 \times 10^{-1}$ | $\beta_{2,2}$ | $-2.60 \times 10^{-2}$ |
| $\mu_3$ | 2.00 | $\beta_{1,3}$ | $-2.00 \times 10^{-1}$ | $\beta_{2,3}$ | $5.00 \times 10^{-2}$ |
| **Initial conditions (IC):** $X_{10} = 10.00$, $X_{20} = 14.00$, $X_{30} = 4.00$ | | | | | |

Table M: Case study ***GLV3***: Parameter values and initial conditions (IC) for the case study of the *Generalized Lotka-Volterra* three-species model, as obtained from Remien *et al.* (2021). The parameters ($\mu_i$, $\beta_{i,j}$, $\beta_{i,3}$) represent the growth and interaction rates, while the initial conditions ($X_{i0}$) denote the starting population densities.

##### 6.6.2 Structural Identifiability Analysis (SIA)

We analyzed three observation scenarios: one fully observed ($X_1$,$X_2$,$X_3$) and two partially observed ($X_1$+$X_2$, $X_3$; $X_1$,$X_2$). As with the ***GLV2*** model, `GenSSI2` has the longest execution times (Table N). It provided locally identifiable results for parameters in the partially observed scenarios, which the other tools classify as globally identifiable. This difference arises from how initial conditions are defined in `GenSSI2`. It can not include arithmetic operations in the initial conditions, whereas `SIAN` assumes that the initial conditions match the observed scenarios. For example, if the observed scenario is $X_1 + X_2, X_3$, `SIAN` defines the initial conditions as $X_{10} + X_{20}, X_{30}$, but `GenSSI2` does not allow for addition in the initial conditions.

Among the three tools, `Structural Identifiability` is the most flexible regarding initial conditions, as these are defined by the user. For our analysis, we used the observed states at time zero as initial conditions, similar to `SIAN`.

| | `GenSSI2` (**IC Unknown**) | | | | `GenSSI2` (**IC Known**) | | | |
|---|---|---|---|---|---|---|---|---|
| Obs. | Global | Local | Non-Id | Time[s] | Global | Local | Non-Id | Time[s] |
| $X_1$,$X_2$,$X_3$ | $X_{i0}$, $\mu_i$, $\beta_{ii}$, $\beta_{ij}$ | - | - | 20.10 | $\mu_i$, $\beta_{ii}$, $\beta_{ij}$ | - | - | 11.16 |
| $X_1 + X_2$,$X_3$ | $X_{30}$ | $\beta_{ii}$,$\beta_{ij}$, $\mu_i$, $X_{10}, X_{20}$ | - | 1826.44 | - | $\mu_i$, $\beta_{ii}$, $\beta_{ij}$ | - | 1653.39 |
| $X_1$,$X_2$ | - | - | - | - | - | $\mu_i, \beta_{ii}, \beta_{ij}$ | - | 20551.30 |

Table N: Case study ***GLV3***: Structural identifiability analysis (SIA) results obtained with `GenSSI2` for the *Generalized Lotka-Volterra* 3 species model. The analysis was performed considering three observation scenarios ($X_1$,$X_2$,$X_3$; $X_1 + X_2$, $X_3$; $X_1$, $X_2$) and unknown initial conditions (IC Unknown). The parameters are classified as globally identifiable ("Global"), locally identifiable ("Local"), or non-identifiable ("Non-Id"). A " - " indicates that no parameters are found in that category. The table summarizes the computational time and the identifiability results for the model parameters ($\mu_i$, $\beta_{ii}$, $\beta_{ij}$).

| **IC Unknown** | `SIAN` | | | | `Structural Identifiability` | | | |
|---|---|---|---|---|---|---|---|---|
| Obs. | Global | Local | Non-Id | Time[s] | Global | Local | Non-Id | Time[s] |
| $X_1$,$X_2$,$X_3$ | $X_i$,$\mu_i$,$\beta_{ii}$,$\beta_{ij}$ | - | - | 56.48 | $X_i(t)$,$\mu_i$,$\beta_{ii}$,$\beta_{ij}$ | - | - | 19.62 |
| $X_1+X_2$,$X_3$ | $X_3$, $\beta_{33}$,$\mu_3$ | $X_1$, $X_2$ $\beta_{11}$,$\beta_{12}$,$\beta_{13}$, $\beta_{21}$,$\beta_{22}$,$\beta_{23}$, $\beta_{31}$,$\beta_{32}$,$\mu_1$,$\mu_2$ | - | 61.74 | $X_3(t)$, $b_{33}$, $\mu_3$ | $X_1(t)$, $X_2(t)$, $\beta_{11}$,$\beta_{12}$,$\beta_{13}$, $\beta_{21}$,$\beta_{22}$,$\beta_{23}$, $\beta_{31}$,$\beta_{32}$,$\mu_1$,$\mu_2$ | - | 22.29 |
| $X_1$,$X_2$ | $X_1$, $X_2$ $\beta_{11}$, $\beta_{12}$,$\beta_{21}$,$\beta_{22}$, $\beta_{31}$,$\beta_{32}$,$\mu_i$ | - | $X_3$,$\beta_{13}$, $\beta_{23}$,$\beta_{33}$ | 58.95 | $X_1(t)$, $X_2(t)$, $\beta_{11}$,$\beta_{12}$,$\beta_{21}$, $\beta_{22}$,$\beta_{31}$,$\beta_{32}$,$\mu_i$ | - | $X_3(t)$,$\beta_{13}$, $\beta_{23}$,$\beta_{33}$ | 20.61 |

Table O: Case study ***GLV3***: Structural identifiability analysis (SIA) results obtained with `SIAN` and `Structural Identifiability` for the *Generalized Lotka-Volterra* 3 species model. The analysis was performed considering three observation scenarios ($X_1$,$X_2$,$X_3$; $X_1+X_2$, $X_3$; $X_1, X_2$) and unknown initial conditions (IC Unknown). The parameters are classified as globally identifiable ("Global"), locally identifiable ("Local"), or non-identifiable ("Non-Id"). A " - " indicates that no parameters are found in that category. The table summarizes the computational time and the identifiability results for the model parameters ($\mu_i$, $\beta_{ii}$, $\beta_{ij}$).

| **IC Known** | `SIAN` | | | | `Structural Identifiability` | | | |
|---|---|---|---|---|---|---|---|---|
| Obs. | Global | Local | Non-Id | Time[s] | Global | Local | Non-Id | Time[s] |
| $X_1$,$X_2$,$X_3$ | $\mu_i$, $\beta_{ii}$, $\beta_{ij}$ | - | - | 56.61 | $X_{i0}$, $\mu_i$, $\beta_{ii}$, $\beta_{ij}$ | - | - | 26.77 |
| $X_1+X_2$,$X_3$ | $\beta_{33}$,$\mu_3$ | $\beta_{11}$,$\beta_{12}$,$\beta_{13}$, $\beta_{21}$,$\beta_{22}$,$\beta_{23}$, $\beta_{31}$,$\beta_{32}$,$\mu_1$,$\mu_2$ | - | 58.69 | $X_{30}$, $\beta_{33}$,$\mu_3$ | $X_{10}$,$X_{20}$, $\beta_{11}$,$\beta_{12}$,$\beta_{13}$, $\beta_{21}$,$\beta_{22}$,$\beta_{23}$, $\beta_{31}$,$\beta_{32}$,$\mu_1$,$\mu_2$ | - | 33.94 |
| $X_1$,$X_2$ | $\beta_{11}$,$\beta_{12}$, $\beta_{21}$,$\beta_{22}$, $\beta_{31}$,$\beta_{32}$,$\mu_i$ | - | $\beta_{13}$, $\beta_{23}$,$\beta_{33}$ | 59.15 | $X_{10}$,$X_{20}$,$\mu_i$, $\beta_{11}$,$\beta_{12}$,$\beta_{21}$, $\beta_{22}$,$\beta_{31}$,$\beta_{32}$ | - | $X_{30}$,$\beta_{13}$, $\beta_{23}$,$\beta_{33}$ | 26.96 |

Table P: Case study ***GLV3***: Structural identifiability analysis (SIA) results obtained with `SIAN` and `Structural Identifiability` for the *Generalized Lotka-Volterra* 3 species model. The analysis was performed considering three observation scenarios ($X_1$,$X_2$,$X_3$; $X_1$+$X_2$, $X_3$; $X_1, X_2$) and known initial conditions (IC Known). The parameters are classified as globally identifiable ("Global"), locally identifiable ("Local"), or non-identifiable ("Non-Id"). A " - " indicates that no parameters are found in that category. The table summarizes the computational time and the identifiability results for the model parameters ($\mu_i$, $\beta_{ii}$, $\beta_{ij}$).

##### 6.6.3 Practical Identifiability Analysis (PIA)

We addressed a parameter estimation problem involving three observed populations, $X_1$, $X_2$, $X_3$, with unknown initial conditions ($X_{10}$, $X_{20}$, $X_{30}$) (Figure I). To assess the performance and robustness of our estimation methods, we generated three sets of pseudo-experimental data, each incorporating different levels of Gaussian noise: 0%, 5%, and 10%. The generated data was homoscedastic and positive, reflecting the nature of population measurements.

Lotka-Volterra systems are inherently unstable, making them prone to overfitting, especially as the noise level increases (see Figure I for the 5% and 10% Gaussian noise cases). In such cases, the parameters are overly adjusted, resulting in an objective function value lower than its nominal value. This overfitting can lead to inaccurate representations of the underlying dynamics, as the model becomes too closely tailored to the noise rather than the true system behavior, resulting in erroneous parameter estimates. For instance, with 5% noise, the parameters $\beta_{22}$ and $\beta_{33}$ change sign (Table R), and with 10% noise, $\beta_{33}$ changes its sign relative to the nominal value (Table S). These changes in parameter signs can lead to misleading interpretations of the system dynamics and mechanistic errors in understanding the relationships between the species or the underlying processes.

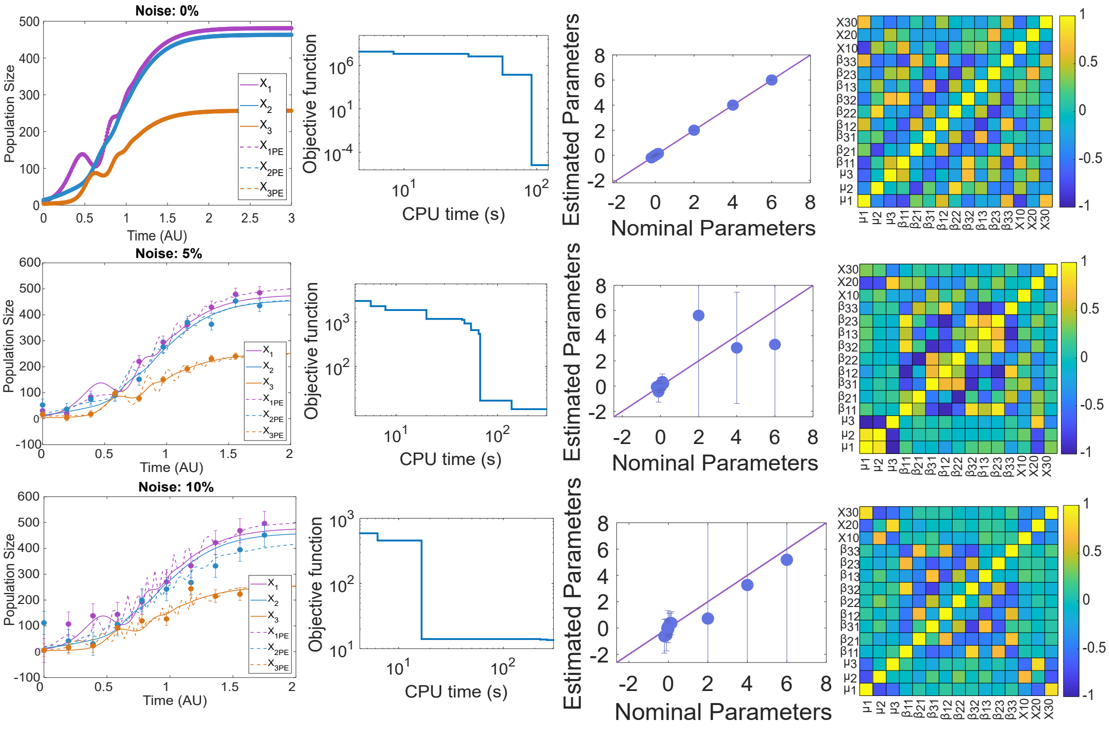

Figure I: Case study ***GLV3***: Model calibration results for the *Generalized Lotka-Volterra* model of three species (Remien *et al.*, 2021). The first row displays results for the simulations with pseudo-experimental data with 0% noise, the second row shows results for 5% noise, and the third row corresponds to 10% noise. In each case, the simulation using the estimated parameters is compared to the simulation with nominal parameters, along with the convergence curve, correlation matrix, and a comparison of nominal and estimated parameters.

| | **LB** | **UB** | **Nominal** | **Estimated** | **EA** | **ER (%)** |
|---|---|---|---|---|---|---|
| $\mu_1$ | $1.00 \times 10^{-1}$ | $1.00 \times 10^{1}$ | 6.00 | 6.00 | $6.01 \times 10^{-6}$ | $6.66 \times 10^{-5}$ |
| $\mu_2$ | $1.00 \times 10^{-1}$ | $1.00 \times 10^{1}$ | 4.00 | 4.00 | $1.99 \times 10^{-6}$ | $9.51 \times 10^{-5}$ |
| $\mu_3$ | $1.00 \times 10^{-1}$ | $1.00 \times 10^{1}$ | 2.00 | 2.00 | $4.99 \times 10^{-6}$ | $5.41 \times 10^{-4}$ |
| $\beta_{11}$ | $-5.00 \times 10^{-1}$ | $5.00 \times 10^{-1}$ | $-5.00 \times 10^{-2}$ | $-5.00 \times 10^{-2}$ | $1.15 \times 10^{-7}$ | $1.85 \times 10^{-4}$ |
| $\beta_{21}$ | $-1.00 \times 10^{-1}$ | $1.00 \times 10^{-1}$ | $-1.00 \times 10^{-2}$ | $-1.00 \times 10^{-2}$ | $6.29 \times 10^{-8}$ | $4.64 \times 10^{-4}$ |
| $\beta_{31}$ | $-1.00$ | 1.00 | $1.00 \times 10^{-1}$ | $1.00 \times 10^{-1}$ | $1.32 \times 10^{-7}$ | $1.21 \times 10^{-4}$ |
| $\beta_{12}$ | $-1.50$ | 1.50 | $1.50 \times 10^{-1}$ | $1.50 \times 10^{-1}$ | $1.90 \times 10^{-7}$ | $3.62 \times 10^{-5}$ |
| $\beta_{22}$ | $-3.00 \times 10^{-1}$ | $3.00 \times 10^{-1}$ | $-2.60 \times 10^{-2}$ | $-2.60 \times 10^{-2}$ | $1.02 \times 10^{-7}$ | $8.23 \times 10^{-4}$ |
| $\beta_{32}$ | $-1.00$ | 1.00 | $-1.00 \times 10^{-1}$ | $-1.00 \times 10^{-1}$ | $1.87 \times 10^{-7}$ | $3.29 \times 10^{-4}$ |
| $\beta_{13}$ | $-1.00$ | 1.00 | $-2.00 \times 10^{-1}$ | $-2.00 \times 10^{-1}$ | $2.60 \times 10^{-7}$ | $2.86 \times 10^{-5}$ |
| $\beta_{23}$ | $-5.00 \times 10^{-1}$ | $5.00 \times 10^{-1}$ | $5.00 \times 10^{-2}$ | $5.00 \times 10^{-2}$ | $1.48 \times 10^{-7}$ | $6.28 \times 10^{-4}$ |
| $\beta_{33}$ | $-1.00 \times 10^{-1}$ | $1.00 \times 10^{-1}$ | $-1.48 \times 10^{-2}$ | $-1.48 \times 10^{-2}$ | $2.46 \times 10^{-7}$ | $5.83 \times 10^{-3}$ |
| $X_1$ | 0 | $2.00 \times 10^{1}$ | 10.00 | 10.00 | $1.44 \times 10^{-5}$ | $2.05 \times 10^{-4}$ |
| $X_2$ | 0 | $2.00 \times 10^{1}$ | 14.00 | 14.00 | $1.07 \times 10^{-5}$ | $1.94 \times 10^{-4}$ |
| $X_3$ | 0 | $1.00 \times 10^{1}$ | 4.00 | 4.00 | $9.43 \times 10^{-6}$ | $6.16 \times 10^{-4}$ |

Table Q: Case study ***GLV3***: Results for the *Generalized Lotka-Volterra* 3 species model with 0% homoscedastic Gaussian noise. The table represents the lower and upper bounds (**LB**, **UB**), nominal and estimated values, as well as the absolute and relative errors (**LB**, **UB**), for each estimated parameter.

| | LB | UB | Nominal | Estimated | EA | ER (%) |
|---|---|---|---|---|---|---|
| $\mu_1$ | $1.00 \times 10^{-1}$ | $1.00 \times 10^{1}$ | 6.00 | 3.30 | 6.39 | $4.50 \times 10^{1}$ |
| $\mu_2$ | $1.00 \times 10^{-1}$ | $1.00 \times 10^{1}$ | 4.00 | 3.04 | 4.42 | $2.41 \times 10^{1}$ |
| $\mu_3$ | $1.00 \times 10^{-1}$ | $1.00 \times 10^{1}$ | 2.00 | 5.61 | $1.51 \times 10^{1}$ | $1.80 \times 10^{2}$ |
| $\beta_{11}$ | $-5.00 \times 10^{-1}$ | $5.00 \times 10^{-1}$ | $-5.00 \times 10^{-2}$ | $-1.56 \times 10^{-1}$ | $3.67 \times 10^{-1}$ | $2.12 \times 10^{2}$ |
| $\beta_{21}$ | $-1.00 \times 10^{-1}$ | $1.00 \times 10^{-1}$ | $-1.00 \times 10^{-2}$ | $-9.90 \times 10^{-2}$ | $2.87 \times 10^{-1}$ | $8.90 \times 10^{2}$ |
| $\beta_{31}$ | $-1.00$ | 1.00 | $1.00 \times 10^{-1}$ | $3.46 \times 10^{-1}$ | $6.15 \times 10^{-1}$ | $2.46 \times 10^{2}$ |
| $\beta_{12}$ | $-1.50$ | 1.50 | $1.50 \times 10^{-1}$ | $1.98 \times 10^{-1}$ | $4.97 \times 10^{-1}$ | $3.18 \times 10^{1}$ |
| $\beta_{22}$ | $-3.00 \times 10^{-1}$ | $3.00 \times 10^{-1}$ | $-2.60 \times 10^{-2}$ | $8.19 \times 10^{-2}$ | $3.44 \times 10^{-1}$ | $4.15 \times 10^{2}$ |
| $\beta_{32}$ | $-1.00$ | 1.00 | $-1.00 \times 10^{-1}$ | $-4.46 \times 10^{-1}$ | $8.30 \times 10^{-1}$ | $3.46 \times 10^{2}$ |
| $\beta_{13}$ | $-1.00$ | 1.00 | $-2.00 \times 10^{-1}$ | $-6.22 \times 10^{-2}$ | $2.73 \times 10^{-1}$ | $6.89 \times 10^{1}$ |
| $\beta_{23}$ | $-5.00 \times 10^{-1}$ | $5.00 \times 10^{-1}$ | $5.00 \times 10^{-2}$ | $3.65 \times 10^{-2}$ | $1.69 \times 10^{-1}$ | $2.70 \times 10^{1}$ |
| $\beta_{33}$ | $-1.00 \times 10^{-1}$ | $1.00 \times 10^{-1}$ | $-1.48 \times 10^{-2}$ | $1.00 \times 10^{-1}$ | $5.82 \times 10^{-1}$ | $7.76 \times 10^{2}$ |
| $X_1$ | 0 | $2.00 \times 10^{1}$ | 10.00 | $1.29 \times 10^{1}$ | $1.16 \times 10^{1}$ | $2.89 \times 10^{1}$ |
| $X_2$ | 0 | $2.00 \times 10^{1}$ | 14.00 | $2.00 \times 10^{1}$ | $2.04 \times 10^{1}$ | $4.29 \times 10^{1}$ |
| $X_3$ | 0 | $1.00 \times 10^{1}$ | 4.00 | 9.40 | $1.74 \times 10^{1}$ | $1.35 \times 10^{2}$ |

Table R: Case study ***GLV3***: Results for the *Generalized Lotka-Volterra* 3 species model with 5% homoscedastic Gaussian noise. The table represents the lower and upper bounds (**LB**, **UB**), nominal and estimated values, as well as the absolute and relative errors (**LB**, **UB**), for each estimated parameter. Values in blue indicate a sign change with respect to the nominal value.

| | LB | UB | Nominal | Estimated | EA | ER (%) |
|---|---|---|---|---|---|---|
| $\mu_1$ | $1.00 \times 10^{-1}$ | $1.00 \times 10^{1}$ | 6.00 | 5.19 | $2.30 \times 10^{1}$ | $1.35 \times 10^{1}$ |
| $\mu_2$ | $1.00 \times 10^{-1}$ | $1.00 \times 10^{1}$ | 4.00 | 3.27 | 7.48 | $1.82 \times 10^{1}$ |
| $\mu_3$ | $1.00 \times 10^{-1}$ | $1.00 \times 10^{1}$ | 2.00 | $7.23 \times 10^{-1}$ | $3.50 \times 10^{1}$ | $6.39 \times 10^{1}$ |
| $\beta_{11}$ | $-5.00 \times 10^{-1}$ | $5.00 \times 10^{-1}$ | $-5.00 \times 10^{-2}$ | $-7.04 \times 10^{-3}$ | $5.50 \times 10^{-1}$ | $8.59 \times 10^{1}$ |
| $\beta_{21}$ | $-1.00 \times 10^{-1}$ | $1.00 \times 10^{-1}$ | $-1.00 \times 10^{-2}$ | $-5.82 \times 10^{-2}$ | $4.29 \times 10^{-1}$ | $4.82 \times 10^{2}$ |
| $\beta_{31}$ | $-1.00$ | 1.00 | $1.00 \times 10^{-1}$ | $3.80 \times 10^{-1}$ | $7.47 \times 10^{-1}$ | $2.80 \times 10^{2}$ |
| $\beta_{12}$ | $-1.50$ | 1.50 | $1.50 \times 10^{-1}$ | $3.76 \times 10^{-1}$ | $8.62 \times 10^{-1}$ | $1.50 \times 10^{2}$ |
| $\beta_{22}$ | $-3.00 \times 10^{-1}$ | $3.00 \times 10^{-1}$ | $-2.60 \times 10^{-2}$ | $-7.98 \times 10^{-2}$ | $6.05 \times 10^{-1}$ | $2.07 \times 10^{2}$ |
| $\beta_{32}$ | $-1.00$ | 1.00 | $-1.00 \times 10^{-1}$ | $-5.22 \times 10^{-1}$ | 1.22 | $4.22 \times 10^{2}$ |
| $\beta_{13}$ | $-1.00$ | 1.00 | $-2.00 \times 10^{-1}$ | $-6.20 \times 10^{-1}$ | 1.28 | $2.10 \times 10^{2}$ |
| $\beta_{23}$ | $-5.00 \times 10^{-1}$ | $5.00 \times 10^{-1}$ | $5.00 \times 10^{-2}$ | $2.33 \times 10^{-1}$ | $9.02 \times 10^{-1}$ | $3.66 \times 10^{2}$ |
| $\beta_{33}$ | $-1.00 \times 10^{-1}$ | $1.00 \times 10^{-1}$ | $-1.48 \times 10^{-2}$ | $1.00 \times 10^{-1}$ | 1.21 | $7.76 \times 10^{2}$ |
| $X_1$ | 0 | $2.00 \times 10^{1}$ | 10.00 | $1.76 \times 10^{1}$ | $3.67 \times 10^{1}$ | $7.56 \times 10^{1}$ |
| $X_2$ | 0 | $2.00 \times 10^{1}$ | 14.00 | $2.00 \times 10^{1}$ | $3.12 \times 10^{1}$ | $4.29 \times 10^{1}$ |
| $X_3$ | 0 | $1.00 \times 10^{1}$ | 4.00 | $1.00 \times 10^{1}$ | $4.02 \times 10^{1}$ | $1.50 \times 10^{2}$ |

Table S: Case study ***GLV3***: Results for the *Generalized Lotka-Volterra* 3 species model with 10% homoscedastic Gaussian noise. The table displays the lower and upper bounds (**LB**, **UB**), nominal and estimated values, as well as the absolute and relative errors (**LB**, **UB**), for each estimated parameter. Values in blue indicate a sign change with respect to the nominal value.

##### 6.6.4 Optimization performance and challenges

We performed two multistart optimization runs, each with 100 iterations, using different local solvers: `nl2sol` and `lsqnonlin`. Both solvers resulted in a significant number of overfitting and underfitting instances (see the histograms in Figure J, and examples in Figure K.

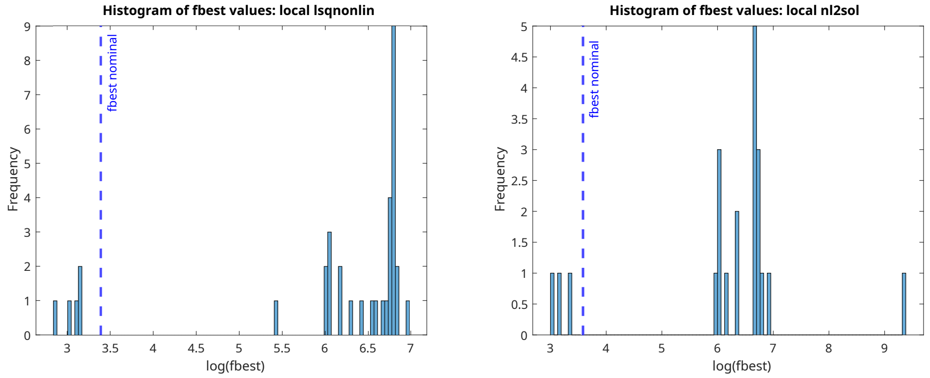

Figure J: Case study ***GLV3***: Histograms of two multistarts with 100 runs. **Left**: `lsqnonlin` local solver. **Rigth**: `nl2sol` local solver.

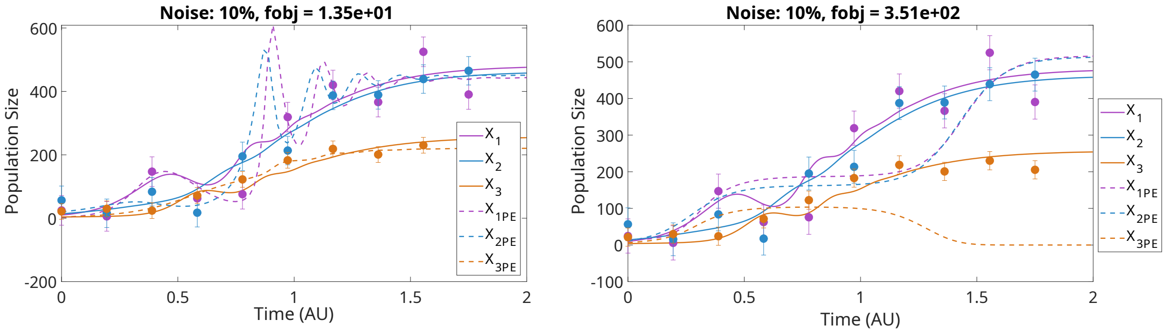

Figure K: Examples of overfitting (**Left**) and underfitting (**Right**) in a ***GLV3*** model using pseudo-experimental data with 10% Gaussian noise.

To illustrate the behavior of these solvers, we selected examples of underfitting and overfitting for further analysis (Figure L), highlighting the challenges and variability encountered during the simulations.

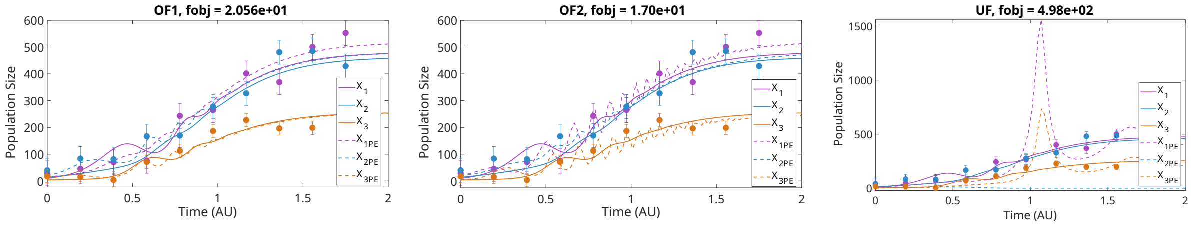

Figure L: Case study ***GLV3***: Examples of overfitting and underfitting using pseudo-experimental data with 10% Gaussian noise with the `nl2sol` solver. The objective function for the nominal values is: 29.691.

A post-analysis was performed on the results obtained during the different multistarts to evaluate the fitting quality. The metrics included the best objective function value ($f_{best}$), root mean squared error ($RMSE$), and normalized root mean squared error ($NRMSE$). The results for the overfittings and underfitting of Figure L are displayed in Table T.

| Metric | OF1 | OF2 | UF |
|---|---|---|---|
| $f_{best}$ | $2.0564 \times 10^1$ | $1.6971 \times 10^1$ | $4.9782 \times 10^2$ |
| $RMSE$ | [37.4739, 34.6127, 22.5524] | [34.3431, 38.0883, 16.4738] | [78.1152, 300.0223, 37.9677] |
| $NRMSE$ | [0.0715, 0.07774, 0.088] | [0.0655, 0.0856, 0.0649] | [0.1491, 0.6739, 0.1495] |

Table T: Case study ***GLV3***: Post-analysis metrics comparing the overfittings (**OF1**, **OF2**) and underfitting (**UF**) scenarios. Results include the best objective function value ($f_{best}$), root-mean-square error ($RMSE$), and normalized root-mean-square error ($NRMSE$) for the scenarios. Refer to Figure L for a graphical illustration.

The previous examples of overfitting and underfitting were chosen to highlight extreme cases and provide a clearer understanding of the solver's behavior. Now, we present a multistart of 200 runs with the `nl2sol` local solver (Figure M), with a more subtle case of overfitting, where, without knowledge of the nominal parameter values, the model fit might appear accurate (Figure N).

In Table U, we present a post-analysis comparing the different fits, and in Table V, we present the parameter estimation results, showing how the change in sign increases with greater overfitting (i.e., lower values of the cost function).

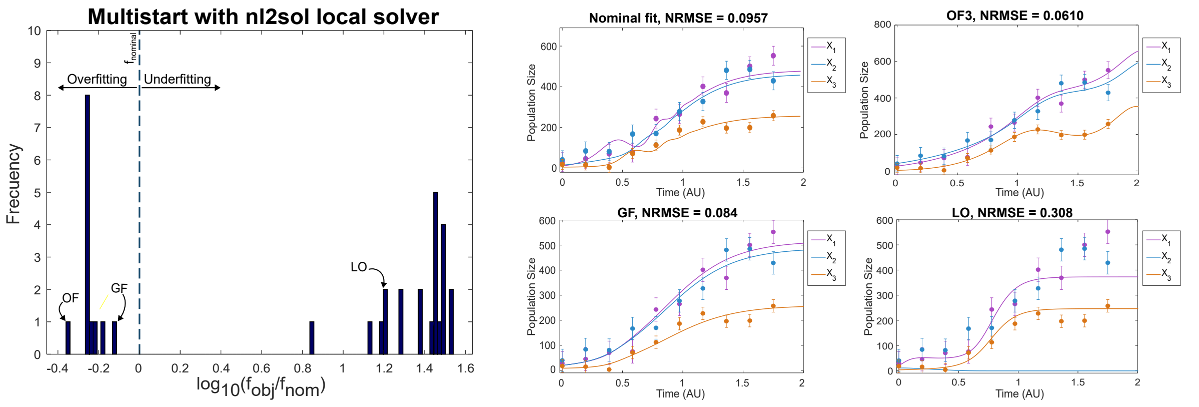

Figure M: Case study ***GLV3***: Multistart optimization using the `nl2sol` local solver. Histogram of the 35 optima obtained in 200 runs, with examples of overfitting (OF), underfitting (local optimum, LO), and a good fit (GF) are presented. 13 out of 200 runs resulted in a logarithm of the normalized objective function value ($log10(fobj/fnom)$) below 0, whereas 22 runs produced higher values. The $x$-axis shows the $\log_{10}$ of the ratio between the objective function achieved and that of the nominal vector of parameters.

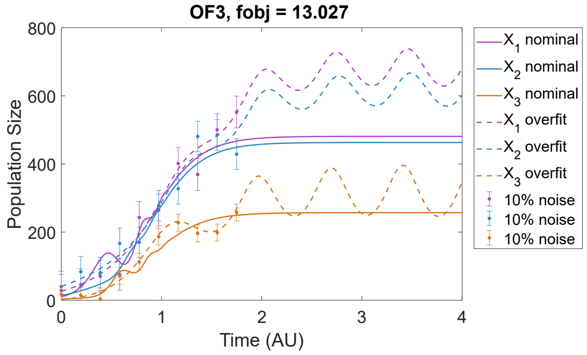

Figure N: Case study ***GLV3***: Example of overfitting (dashed lines) compared with the fit using nominal parameter values (solid line).

| **Metric** | **Nominal** | **GF** | **OF3** | **LO** |
|---|---|---|---|---|
| $f_{best}$ | $2.9691 \times 10^1$ | $2.2885 \times 10^1$ | $1.3027 \times 10^1$ | $4.8593 \times 10^2$ |
| $RMSE$ | [44.3006, 45.2168, 25.6109] | [39.3667, 37.6293, 23.3290] | [34.3713, 34.8123, 9.9659] | [75.4108, 300.6382, 26.5893] |
| $NRMSE$ | [0.0846, 0.1016, 0.1009] | [0.0751, 0.0845, 0.09188] | [0.0656, 0.0782, 0.0393] | [0.1439, 0.6753, 0.1047] |

Table U: Case study ***GLV3***: Post-analysis metrics comparing the overfitting (**OF3**), underfitting (**LO**), and good fit (**GF**) scenarios with the nominal fit. Results include the best objective function value ($f_{best}$), root-mean-square error ($RMSE$), and normalized root-mean-square error ($NRMSE$) for the three scenarios. Refer to Figure M for a graphical illustration.

| | LB | UB | Nominal | Overfitting 1 | | Overfitting 2 | | Overfitting 3 | | Good Fit | |
|---|---|---|---|---|---|---|---|---|---|---|---|
| | | | | Estimated | EA | Estimated | EA | Estimated | EA | Estimated | EA |
| $\mu_1$ | $1 \times 10^{-1}$ | $1.0 \times 10^{1}$ | 6.00 | 2.42 | $1.41 \times 10^{1}$ | 3.67 | $1.88 \times 10^{1}$ | 4.01 | 8.97 | 2.63 | 265.40 |
| $\mu_2$ | $1 \times 10^{-1}$ | $1.0 \times 10^{1}$ | 4.00 | 6.11 | $2.44 \times 10^{1}$ | 4.35 | 7.72 | 3.12 | 6.74 | 2.32 | 87.97 |
| $\mu_3$ | $1 \times 10^{-1}$ | $1.0 \times 10^{1}$ | 2.00 | $1.00 \times 10^{1}$ | $4.63 \times 10^{1}$ | 9.98 | $3.61 \times 10^{1}$ | $1.00 \times 10^{1}$ | 38.70 | 0.10 | 423.70 |
| $\beta_{11}$ | $-5 \times 10^{-1}$ | $5 \times 10^{-1}$ | $-5 \times 10^{-2}$ | $-3.07 \times 10^{-2}$ | $6.41 \times 10^{-1}$ | $1.7 \times 10^{-3}$ | $6.02 \times 10^{-1}$ | $3.12 \times 10^{-2}$ | 0.27 | -0.50 | 183.37 |
| $\beta_{12}$ | $-1.5$ | 1.5 | $1.5 \times 10^{-1}$ | $6.98 \times 10^{-2}$ | $5.51 \times 10^{-1}$ | $2.17 \times 10^{-1}$ | $8.54 \times 10^{-1}$ | $-4.36 \times 10^{-2}$ | 0.30 | 0.47 | 134.19 |
| $\beta_{13}$ | $-2.0$ | 2.0 | $-2 \times 10^{-1}$ | $-7.85 \times 10^{-2}$ | 1.35 | $-4.36 \times 10^{-1}$ | 1.03 | $5.09 \times 10^{-3}$ | 0.07 | 0.10 | 112.66 |
| $\beta_{21}$ | $-1 \times 10^{-1}$ | $1.0 \times 10^{-1}$ | $-1 \times 10^{-2}$ | $-6.30 \times 10^{-2}$ | $7.96 \times 10^{-1}$ | $5.87 \times 10^{-2}$ | $4.10 \times 10^{-1}$ | $2.08 \times 10^{-2}$ | 0.22 | -0.033 | 51.40 |
| $\beta_{22}$ | $-3 \times 10^{-1}$ | $3 \times 10^{-1}$ | $-2.6 \times 10^{-2}$ | $-9.87 \times 10^{-2}$ | $9.0 \times 10^{-1}$ | $-1.08 \times 10^{-1}$ | $6.78 \times 10^{-1}$ | $-3.23 \times 10^{-2}$ | 0.25 | 0.11 | 36.75 |
| $\beta_{23}$ | $-5 \times 10^{-1}$ | $5 \times 10^{-1}$ | $5 \times 10^{-2}$ | $2.89 \times 10^{-1}$ | 1.20 | $6.65 \times 10^{-2}$ | $6.94 \times 10^{-1}$ | $8.27 \times 10^{-3}$ | 0.04 | -0.14 | 33.26 |
| $\beta_{31}$ | $-1.0$ | 1.0 | $1 \times 10^{-1}$ | $2.12 \times 10^{-1}$ | 1.14 | $4.05 \times 10^{-1}$ | 1.01 | $1.60 \times 10^{-1}$ | 1.15 | -0.65 | 287.92 |
| $\beta_{32}$ | $-1.0$ | 1.0 | $-1 \times 10^{-1}$ | $-3.28 \times 10^{-1}$ | 1.54 | $-5.40 \times 10^{-1}$ | 1.70 | $-1.86 \times 10^{-1}$ | 1.28 | 0.77 | 210.00 |
| $\beta_{33}$ | $-1.5 \times 10^{-1}$ | $1.5 \times 10^{-1}$ | $-1.48 \times 10^{-2}$ | $1.50 \times 10^{-1}$ | 1.92 | $1.50 \times 10^{-1}$ | 2.02 | $-1.30 \times 10^{-2}$ | $2.42 \times 10^{-1}$ | -0.15 | 178.02 |
| $X_1$ | 0 | $2.0 \times 10^{1}$ | $1.00 \times 10^{1}$ | $1.59 \times 10^{1}$ | $3.85 \times 10^{1}$ | $2.00 \times 10^{1}$ | $5.03 \times 10^{1}$ | $2.64 \times 10^{1}$ | $4.35 \times 10^{1}$ | $2.00 \times 10^{1}$ | $7.77 \times 10^{1}$ |
| $X_2$ | 0 | $2.0 \times 10^{1}$ | $1.40 \times 10^{1}$ | $2.00 \times 10^{1}$ | $6.57 \times 10^{1}$ | $2.00 \times 10^{1}$ | $2.02 \times 10^{1}$ | $4.07 \times 10^{1}$ | $4.97 \times 10^{1}$ | $2.00 \times 10^{1}$ | $6.19 \times 10^{1}$ |
| $X_3$ | 0 | $1.0 \times 10^{1}$ | 4.0 | $1.00 \times 10^{1}$ | $3.86 \times 10^{1}$ | 4.25 | $1.78 \times 10^{1}$ | 2.90 | $1.42 \times 10^{1}$ | 8.58 | $4.21 \times 10^{1}$ |
| $f_{best}$ | | | $2.9691 \times 10^{1}$ | 20.564 | | 16.971 | | 13.027 | | 22.886 | |

Table V: Case study ***GLV3***: Results for the *Generalized Lotka-Volterra* 3 species model with 10% homoscedastic Gaussian noise for three overfitting scenarios, and good fit. The table represents the lower and upper bounds (**LB**, **UB**), nominal and estimated values, as well as the absolute errors (**EA**), for each estimated parameter. Values in blue indicate a sign change with respect to the nominal value.

We selected the overfitting case, the well-fitted model and local optimum (underfitting) from Figure M. To further evaluate the residuals, we performed a QQ analysis, which revealed a deviation from normality in the overfitted case, whereas the well-fitted model showed residuals closely following a normal distribution.

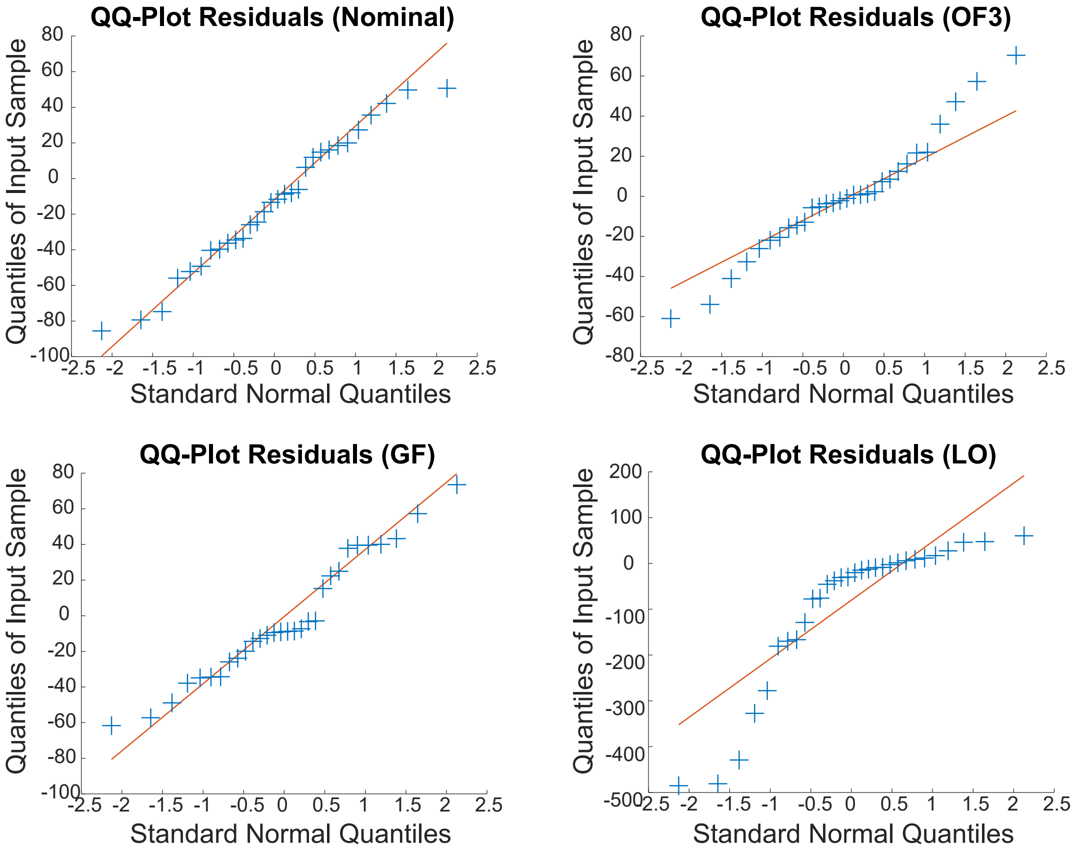

Figure O: Case study ***GLV3***: QQanalysis of the cases from Figure M, overfitting (OF3), underfitting (local optimum, LO), a good fit (GF) and the nominal fit are presented.

##### 6.6.5 Stability Analysis

We perform a stability analysis for the ***GLV3*** model using nominal, overfit and good fit parameter sets (Table W, Table X, Table Y). This analysis aims to identify the steady states of the system, evaluate their stability through eigenvalues of the Jacobian matrix, and visualize the system's dynamics using phase portraits and population trajectories. The phase portraits are shown in Figure P.

| Nominal fit | | | |
|---|---|---|---|
| **Steady State** | **($X_1$,$X_2$,$X_3$)** | **Eigenvalues of the Jacobian** | **Stability** |
| 1 | (0, 0, 0) | 2.00<br>4.00<br>6.00 | Unstable |
| 2 | (120, 0, 0) | $-6.00$<br>2.80<br>14.00 | Saddle Point |
| 3 | (270, 50, 0) | $-11.52$<br>$-3.28$<br>24.00 | Saddle Point |
| 4 | (0, 153.85, 0) | $-4.00$<br>29.08<br>$-13.38$ | Saddle Point |
| 5 | (0, 0, 135.14) | $-2.00$<br>$-21.03$<br>10.76 | Saddle Point |
| **6** | (481.14, 463.10, 257.04) | $-17.92 + 57.99i$<br>$-17.92 - 57.99i$<br>$-4.06 + 0.00i$ | Stable |

Table W: Case study ***GLV3***: Summary of steady states with nominal parameters, including their coordinates, eigenvalues of the Jacobian matrix, and stability classification.

| OF3 | | | |
|---|---|---|---|
| **Steady State** | **($X_1$,$X_2$,$X_3$)** | **Eigenvalues of the Jacobian** | **Stability** |
| 1 | $(0, 0, 0)$ | 3.12<br>4.01<br>10.00 | Unstable |
| 2 | $(0, 0, 769.23)$ | $-10.00$<br>7.93<br>9.50 | Saddle point |
| 3 | $(0, 96.57, 0)$ | $-3.12$<br>$-0.20$<br>$-7.98$ | Stable |
| 4 | $(64.61, 138.18, 0)$ | 0.32<br>$-2.77$<br>$-5.40$ | Saddle point |
| **5** | $(681.45, 616.50, 315.63)$ | $0.090 + 8.83i$<br>$0.090 - 8.83i$<br>$-2.93 + 0.00i$ | Saddle point |

Table X: Case study ***GLV3***: Summary of steady states with estimated parameters obtained in the overfitting case 3 (**OF3**), including their coordinates, eigenvalues of the Jacobian matrix, and stability classification.

| Good fit | | | |
|---|---|---|---|
| **Steady State** | **($X_1$,$X_2$,$X_3$)** | **Eigenvalues of the Jacobian** | **Stability** |
| 1 | $(0, 0, 0)$ | $-0.10$<br>2.19<br>2.83 | Saddle point |
| 2 | $(0, 0, 0.67)$ | $-0.10$<br>2.19<br>2.83 | Saddle point |
| 3 | $(5.27, 0, 0)$ | $-0.10$<br>2.19<br>2.83 | Saddle point |
| 4 | $(2.33, 0, 2.24)$ | $-0.10$<br>2.19<br>2.83 | Saddle point |

Table Y: Case study ***GLV3***: Summary of steady states with estimated parameters obtained in the good fit case (**Good Fit**), including their coordinates, eigenvalues of the Jacobian matrix, and stability classification.

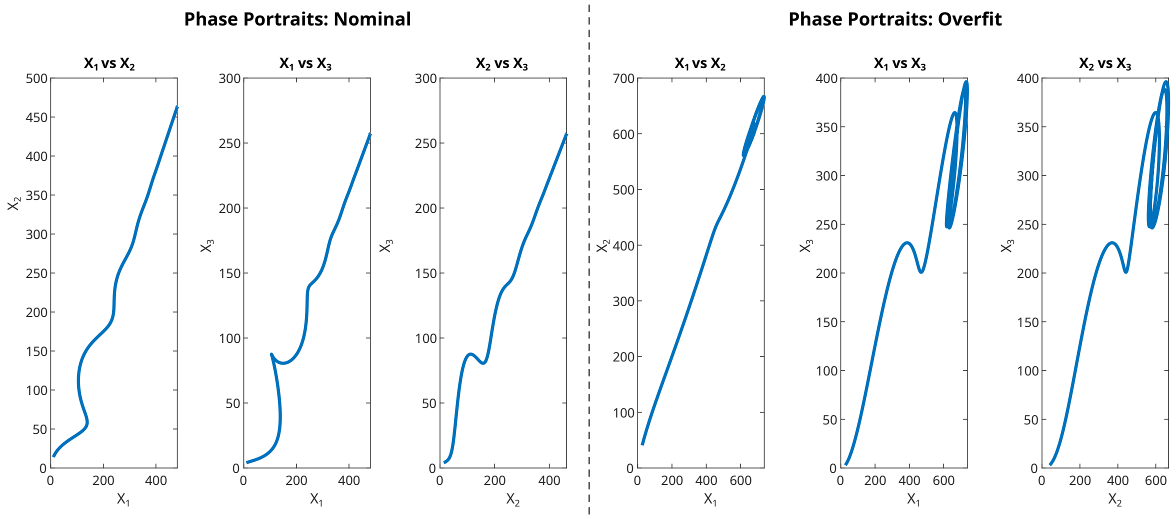

Figure P: Case study ***GLV3***: Phase portraits of the 3-species *Generalized Lotka-Volterra* model. The plots on the left correspond to the nominal parameters and initial conditions, while the plots on the right correspond to the overfitted parameters and initial conditions (**OF3**). Each column represents a different pairwise interaction: (**Left** ) Species 1 ($X_1$) vs. Species 2 ($X_2$), (**Center**) Species 1 ($X_1$) vs. Species 3 ($X_3$), and (**Right**) Species 2 ($X_2$) vs. Species 3 ($X_3$).

##### 6.6.6 Predictive power

Even though the estimation in Figure N (Overfitting 3) appears to be a good fit, its predictive capability is completely inaccurate. To demonstrate this, we changed the initial conditions from $X_{10} = 10.00$, $X_{20} = 14.00$, $X_{30} = 4.00$, to $X_{10} = 4.00$, $X_{20} = 10.00$, $X_{30} = 14.00$. When applying the parameters from the overfit scenario number 3 with these new initial conditions, we observe that the fit is completely erroneous, highlighting the issue of relying on an overfitted model (Figure Q).

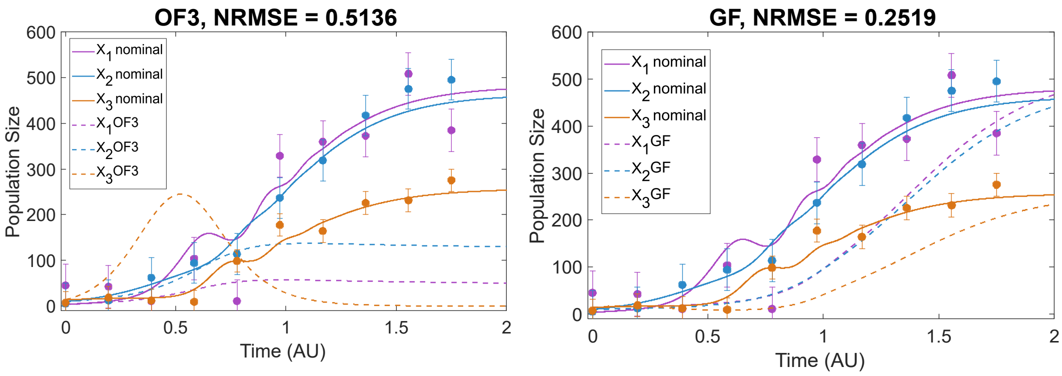

Figure Q: Case study ***GLV3***: Comparison of the fits obtained with new initial conditions ($X_{10} = 4.00$, $X_{20} = 10.00$, $X_{30} = 14.00$). The fit with dashed lines was generated using the parameters from the overfitting scenario number 3 (**OF3**) or the good fit (**GF**), while the fit with a solid line used the nominal parameters.

When working with experimental data in the laboratory, where the nominal parameters are unknown, there is a high risk of overfitting, especially when relying on local solvers. This is because local solvers may find solutions that fit the data well but do not accurately reflect the true underlying model, leading to poor predictive performance. Therefore, it is crucial to use global solvers, such as `Enhanced Scatter Search`, which are better suited for exploring the parameter space and avoiding local minima, ensuring more reliable and robust parameter estimates.

#### 6.7 ***CompV*: *Competition with a virus***

##### 6.7.1 Model

A mathematical model for a case where two bacteria ($X_1$, $X_2$) compete for a unique, essential growth-limiting substrate ($S$) in the presence of a virus ($V$) associated only with the first species is presented. Although the classical models without viral infection suggest competitive exclusion, this model (equations 21-24) exhibits the stable coexistence of both species (Alsolami and El Hajji, 2023).

$$\frac{dS}{dt} = -\mu_1(S, X_1)X_1 - \mu_2(S, X_2)X_2 + D(S_{in} - S) \tag{21}$$

$$\frac{dX_1}{dt} = \mu_1(S, X_1)X_1 - DX_1 - VX_1 \tag{22}$$

$$\frac{dX_2}{dt} = \mu_2(S, X_2)X_2 - DX_2 \tag{23}$$

$$\frac{dV}{dt} = \sigma VX_1 - DV \tag{24}$$

The specific growth functions are defined by:

$$\mu_i = \frac{\mu_{mi}S}{k_i + S} \tag{25}$$

Table Z summarizes the parameter values reported by Alsolami and El Hajji (2023). $D$ is the dilution rate, $\mu_i$ the maximum specific growth rate for species $i$, $k_i$ the half-saturation constant for species $i$, $S_{in}$ the input concentration of substrate $i$, $\sigma = \alpha \cdot k \cdot \gamma_1$, where $\alpha$ is the fixed number of contacts per unit of time that are sufficient to become infectious, $k$ denotes the species 1-to-virus yield, and $\gamma_1$ the yield coefficient referred to as the substrate-to-species 1.

| Parameter | Value [AU] | Parameter | Value [AU] | Parameter | Value [AU] |
|---|---|---|---|---|---|
| $\mu_{m1}$ | 4.00 | $S_{in}$ | 10.00 | $k_1$ | 6.00 |
| $\mu_{m2}$ | 3.00 | $D$ | 0.9375 | $k_2$ | 6.00 |
| $\sigma$ | 2.00 | | | | |
| **Initial conditions (IC):** $S_0 = 0.50$, $X_{10} = 0.50$, $X_{20} = 2.00$, $V_0 = 0.25$ | | | | | |

Table Z: Case study ***CompV***: Parameter values and initial conditions (IC) for the case study of the competitive model with presence of a virus. The parameters ($\mu_{m,i}$, $\sigma$, $k_i$, $D$, $S_{in}$) represent the maximum specific growth rates for species $i$, a fixed number, the half-saturation constant for species $i$ , dilution rate, and inlet substrate concentration, respectively. The initial conditions ($S_0$, $X_{10}$, $X_{20}$, $V_0$) denote the starting values for substrate, population densities, and virus. Nominal parameter values are obtained from Alsolami and El Hajji (2023).

##### 6.7.2 Structural Identifiability Analysis (SIA)

For the Competition with a virus model (***CompV***), the structural identifiability analysis conducted with `GenSSI2` is shown in Table . Both `SIAN` and `Structural Identifiability` provided consistent results (Table , Table ), identifying all parameters as globally identifiable, while `GenSSI2` classified some of them as locally identifiable. This discrepancy highlights the limitations of `GenSSI2` in ensuring global identifiability due to its symbolic manipulation approach, which can lead to conservative outcomes in complex models. Additionally, `GenSSI2` cannot handle arithmetic operations in the initial conditions, while `SIAN` and `Structural Identifiability` match the initial conditions with the observed states.

| | `GenSSI2` | (**IC Unknown**) | | | `GenSSI2` | (**IC Known**) | | |
|---|---|---|---|---|---|---|---|---|
| Obs. | Global | Local | Non-Id | Time[min] | Global | Local | Non-Id | Time[min] |
| $X_1$,$X_2$,$S$,$V$ | $X_{10}, X_{20}, S_0, V_0$ | $D, S_{in}, k_i, \mu_{mi}, \sigma$ | - | 60.86 | - | $D, S_{in}, k_i, \mu_{mi}, \sigma$ | - | 746.59 |
| $X_1$,$X_2$,$V$ | $X_{10}, X_{20}, V_0$ | $S_0, D, S_{in}, k_i, \mu_{mi}, \sigma$ | - | 16.14 | - | $D, S_{in}, k_i, \mu_{mi}, \sigma$ | - | 1521.13 |
| $X_1 + X_2$,$V$ | $V_0$ | $X_{10}, X_{20}, S_0, D, S_{in}, k_i, \mu_{mi}, \sigma$ | - | 28.76 | - | $D, S_{in}, k_i, \mu_{mi}, \sigma$ | - | 19.69 |

Table : Case study ***CompV***: Structural identifiability analysis (SIA) results obtained with `GenSSI2` for the *Competition with a virus* model. The analysis was performed considering three observed scenarios ($X_1$,$X_2$,$S$,$V$; $X_1$,$X_2$,$V$; $X_1 + X_2$,$V$), and two cases for initial conditions: unknown (IC Unknown) and known (IC Known). The parameters are classified as globally identifiable ("Global"), locally identifiable ("Local"), or non-identifiable ("Non-Id"). A " - " indicates that no parameters are found in that category. The table summarizes the computational time and the identifiability results for the model parameters ($D$, $S_{in}$, $k_i$, $\mu_{mi}$, $\sigma$). The results are the same for both known or unknown initial conditions.

| **IC Unknown** | `SIAN` | | | | `Structural Identifiability` | | | |
|---|---|---|---|---|---|---|---|---|
| Obs. | Global | Local | Non-Id | Time [s] | Global | Local | Non-Id | Time [s] |
| $X_1$,$X_2$,$S$,$V$ | $X_i, S, V, D$ $S_{in}, k_i, \mu_{mi}, \sigma$ | - | - | 57.79 | $X_i(t)$,$S(t)$,$V(t)$, $D$,$S_{in}$,$k_i$, $\mu_{mi}, \sigma$ | - | - | 12.48 |
| $X_1$,$X_2$,$V$ | $X_i, S, V, D$ $S_{in}, k_i, \mu_{mi}, \sigma$ | - | - | 58.50 | $X_i(t)$,$S(t)$,$V(t)$, $D$,$S_{in}$,$k_i$, $\mu_{mi}, \sigma$ | - | - | 14.28 |
| $X_1 + X_2$,$V$ | $X_i, S, V, D$ $S_{in}, k_i, \mu_{mi}, \sigma$ | - | - | 155.50 | $X_i(t)$,$S(t)$,$V(t)$, $D$,$S_{in}$,$k_i$, $\mu_{mi}, \sigma$ | - | - | 13.56 |

Table : Case study ***CompV***: Structural identifiability analysis (SIA) results obtained with `SIAN` and `Structural Identifiability` for the *Competition with a virus* model. The analysis was performed considering three observed scenarios ($X_1$,$X_2$,$S$,$V$; $X_1$,$X_2$,$V$; $X_1 + X_2$,$V$), and unknown initial conditions (IC Unknown). The parameters are classified as globally identifiable ("Global"), locally identifiable ("Local"), or non-identifiable ("Non-Id"). A " - " indicates that no parameters are found in that category. The table summarizes the computational time and the identifiability results for the model parameters ($D$, $S_{in}$, $k_i$, $\mu_{mi}$, $\sigma$).

| IC Known | SIAN | | | | Structural Identifiability | | | |
|---|---|---|---|---|---|---|---|---|
| Obs. | Global | Local | Non-Id | Time [s] | Global | Local | Non-Id | Time [s] |
| $X_1$,$X_2$,$S$,$V$ | $D, S_{in}$, $k_i, \mu_{mi}, \sigma$ | - | - | 57.76 | $X_{i0}$,$S_0$,$V_0$, $D$, $S_{in}$,$k_i$, $\mu_{mi}, \sigma$ | - | - | 24.20 |
| $X_1$,$X_2$,$V$ | $D, S_{in}$, $k_i, \mu_{mi}, \sigma$ | - | - | 56.23 | $X_{i0}$,$S_0$,$V_0$, $D$, $S_{in}$,$k_i$, $\mu_{mi}, \sigma$ | - | - | 16.45 |
| $X_1 + X_2$,$V$ | $D, S_{in}$, $k_i, \mu_{mi}, \sigma$ | - | - | 151.95 | $X_{i0}$,$S_0$,$V_0$, $D$, $S_{in}$,$k_i$, $\mu_{mi}, \sigma$ | - | - | 16.54 |

Table : Case study ***CompV***: Structural identifiability analysis (SIA) results obtained with `SIAN` and `Structural Identifiability`, for the *Competition with a virus* model. The analysis was performed considering three observed scenarios ($X_1$,$X_2$,$S$,$V$; $X_1$,$X_2$,$V$; $X_1 + X_2$,$V$), and known initial conditions (IC Known). The parameters are classified as globally identifiable ("Global"), locally identifiable ("Local"), or non-identifiable ("Non-Id"). A " - " indicates that no parameters are found in that category. The table summarizes the computational time and the identifiability results for the model parameters ($D$, $S_{in}$, $k_i$, $\mu_{mi}$, $\sigma$).

##### 6.7.3 Practical Identifiability Analysis (PIA)

We addressed a parameter estimation problem involving two observed populations ($X_1$, $X_2$), the substrate ($S$), and the virus ($V$) with unknown initial conditions ($X_{10}$, $X_{20}$, $S_0$, $V_0$) (Figure R). To assess the robustness of our estimation methods, we generated three sets of pseudo-experimental data, each with a different Gaussian noise level: 0%, 5%, 10%. The generated data was homoscedastic and positive, reflecting the nature of population measurements.

For the dataset with 0% noise, we employed a weighted least squares approach as the cost function. For datasets with 5% and 10% noise levels, we used a cost function based on the negative log-likelihood, which is more appropriate for handling data with higher noise levels.

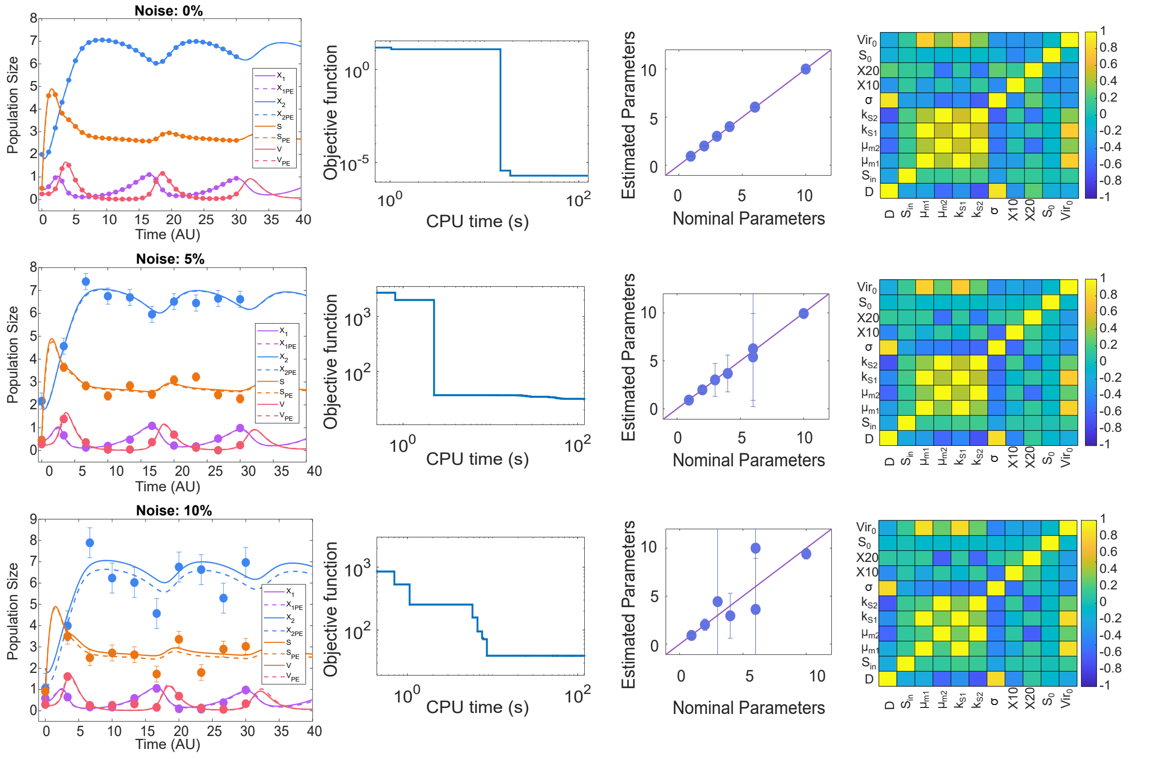

Figure R: Case study ***CompV***: Model calibration results for the *Competition with a virus* model (Alsolami and El Hajji, 2023). The first row displays results for the simulations with pseudo-experimental data and 0% noise, the second row shows results for 5% noise, and the third row corresponds to 10% noise. In each case, the simulation using the estimated parameters is compared to the simulation with nominal parameters, along with the convergence curve, correlation matrix, and a comparison of nominal and estimated parameters. These results demonstrate the model's ability to accurately fit the data under varying noise.

| | LB | UB | Nominal | Estimated | EA | ER (%) |
|---|---|---|---|---|---|---|
| $D$ | $1.00 \times 10^{-2}$ | 1.00 | $9.38 \times 10^{-1}$ | $9.37 \times 10^{-1}$ | $1.84 \times 10^{-4}$ | $6.04 \times 10^{-2}$ |
| $S_{in}$ | 1.00 | $1.00 \times 10^{2}$ | $1.00 \times 10^{1}$ | $1.00 \times 10^{1}$ | $4.04 \times 10^{-4}$ | $4.50 \times 10^{-3}$ |
| $\mu_{m1}$ | $1.00 \times 10^{-1}$ | $1.00 \times 10^{1}$ | 4.00 | 4.02 | $3.43 \times 10^{-3}$ | $3.91 \times 10^{-1}$ |
| $\mu_{m2}$ | $1.00 \times 10^{-1}$ | $1.00 \times 10^{1}$ | 3.00 | 3.01 | $2.58 \times 10^{-3}$ | $4.09 \times 10^{-1}$ |
| $k_1$ | $1.00 \times 10^{-1}$ | $1.00 \times 10^{1}$ | 6.00 | 6.04 | $7.96 \times 10^{-3}$ | $6.38 \times 10^{-1}$ |
| $k_2$ | $1.00 \times 10^{-1}$ | $1.00 \times 10^{1}$ | 6.00 | 6.04 | $8.78 \times 10^{-3}$ | $7.04 \times 10^{-1}$ |
| $\sigma$ | $1.00 \times 10^{-1}$ | $1.00 \times 10^{1}$ | 2.00 | 2.00 | $4.17 \times 10^{-4}$ | $6.52 \times 10^{-2}$ |
| $X_{10}$ | $5.00 \times 10^{-2}$ | 5.00 | $5.00 \times 10^{-1}$ | $5.00 \times 10^{-1}$ | $1.53 \times 10^{-4}$ | $1.46 \times 10^{-2}$ |
| $X_{20}$ | $2.00 \times 10^{-1}$ | $2.00 \times 10^{1}$ | 2.00 | 2.00 | $8.23 \times 10^{-4}$ | $5.62 \times 10^{-2}$ |
| $S_0$ | $5.00 \times 10^{-2}$ | 5.00 | $5.00 \times 10^{-1}$ | $4.99 \times 10^{-1}$ | $1.07 \times 10^{-3}$ | $1.08 \times 10^{-1}$ |
| $V_0$ | $2.50 \times 10^{-2}$ | 2.50 | $2.50 \times 10^{-1}$ | $2.51 \times 10^{-1}$ | $1.82 \times 10^{-4}$ | $2.66 \times 10^{-1}$ |

Table : Case study ***CompV***: Results for the *Competition with a virus* model with 0% homoscedastic Gaussian noise. The table displays the lower and upper bounds (**LB**, **UB**), nominal and estimated values, as well as the absolute and relative errors (**EA**, **ER**), for each estimated parameter.

| | LB | UB | Nominal | Estimated | EA | ER (%) |
|---|---|---|---|---|---|---|
| $D$ | $1.00 \times 10^{-2}$ | 1.00 | $9.38 \times 10^{-1}$ | $9.04 \times 10^{-1}$ | $1.28 \times 10^{-1}$ | 3.62 |
| $S_{in}$ | 1.00 | $1.00 \times 10^{2}$ | $1.00 \times 10^{1}$ | 9.94 | $2.68 \times 10^{-1}$ | $6.28 \times 10^{-1}$ |
| $\mu_{m1}$ | $1.00 \times 10^{-1}$ | $1.00 \times 10^{1}$ | 4.00 | 3.68 | 1.92 | 7.91 |
| $\mu_{m2}$ | $1.00 \times 10^{-1}$ | $1.00 \times 10^{1}$ | 3.00 | 3.01 | 1.72 | $3.82 \times 10^{-1}$ |
| $k_1$ | $1.00 \times 10^{-1}$ | $1.00 \times 10^{1}$ | 6.00 | 5.41 | 4.53 | 9.85 |
| $k_2$ | $1.00 \times 10^{-1}$ | $1.00 \times 10^{1}$ | 6.00 | 6.26 | 6.04 | 4.33 |
| $\sigma$ | $1.00 \times 10^{-1}$ | $1.00 \times 10^{1}$ | 2.00 | 1.97 | $2.98 \times 10^{-1}$ | 1.33 |
| $X_{10}$ | $5.00 \times 10^{-2}$ | 5.00 | $5.00 \times 10^{-1}$ | $4.94 \times 10^{-1}$ | $9.14 \times 10^{-2}$ | 1.14 |
| $X_{20}$ | $2.00 \times 10^{-1}$ | $2.00 \times 10^{1}$ | 2.00 | 2.06 | $5.56 \times 10^{-1}$ | 2.89 |
| $S_0$ | $5.00 \times 10^{-2}$ | 5.00 | $5.00 \times 10^{-1}$ | $4.60 \times 10^{-1}$ | $3.61 \times 10^{-1}$ | 8.07 |
| $V_0$ | $2.50 \times 10^{-2}$ | 2.50 | $2.50 \times 10^{-1}$ | $2.46 \times 10^{-1}$ | $1.25 \times 10^{-1}$ | 1.76 |

Table : Case study ***CompV***: Results for the *Competition with a virus* model with 5% homoscedastic Gaussian noise. The table displays the lower and upper bounds (**LB**, **UB**), nominal and estimated values, as well as the absolute and relative errors (**EA**, **ER**), for each estimated parameter.

| | LB | UB | Nominal | Estimated | EA | ER (%) |
|---|---|---|---|---|---|---|
| $D$ | $1.00 \times 10^{-2}$ | 1.00 | $9.38 \times 10^{-1}$ | $9.06 \times 10^{-1}$ | $2.56 \times 10^{-1}$ | 3.40 |
| $S_{in}$ | 1.00 | $1.00 \times 10^{2}$ | $1.00 \times 10^{1}$ | 9.41 | $5.36 \times 10^{-1}$ | 5.90 |
| $\mu_{m1}$ | $1.00 \times 10^{-1}$ | $1.00 \times 10^{1}$ | 4.00 | 2.95 | 2.35 | $2.61 \times 10^{1}$ |
| $\mu_{m2}$ | $1.00 \times 10^{-1}$ | $1.00 \times 10^{1}$ | 3.00 | 4.43 | 8.37 | $4.77 \times 10^{1}$ |
| $k_1$ | $1.00 \times 10^{-1}$ | $1.00 \times 10^{1}$ | 6.00 | 3.62 | 5.32 | $3.97 \times 10^{1}$ |
| $k_2$ | $1.00 \times 10^{-1}$ | $1.00 \times 10^{1}$ | 6.00 | $1.00 \times 10^{1}$ | $2.63 \times 10^{1}$ | $6.67 \times 10^{1}$ |
| $\sigma$ | $1.00 \times 10^{-1}$ | $1.00 \times 10^{1}$ | 2.00 | 2.02 | $6.00 \times 10^{-1}$ | 1.12 |
| $X_{10}$ | $5.00 \times 10^{-2}$ | 5.00 | $5.00 \times 10^{-1}$ | $5.03 \times 10^{-1}$ | $1.81 \times 10^{-1}$ | $5.84 \times 10^{-1}$ |
| $X_{20}$ | $2.00 \times 10^{-1}$ | $2.00 \times 10^{1}$ | 2.00 | 1.28 | 1.14 | $3.59 \times 10^{1}$ |
| $S_0$ | $5.00 \times 10^{-2}$ | 5.00 | $5.00 \times 10^{-1}$ | $8.49 \times 10^{-1}$ | $7.23 \times 10^{-1}$ | $6.99 \times 10^{1}$ |
| $V_0$ | $2.50 \times 10^{-2}$ | 2.50 | $2.50 \times 10^{-1}$ | $2.11 \times 10^{-1}$ | $2.45 \times 10^{-1}$ | $1.58 \times 10^{1}$ |

Table : Case study ***CompV***: Results for the *Competition with a virus* model with 10% homoscedastic Gaussian noise. The table displays the lower and upper bounds (**LB**, **UB**), nominal and estimated values, as well as the absolute and relative errors (**EA**, **ER**), for each estimated parameter.

After estimating the parameters, we conducted a sensitivity analysis to evaluate their influence on the population dynamics. This involved calculating the mean relative sensitivity (Figure S) and perturbing individually each parameter by +5% to identify which ones have the greatest impact on each variable (Figure T).

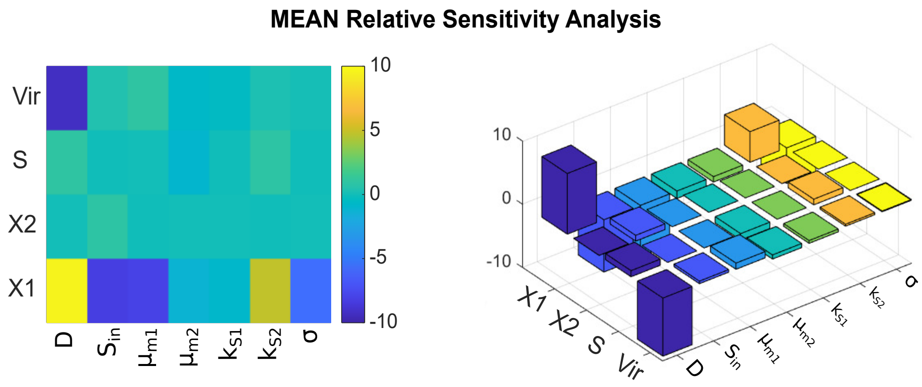

Figure S: Case study ***CompV***: Mean Sensitivity Analysis for the *Competition with a virus* model. **Left**: A heatmap illustrates the sensitivity of each observable (Y-axis) to each parameter (X-axis), highlighting how parameter variations affect the system's behavior. **Right**: The same analysis is presented in 3D, offering a clearer visualization of the sensitivities across different parameters and observables.

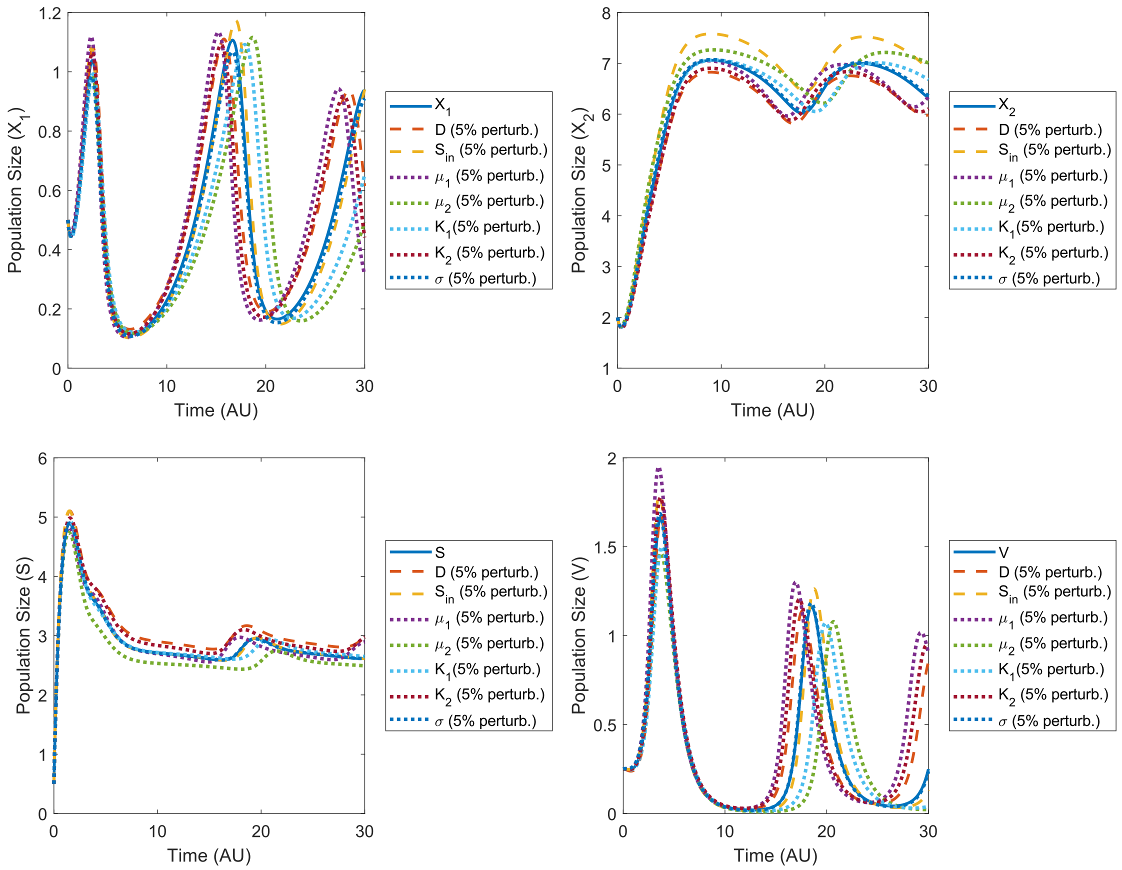

Figure T: Case study ***CompV***: Simulation results for the *Competition model with a virus*. The four plots show the dynamics of the observables: $X_1$ (top-left), $X_2$ (top-right), $S$ (bottom-left), and $V$ (bottom-right). In each plot, the simulations are presented with a positive 5% perturbation applied to each parameter separately, illustrating the effect of individual parameter variations on the system's behavior.

###### 6.7.4 Estimation using multistart of local methods

To highlight the advantages of using a global optimization strategy (eSS) over a multistart approach with local solvers, we present below the results obtained for this case study. The histograms illustrate the distribution of objective function values obtained from multiple runs, allowing for a comparison between the robustness and consistency of both approaches.

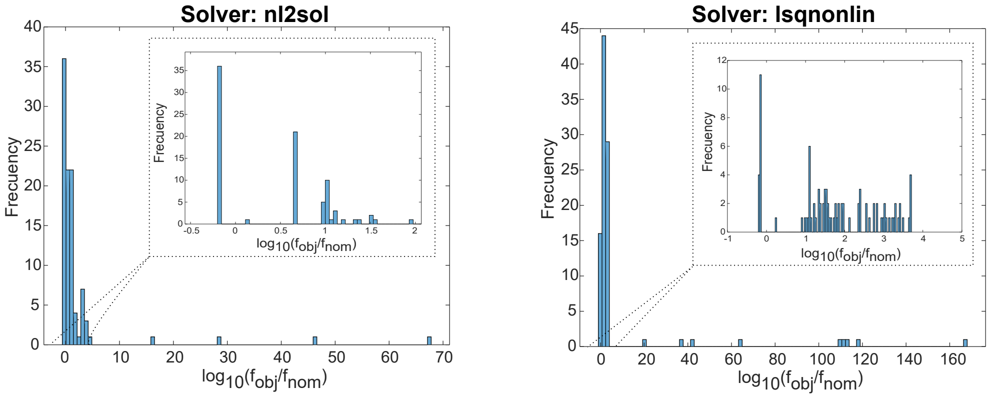

Figure U: Case study ***CompV***: Distribution of objective function values from 100 runs using the `nl2sol` (**Left**) and `lsqnonlin` (**Right**) local solvers. The `lsqnonlin` solver converged to 98 optima out of 100 runs, while `nl2sol` converged to 100 out of 100 runs. For `nl2sol`, 36 out of 100 runs resulted in a logarithm of the normalized objective function value ($log_{10}(f_{obj}/f_{nom})$) below 0, whereas 64 runs produced higher values. For `lsqnonlin`, 15 out of 100 runs achieved values below 0, with 83 runs yielding higher outcomes.

#### 6.8 *PCM*: *Phage Cocktail Model*

A large number of mathematical models have been developed to study the dynamics between host organisms and their parasites, and bacteria–phages are no exception. A *Phage Therapy* model proposed by Li *et al.* (2020) will be studied. It considers the nonlinear dynamics arising from interactions between sensitive bacteria $S$ (uninfected), phage-resistant bacteria $R$, phage $P_S$ (only targeting sensitive bacteria), phage $P_R$ (only targeting phage-resistant bacteria), and the host innate immune response $I$ (see Figure V).

Two bacterial strains ($S$ and $R$) reproduce within the constraints of the limited environmental capacity, $K_C$. Phage-resistant bacteria arise from sensitive bacteria through mutation, occurring with a fixed probability $\mu$ per cellular division. Both bacterial strains are targeted by the immune response, which eliminates them while also being activated by their presence. The immune system triggers an activation rate with a maximum value $\alpha$, reaching a capacity limit $K_I$. Phage populations, $P_S$ and $P_R$, infect and lyse sensitive and phage-resistant bacterial populations at rates $F(P_S)$ and $F(P_R)$, respectively. For simplicity , it is assumed that the two phage types ($P_S$ and $P_R$) have identical adsorption rate $\phi$, burst size $\beta$, and decay rate $\omega$ (Li *et al.*, 2020).

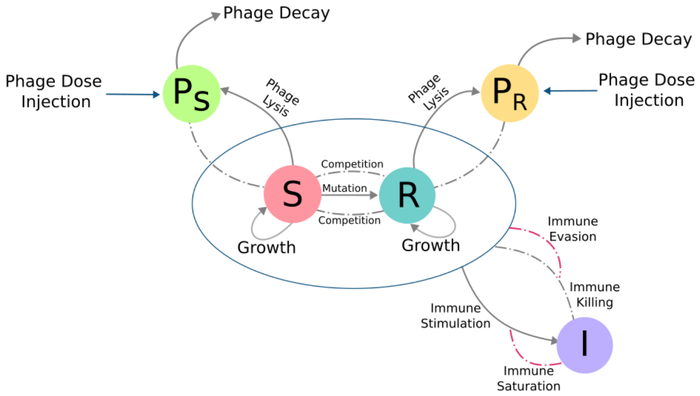

Figure V: Diagram of a schematic *Phage Therapy Model*. Sensitive bacteria ($S$) and phage-resistant bacteria ($R$) are targeted by phage ($P_S$) and phage ($P_R$). Innate immunity ($I$) is activated by the presence of bacteria and it attacks both bacterial strains. Adapted from (Li *et al.*, 2020).

##### 6.8.1 Model

The dynamics of bacteria, phage, and the innate immune system, can be modeled with the following system of nonlinear differential equations (eqs. 26- 31).

$$\frac{dS}{dt} = \underbrace{rS \cdot \left(1 - \frac{S+R}{K_C}\right) \cdot (1-\mu)}_{\text{logistic growth, mutation}} - \overbrace{S \cdot F(P_S)}^{\text{lysis}} - \underbrace{\frac{\epsilon IS}{1 + (S+R)/K_D}}_{\text{immune killing}} \tag{26}$$

$$\frac{dR}{dt} = \underbrace{r'R\left(1 - \frac{S+R}{K_C}\right)}_{\text{logistic growth}} + \overbrace{\mu rS \cdot \left(1 - \frac{S+R}{K_C}\right)}^{\text{mutation from sensitive host}} \tag{27}$$

$$-\underbrace{R \cdot F(P_R)}_{\text{lysis}} - \overbrace{\frac{\epsilon IR}{1 + (S+R)/K_D}}^{\text{immune killing}} \tag{28}$$

$$\frac{dP_s}{dt} = \underbrace{\beta \cdot S \cdot F(P_s)}_{\text{release of viruses}} - \overbrace{\phi \cdot S \cdot P_S}^{\text{adsorption}} - \underbrace{\omega \cdot P_S}_{\text{decay}} + \overbrace{\rho_s}^{\text{phage injection}} \tag{29}$$

$$\frac{dI}{dt} = \underbrace{\alpha \cdot I \cdot (1 - I/K_I) \cdot \left(\frac{S+R}{S+R+K_N}\right)}_{\text{immune stimulation, activation and immune saturation}} \tag{30}$$

$$\frac{dP_R}{dt} = \underbrace{\beta \cdot R \cdot F(P_R)}_{\text{release of viruses}} - \overbrace{\phi \cdot R \cdot P_R}^{\text{adsorption}} - \underbrace{\omega \cdot P_R}_{\text{decay}} + \overbrace{\rho_R}^{\text{phage injection}} \tag{31}$$

Where $F(P_i) = \phi P_i / \left(1 + \frac{P_i}{P_C}\right)$ for $i \in \{S, R\}$ is the per-capita phage-induced bacterial lysis rate that characterizes the effect of phage saturation during the infection. A description of the parameters can be found in Table .

| Parameter | Nominal Value | Description |
|---|---|---|
| $r$ | $7.5 \times 10^{-1}$ | Maximum growth rate of bacteria $[h^{-1}]$ |
| $r'$ | $6.75 \times 10^{-1}$ | Maximum growth rate of phage-resistant bacteria $[h^{-1}]$ |
| $\mu$ | $2.85 \times 10^{-8}$ | Probability of emergence of phage-resistant mutant per cellular division |
| $K_C$ | $1 \times 10^{10}$ | Carrying capacity of bacteria $[CFU/g]$[1] |
| $\beta$ | $1.00 \times 10^{2}$ | Burst size of phage |
| $\omega$ | $7 \times 10^{-2}$ | Decay rate of phage $[h^{-1}]$ |
| $\epsilon$ | $8.2 \times 10^{-8}$ | Killing rate parameter of the immune response $[g/(hcell)]$ |
| $\alpha$ | $9.7 \times 10^{-1}$ | Maximum growth rate of the immune response $[h^{-1}]$ |
| $K_I$ | $2.4 \times 10^{7}$ | Maximum capacity of the immune response $[cell/g]$ |
| $K_D$ | $4.1 \times 10^{7}$ | Bacterial concentration at which the immune response is half as effective $[CFU/g]$ |
| $K_N$ | $1.0 \times 10^{7}$ | Bacterial concentration when the immune response growth rate is half its maximum $[CFU/g]$ |
| $\phi$ | $5.4 \times 10^{-8}$ | Bacterial concentration when the immune response growth rate is half its maximum $[g/(hPFU)]$ |
| $P_C$ | $1.5 \times 10^{7}$ | Phage density at half saturation $[PFU/g]$[2] |
| $\rho$ | $0 \times 10^{7}$ | Injection rates $[PFU/(hg)]$ |

Table : Case study ***PCM***: Description of the parameters employed in the model. The nominal values were obtained from Li *et al.* (2020); Roach *et al.* (2017). For an immunodeficient host, the carrying capacity is set to $K_C = 8.5 \times 10^{11}$ $[CFU/g]$, with no innate immune activation ($K_I = I_0$).

##### 6.8.2 Structural Identifiability Analysis (SIA)

In the case of the *Phage Cocktail Model* (***PCM***), structural identifiability analysis using `GenSSI2` reveals significant limitations. For the fully observed scenario, `GenSSI2` classifies parameters as locally identifiable due to its symbolic manipulation constraints. Moreover, execution times are notably longer, and the analysis fails with an "*Out of Memory*" when considering the observables ($S$, $R$, $P_S$, $P_R$). `SIAN` and `Structural Identifiability` tools demonstrate superior performance, they are able to resolve the identifiability problem in most cases within less than two minutes.

| **IC Unknown** | `GenSSI2` | | | |
|---|---|---|---|---|
| Obs. | Global | Local | Non-Identifiable | Time [min] |
| $S, R, I$ $P_S, P_R$ | $S_0, R_0, P_{S0}, I_0, P_{R0}$ | $r$,$r'$,$\mu$,$P_C$,$K_C$, $K_D$,$K_I$,$K_N$,$\alpha$, $\beta$,$\phi$,$\omega$,$\epsilon$,$\rho_S$,$\rho_R$ | - | 54.79 |
| $S, R,$ $P_S, P_R$ | - | - | - | ** |

Table : Case study ***PCM***: Structural identifiability analysis (SIA) results obtained with `GenSSI2` for the *Phage Cocktail Model.* The analysis was performed considering two observation scenarios ($S$,$R$,$I$,$P_S$,$P_R$; $S$,$R$,$P_S$,$P_R$) and unknown initial conditions (IC Unknown). The parameters are classified as globally identifiable ("Global"), locally identifiable ("Local"), or non-identifiable ("Non-Identifiable"). A " - " indicates that no parameters are found in that category. The table summarizes the computational time and the identifiability results for the model parameters ($r$, $r'$, $\mu$, $P_C$, $K_C$, $K_D$, $K_I$, $K_N$, $\alpha$, $\beta$, $\phi$, $\omega$, $\epsilon$, $\rho_S$, $\rho_R$). The partially observed problem with known initial conditions (IC Known) failed to produce a result even after 30 hours of computation.

[1] $CFU/g$ stands for "Colony-Forming Units per gram". For example, if a soil sample is appropriately diluted and plated on agar, and after incubation, 50 visible colonies are counted on a plate with a 1:10 dilution, the estimated concentration would be $500 CFU/g$. This indicates the presence of approximately 500 viable bacterial colonies per gram of soil.

[2] $PFU/g$ typically stands for "Plaque-Forming Units per gram". For instance, if a sample is cultivated and results in 100 plaques, and the initial sample weight was 0.1 grams, the concentration would be $1000 PFU/g$ (100 plaques / 0.1 grams).

| **IC Unknown** | SIAN | | | | Structural Identifiability | | | |
|---|---|---|---|---|---|---|---|---|
| Obs. | Global | Local | Non-Id | Time[s] | Global | Local | Non-Id | Time[s] |
| $S, R, I$ $P_S, P_R$ | $S, R, P_S, I, P_R$, $r$,$r'$,$\mu$,$P_C$,$K_C$, $K_D$,$K_I$, $K_N$,$\alpha$, $\beta$,$\phi$,$\omega$,$\epsilon$,$\rho_S$,$\rho_R$ | - | - | 111.41 | $S(t)$,$R(t)$,$P_S(t)$, $I(t)$,$P_R(t)$, $r$,$r'$,$\mu$, $P_C$,$K_C$,$K_D$,$K_I$, $K_N$,$\alpha$,$\beta$,$\phi$,$\omega$,$\epsilon$,$\rho_S$,$\rho_R$ | - | - | 23.40 |
| $S, R$, $P_S, P_R$ | $S, R, P_S, P_R$, $r$,$r'$,$\mu$,$P_C$,$K_C$, $K_D$, $K_N$,$\alpha$, $\beta$,$\phi$,$\omega$,$\rho_S$,$\rho_R$ | - | $I, K_I, \epsilon$ | 146.01 | $S(t)$,$R(t)$,$P_S(t)$, $P_R(t)$,$r$,$r'$,$\mu$,$P_C$, $K_C$,$K_D$, $K_N$, $\alpha$,$\beta$,$\phi$,$\omega$,$\rho_S$,$\rho_R$ | - | $I(t), K_I, \epsilon$ | 29.32 |

Table : Case study ***PCM***: Structural identifiability analysis (SIA) results obtained with SIAN and Structural Identifiability for the *Phage Cocktail Model.* The analysis was performed considering two observation scenarios ($S$,$R$,$I$,$P_S$,$P_R$; $S$,$R$,$P_S$,$P_R$), and unknown initial conditions (IC Unknown). The parameters are classified as globally identifiable ("Global"), locally identifiable ("Local"), or non-identifiable ("Non-Id"). A " - " indicates that no parameters are found in that category. The table summarizes the computational time and the identifiability results for the model parameters ($r$, $r'$, $\mu$, $P_C$, $K_C$, $K_D$, $K_I$, $K_N$, $\alpha$, $\beta$, $\phi$, $\omega$, $\epsilon$, $\rho_S$, $\rho_R$).

| **IC Known** | SIAN | | | | Structural Identifiability | | | |
|---|---|---|---|---|---|---|---|---|
| Obs. | Global | Local | Non-Id | Time[s] | Global | Local | Non-Id | Time[s] |
| $S, R, I$ $P_S, P_R$ | $r$,$r'$,$\mu$,$P_C$,$K_C$, $K_D$,$K_I$,$K_N$,$\alpha$, $\beta$,$\phi$,$\omega$,$\epsilon$,$\rho_S$,$\rho_R$ | - | - | 110.26 | $S_0$,$R_0$,$P_{S0}$ $I_0$, $P_{R0}$,$r$,$r'$,$\mu$,$P_C$, $K_C$,$K_D$,$K_I$,$K_N$, $\alpha$,$\beta$,$\phi$,$\omega$,$\epsilon$,$\rho_S$,$\rho_R$ | - | - | 38.17 |
| $S, R$, $P_S, P_R$ | $r$,$r'$,$\mu$,$P_C$,$K_C$, $K_D$,$K_N$,$\alpha$, $\beta$,$\phi$,$\omega$,$\rho_S$,$\rho_R$ | - | $K_I$,$\epsilon$ | 141.08 | $S_0$,$R_0$,$P_{S0}$, $P_{R0}$,$r$,$r'$,$\mu$,$P_C$, $K_C$,$K_D$,$K_N$,$\alpha$, $\beta$,$\phi$,$\omega$,$\rho_S$,$\rho_R$ | - | $I_0$,$K_I$,$\epsilon$ | 43.08 |

Table : Case study ***PCM***: Structural identifiability analysis (SIA) results obtained with SIAN and Structural Identifiability for the *Phage Cocktail Model.* The analysis was performed considering two observation scenarios ($S$,$R$,$I$,$P_S$,$P_R$; $S$,$R$,$P_S$,$P_R$), and known initial conditions (IC Known). The parameters are classified as globally identifiable ("Global"), locally identifiable ("Local"), or non-identifiable ("Non-Id"). A " - " indicates that no parameters are found in that category. The table summarizes the computational time and the identifiability results for the model parameters ($r$, $r'$, $\mu$, $P_C$, $K_C$, $K_D$, $K_I$, $K_N$, $\alpha$, $\beta$, $\phi$, $\omega$, $\epsilon$, $\rho_S$, $\rho_R$).

##### 6.8.3 Practical Identifiability Analysis (PIA)

We addressed a parameter estimation problem involving all the observed states, $S$, $R$, $I$, $P_R$, $P_S$, with unknown initial conditions ($S_0$, $R_0$, $I_0$, $P_{R0}$, $P_{S0}$). To evaluate the performance and robustness of our estimation methods, we generated three sets of pseudo-experimental data, each incorporating different levels of Gaussian noise: 0%, 5%, and 10%. The generated data was heteroscedastic and positive, as a result of the differing magnitudes among the observables in the population measurements.

For this model, under the noiseless data scenario, we employed the *least-squares* criterion configured as *Q_expmax* with a tolerance of $10^{-3}$, which facilitates a more rapid decrease in the objective function. The estimated parameters obtained from this step served as the initial guess for a subsequent estimation, aiming to further refine the objective function. The new integration tolerance is set to $10^{-6}$. The results are shown in Figure W.

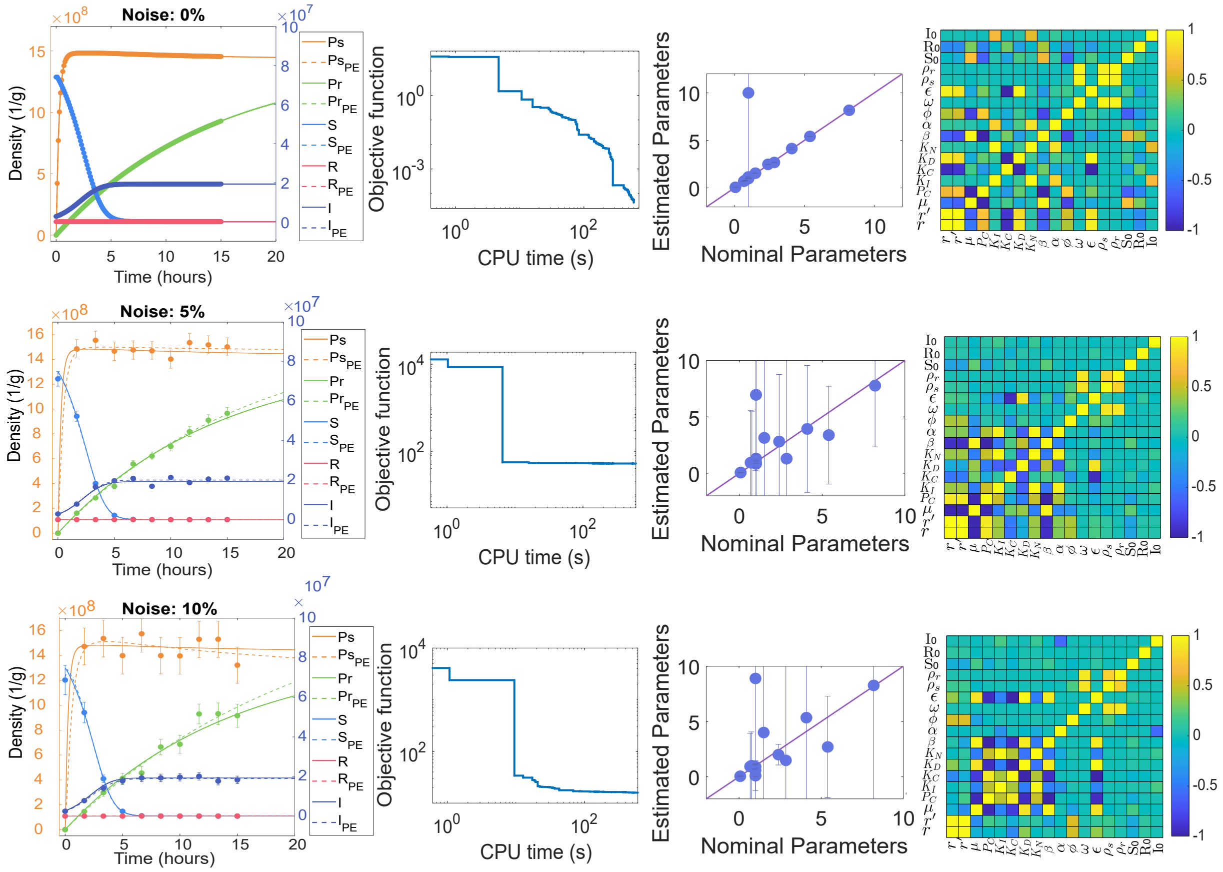

Figure W: Case study ***PCM***: Model calibration results for the *Phage Cocktail Model* (Li *et al.*, 2020). The first row displays results for the simulations with pseudo-experimental data and 0% noise, the second row shows results for 5% noise, and the third row corresponds to 10% noise. In each case, the simulation using the estimated parameters is compared to the simulation with nominal parameters, along with the convergence curve, correlation matrix, and a comparison of nominal and estimated parameters.

| | **LB** | **UB** | **Nominal** | **Estimated** | **EA** | **ER(%)** |
|---|---|---|---|---|---|---|
| $r$ | $1.00 \times 10^{-2}$ | 1.00 | $7.50 \times 10^{-1}$ | $5.69 \times 10^{-1}$ | 1.16 | $2.416 \times 10^{1}$ |
| $r'$ | $1.00 \times 10^{-2}$ | 1.00 | $6.75 \times 10^{-1}$ | $8.07 \times 10^{-1}$ | $1.50 \times 10^{5}$ | $1.958 \times 10^{1}$ |
| $\mu$ | $1.00 \times 10^{-1}$ | $1.00 \times 10^{1}$ | 2.85 | $1.62 \times 10^{-1}$ | $6.17 \times 10^{4}$ | $9.432 \times 10^{1}$ |
| $P_C$ | $1.00 \times 10^{-1}$ | $1.00 \times 10^{1}$ | 1.50 | $8.72 \times 10^{-1}$ | $5.98 \times 10^{-2}$ | $4.189 \times 10^{1}$ |
| $K_I$ | $1.00 \times 10^{-1}$ | $1.00 \times 10^{1}$ | 2.40 | 2.03 | $1.99 \times 10^{-1}$ | $1.550 \times 10^{1}$ |
| $K_C$ | $1.00 \times 10^{-1}$ | $1.00 \times 10^{1}$ | 1.00 | 8.81 | $1.92 \times 10^{4}$ | $7.8116 \times 10^{2}$ |
| $K_D$ | $1.00 \times 10^{-1}$ | $1.00 \times 10^{1}$ | 4.10 | 9.99 | 5.18 | $1.4376 \times 10^{2}$ |
| $K_N$ | $1.00 \times 10^{-1}$ | $1.00 \times 10^{1}$ | 1.00 | $1.00 \times 10^{-1}$ | $2.81 \times 10^{-1}$ | $9.000 \times 10^{1}$ |
| $\beta$ | $1.00 \times 10^{1}$ | $1.00 \times 10^{3}$ | 100 | $1.71 \times 10^{2}$ | $1.17 \times 10^{1}$ | $7.138 \times 10^{1}$ |
| $\alpha$ | $1.00 \times 10^{-2}$ | 1.00 | $9.70 \times 10^{-1}$ | $8.42 \times 10^{-1}$ | $7.44 \times 10^{-2}$ | $1.322 \times 10^{1}$ |
| $\phi$ | $1.00 \times 10^{-1}$ | $1.00 \times 10^{1}$ | 5.40 | 5.38 | $3.28 \times 10^{-2}$ | $3.1 \times 10^{-1}$ |
| $\omega$ | $1.00 \times 10^{-3}$ | $1.00 \times 10^{-1}$ | $7.00 \times 10^{-2}$ | $7.00 \times 10^{-2}$ | $7.47 \times 10^{-4}$ | $5 \times 10^{-2}$ |
| $\epsilon$ | $1.00 \times 10^{-1}$ | $1.00 \times 10^{1}$ | 8.20 | 8.26 | 6.74 | $7.6 \times 10^{-1}$ |
| $\rho_s$ | $1.00 \times 10^{-1}$ | $1.00 \times 10^{1}$ | 1.00 | $9.99 \times 10^{-1}$ | $1.10 \times 10^{-2}$ | $8 \times 10^{-2}$ |
| $\rho_r$ | $1.00 \times 10^{-1}$ | $1.00 \times 10^{1}$ | 1.00 | 1.00 | $5.45 \times 10^{-3}$ | $3 \times 10^{-2}$ |
| $S$ | $1.00 \times 10^{6}$ | $1.00 \times 10^{8}$ | $7.40 \times 10^{7}$ | $7.13 \times 10^{7}$ | $5.92 \times 10^{5}$ | 3.64 |
| $R$ | 0.00 | $1.00 \times 10^{1}$ | 1.00 | $4.83 \times 10^{-3}$ | $1.97 \times 10^{4}$ | $9.952 \times 10^{1}$ |
| $I$ | $1.00 \times 10^{5}$ | $1.00 \times 10^{7}$ | $2.70 \times 10^{6}$ | $3.14 \times 10^{6}$ | $3.42 \times 10^{5}$ | $1.647 \times 10^{1}$ |

Table : Case study ***PCM***: Results for the *Phage Cocktail Model* with 0% heteroscedastic Gaussian noise. The table presents the lower and upper bounds (**LB**, **UB**), nominal and estimated values, as well as the absolute and relative errors (**EA**, **ER**), for each estimated parameter.

| | LB | UB | Nominal | Estimated | EA | ER(%) |
|---|---|---|---|---|---|---|
| $r$ | $1.00 \times 10^{-2}$ | 1.00 | $7.50 \times 10^{-1}$ | $9.58 \times 10^{-1}$ | 4.50 | $2.77 \times 10^{1}$ |
| $r'$ | $1.00 \times 10^{-2}$ | 1.00 | $6.75 \times 10^{-1}$ | $9.37 \times 10^{-1}$ | 4.64 | $3.88 \times 10^{1}$ |
| $\mu$ | $1.00 \times 10^{-1}$ | $1.00 \times 10^{1}$ | $2.85 \times 10^{-8}$ | 1.29 | $1.22 \times 10^{1}$ | $4.52 \times 10^{8}$ |
| $P_C$ | $1.00 \times 10^{-1}$ | $1.00 \times 10^{1}$ | $1.50 \times 10^{7}$ | 3.14 | $1.27 \times 10^{1}$ | $1.00 \times 10^{2}$ |
| $K_I$ | $1.00 \times 10^{-1}$ | $1.00 \times 10^{1}$ | $2.40 \times 10^{7}$ | 2.81 | 5.95 | $1.00 \times 10^{2}$ |
| $K_C$ | $1.00 \times 10^{-1}$ | $1.00 \times 10^{1}$ | $1.00 \times 10^{10}$ | 6.95 | $6.84 \times 10^{3}$ | $1.00 \times 10^{2}$ |
| $K_D$ | $1.00 \times 10^{-1}$ | $1.00 \times 10^{1}$ | $4.10 \times 10^{7}$ | 3.93 | 5.63 | $1.00 \times 10^{2}$ |
| $K_N$ | $1.00 \times 10^{-1}$ | $1.00 \times 10^{1}$ | $1.00 \times 10^{7}$ | 1.30 | 9.98 | $1.00 \times 10^{2}$ |
| $\beta$ | $1.00 \times 10^{1}$ | $1.00 \times 10^{3}$ | 100 | $4.90 \times 10^{1}$ | $1.95 \times 10^{2}$ | $5.10 \times 10^{1}$ |
| $\alpha$ | $1.00 \times 10^{-2}$ | 1.00 | $9.70 \times 10^{-1}$ | $9.34 \times 10^{-1}$ | $7.04 \times 10^{-1}$ | 3.71 |
| $\phi$ | $1.00 \times 10^{-1}$ | $1.00 \times 10^{1}$ | $5.40 \times 10^{-8}$ | 3.38 | 4.34 | $6.26 \times 10^{9}$ |
| $\omega$ | $1.00 \times 10^{-3}$ | $1.00 \times 10^{-1}$ | $7 \times 10^{-2}$ | $5.74 \times 10^{-2}$ | $1.81 \times 10^{-2}$ | $1.80 \times 10^{1}$ |
| $\epsilon$ | $1.00 \times 10^{-1}$ | $1.00 \times 10^{1}$ | $8.20 \times 10^{-8}$ | 7.77 | 5.44 | $9.48 \times 10^{9}$ |
| $\rho_s$ | $1.00 \times 10^{-1}$ | $1.00 \times 10^{1}$ | $1.00 \times 10^{8}$ | $8.42 \times 10^{-1}$ | $3.07 \times 10^{-1}$ | $1.00 \times 10^{2}$ |
| $\rho_r$ | $1.00 \times 10^{-1}$ | $1.00 \times 10^{1}$ | $1.00 \times 10^{8}$ | $9.53 \times 10^{-1}$ | $7.30 \times 10^{-2}$ | $1.00 \times 10^{2}$ |
| $S$ | $1.00 \times 10^{6}$ | $1.00 \times 10^{8}$ | $7.40 \times 10^{7}$ | $7.13 \times 10^{7}$ | $7.25 \times 10^{6}$ | 3.65 |
| $R$ | 0.00 | $1.00 \times 10^{1}$ | 1.00 | $9.85 \times 10^{-1}$ | $9.79 \times 10^{-2}$ | 1.50 |
| $I$ | $1.00 \times 10^{5}$ | $1.00 \times 10^{7}$ | $2.70 \times 10^{6}$ | $2.84 \times 10^{6}$ | $2.64 \times 10^{5}$ | 5.18 |

Table : Case study ***PCM***: Results for the *Phage Cocktail Model* with 5% heteroscedastic Gaussian noise. The table represents the lower and upper bounds (**LB**, **UB**), nominal and estimated values, as well as the absolute and relative errors (**EA**, **ER**), for each estimated parameter.

| | LB | UB | Nominal | Estimated | EA | ER(%) |
|---|---|---|---|---|---|---|
| $r$ | $1.00 \times 10^{-2}$ | 1.00 | $7.50 \times 10^{-1}$ | 1.00 | 3.06 | $3.33 \times 10^{1}$ |
| $r'$ | $1.00 \times 10^{-2}$ | 1.00 | $6.75 \times 10^{-1}$ | $9.26 \times 10^{-1}$ | 3.01 | $3.72 \times 10^{1}$ |
| $\mu$ | $1.00 \times 10^{-1}$ | $1.00 \times 10^{1}$ | $2.85 \times 10^{-8}$ | 1.48 | $1.77 \times 10^{1}$ | $5.19 \times 10^{9}$ |
| $P_C$ | $1.00 \times 10^{-1}$ | $1.00 \times 10^{1}$ | $1.50 \times 10^{7}$ | 4.01 | $2.43 \times 10^{1}$ | $1.00 \times 10^{2}$ |
| $K_I$ | $1.00 \times 10^{-1}$ | $1.00 \times 10^{1}$ | $2.40 \times 10^{7}$ | 2.00 | $9.32 \times 10^{-1}$ | $1.00 \times 10^{2}$ |
| $K_C$ | $1.00 \times 10^{-1}$ | $1.00 \times 10^{1}$ | $1.00 \times 10^{10}$ | 8.90 | $5.64 \times 10^{4}$ | $1.00 \times 10^{2}$ |
| $K_D$ | $1.00 \times 10^{-1}$ | $1.00 \times 10^{1}$ | $4.10 \times 10^{7}$ | 5.35 | $4.50 \times 10^{1}$ | $1.00 \times 10^{2}$ |
| $K_N$ | $1.00 \times 10^{-1}$ | $1.00 \times 10^{1}$ | $1.00 \times 10^{7}$ | $1.00 \times 10^{-1}$ | 1.35 | $1.00 \times 10^{2}$ |
| $\beta$ | $1.00 \times 10^{1}$ | $1.00 \times 10^{3}$ | 100 | $3.96 \times 10^{1}$ | $2.34 \times 10^{2}$ | $6.04 \times 10^{-1}$ |
| $\alpha$ | $1.00 \times 10^{-2}$ | 1.00 | $9.70 \times 10^{-1}$ | $8.93 \times 10^{-1}$ | $2.28 \times 10^{-1}$ | 7.94 |
| $\phi$ | $1.00 \times 10^{-1}$ | $1.00 \times 10^{1}$ | $5.40 \times 10^{-8}$ | 2.70 | 4.60 | $4.99 \times 10^{8}$ |
| $\omega$ | $1.00 \times 10^{-3}$ | $1.00 \times 10^{-1}$ | $7 \times 10^{-2}$ | $4.71 \times 10^{-2}$ | $3.51 \times 10^{-2}$ | $3.27 \times 10^{1}$ |
| $\epsilon$ | $1.00 \times 10^{-1}$ | $1.00 \times 10^{1}$ | $8.20 \times 10^{-8}$ | 8.27 | $3.61 \times 10^{1}$ | $1.01 \times 10^{10}$ |
| $\rho_s$ | $1.00 \times 10^{-1}$ | $1.00 \times 10^{1}$ | $1.00 \times 10^{8}$ | $5.93 \times 10^{-1}$ | $5.89 \times 10^{-1}$ | $1.00 \times 10^{2}$ |
| $\rho_r$ | $1.00 \times 10^{-1}$ | $1.00 \times 10^{1}$ | $1.00 \times 10^{8}$ | $9.17 \times 10^{-1}$ | $1.41 \times 10^{-1}$ | $1.00 \times 10^{2}$ |
| $S$ | $1.00 \times 10^{6}$ | $1.00 \times 10^{8}$ | $7.40 \times 10^{7}$ | $7.02 \times 10^{7}$ | $1.45 \times 10^{7}$ | 5.14 |
| $R$ | 0.00 | $1.00 \times 10^{1}$ | 1.00 | $9.20 \times 10^{-1}$ | $1.96 \times 10^{-1}$ | 8.00 |
| $I$ | $1.00 \times 10^{5}$ | $1.00 \times 10^{7}$ | $2.70 \times 10^{6}$ | $2.48 \times 10^{6}$ | $5.23 \times 10^{5}$ | 8.15 |

Table : Case study ***PCM***: Results for the *Phage Cocktail Model* with 10% heteroscedastic Gaussian noise. The table represents the lower and upper bounds (**LB**, **UB**), nominal and estimated values, as well as the absolute and relative errors (**EA**, **ER**), for each estimated parameter.

After estimating the parameters, we conducted a sensitivity analysis to evaluate their influence on the population dynamics (Figure X).

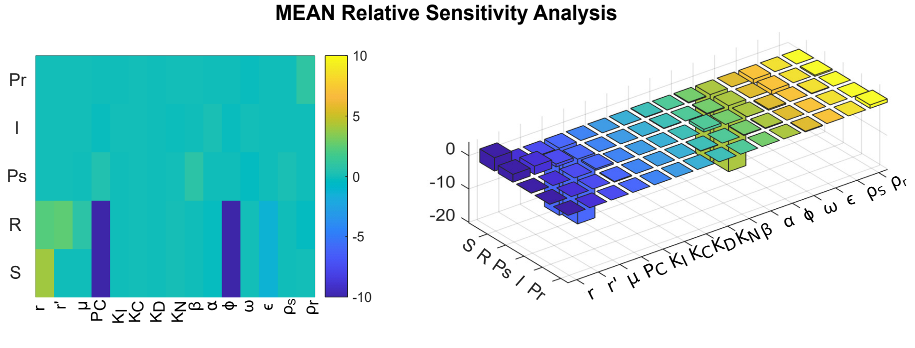

Figure X: Case study ***PCM***: Mean Sensitivity Analysis for the *Phage Cocktail Model*. **Left**: Heatmap with the sensitivity of each observable (Y-axis) to each parameter (X-axis), highlighting how parameter variations affect the system's behavior. **Right**: Sensitivity analysis presented in 3D.

##### 6.8.4 Estimation using multistart of local methods

To highlight the advantages of using a global optimization strategy (eSS) over a multistart approach with a local solver, we present below the results obtained for this case study. The histogram illustrates the distribution of objective function values obtained from multiple runs.

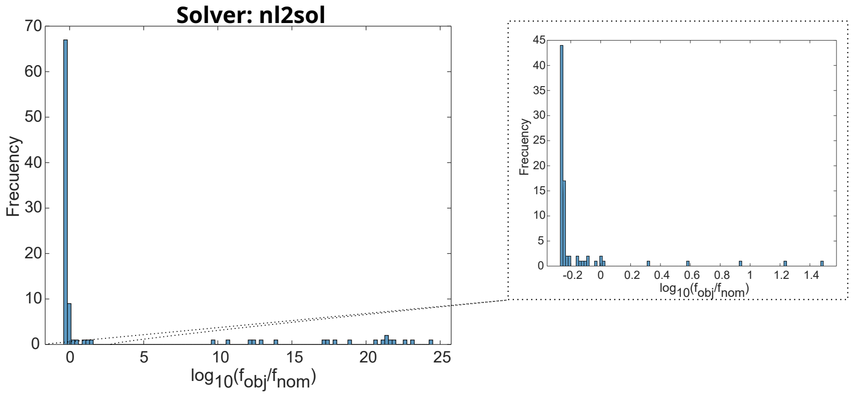

Figure Y: Case study ***PCM***: Distribution of objective function values from 100 runs using the `nl2sol` local solver. All runs successfully converged, 73 out of 100 runs resulted in a logarithm of the normalized objective function value ($log_{10}(f_{obj}/f_{nom})$) below 0, whereas 27 runs produced higher values.

#### 6.9 *EPHP*: *Enhanced Production of Heterologous Proteins*

A significant limitation of the previous models is that they neglect resource dynamics, making it difficult to understand how ecosystem properties depend on both the external environment and the consumer preferences of species. An alternative approach to the models already introduced is a *Consumer-Resource* explicit model ($CRMs$), where a population's growth rate depends on chemical mediators such as nutrients, toxins, or signals, whose concentrations are coupled to population dynamics. Unlike *Lotka–Volterra* models, where the species interactions are assumed a priori, forcing researchers to grapple with their mechanistic interpretation, *Consumer-Resource* models are "mechanistically" defined by how species consume and deplete resources, excrete by-products, and are influenced by signals, either positively or negatively.

In this type of model, species are modeled as consumers that can consume resources and sometimes also produce them. It has been proven that such models, initialized with random parameters, can predict lab experiments on complex microbial communities and reproduce large-scale ecological patterns, including the Earth and human microbiome projects (Goldford *et al.*, 2018; Marsland *et al.*, 2020).

The original *MacArthur Consumer-Resource* model (MacArthur and Levins, 1967) consists of $S$ consumers with abundances $N_i$ ($i = 1, ..., S$) that can consume one of $M$ possible resources with abundances $R_\alpha(\alpha = 1, ..., M)$, whose dynamics are described by the equations:

$$\frac{dN_i}{dt} = N_i \left( \sum_\beta C_{i\beta} R_\beta - m_i \right) \tag{32}$$

$$\frac{dR_\alpha}{dt} = R_\alpha \left( K_\alpha - R_\alpha - \sum_j N_j C_{j\alpha} \right) \tag{33}$$

The rate at which species consume a particular resource ($\alpha$) is represented by the value $C_{i\alpha}$ in the $S \times M$ matrix $\mathbf{C}$. $K_\alpha$ denotes the carrying capacity of the resource being consumed, while $m_i$ indicates the minimum amount of energy required by a species to survive.

##### 6.9.1 Model

This model, presented by Mauri *et al.* (2020), considers the *Enhanced Production of Heterologous Proteins* (***EPHP***) by a consortium (see Figure Z) consisting of two bacterial species (*E. coli*). The first bacterial species, which grows efficiently on glucose (the producer), produces a heterologous protein and secretes acetate as a by-product. The second species (the cleaner) grows preferentially on acetate, thereby removing it from the environment and alleviating its inhibitory effect on the growth of the producer. Assuming a well-stirred reaction volume, $B_P$ and $B_C$ denote the biomass concentrations of the producer and cleaner species, respectively. $H$ is the concentration of the heterologous protein, $A$ and $G$ denote the concentrations of acetate and glucose. This consortium grows in a bioreactor with equal and constant inflow and outflow rates, which ensures that the bioreactor volume remains constant, allowing for the eventual establishment of a well-defined steady state.

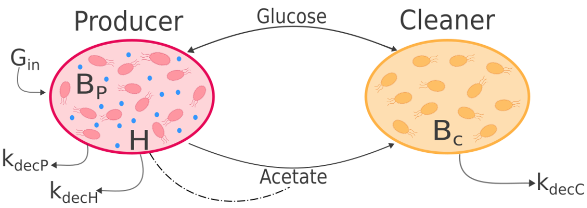

Figure Z: Synthetic consortium for the production of a heterologous protein.

The model introduced below describes the uptake of nutrients, glucose and acetate, at rates $r^g_{up}$, $r^a_{up}$, respectively. As well as the pathways for acetate overflow ($r^a_{over}$), production of autocatalytic biomass, and heterologous protein in proportions determined by the yield coefficients $Y_h$ and degradation of biomass ($k_{deg}$). The bacterial populations grow in a bioreactor, operating in continuous mode, whereby the dilution rate $D$ and the glucose concentration in the inflow $G_{in}$ can be tuned. Concentrations in the bioreactor are denoted by glucose ($G$), acetate ($A$), heterologous protein ($H$) and autocatalytic biomass ($B$) (Mauri *et al.*, 2020).

The ordinary differential equations defining the producer-cleaner consortium model are represented by the following set of equations (eqs. 34-38). A detailed explanation of each parameter is provided in Table ).

$$\frac{dG}{dt} = -r^g_{upP}B_P - r^g_{upC}B_C + D(G_{in} - G) \tag{34}$$

$$\frac{dA}{dt} = (r^a_{overP} - r^a_{upP})B_P + (r^a_{overC} - r^a_{upC})B_C - D \cdot A \tag{35}$$

$$\frac{dB_P}{dt} = (1 - Y_h)\left(Y_g r^g_{upP} + Y_a(r^a_{upP} - r^a_{overP})\right)B_P - k_{deg}B_P - D \cdot B_P \tag{36}$$

$$\frac{dH}{dt} = Y_h\left(Y_g r^g_{upP} + Y_a(r^a_{upP} - r^a_{overP})\right)B_P - k_{deg}H - D \cdot H \tag{37}$$

$$\frac{dB_C}{dt} = \left(Y_g r^g_{upC} + Y_a(r^a_{upC} - r^a_{overC})\right)B_C - k_{deg}B_C - D \cdot B_C \tag{38}$$

The glucose and acetate uptake, and acetate secretion rates of the producer, are expressed by:

$$r^g_{upP}(G, A) = k_g \frac{G}{G + K_g}\frac{\Theta^n_a}{A^n + \Theta^n_a} \tag{39}$$

$$r^a_{upP}(G, A) = k_a \frac{A}{A + K_a}\frac{\Theta^m_g}{r^g_{upP}(G, A)^m + \Theta^m_g} \tag{40}$$

$$r^a_{overP}(G, A) = k_{over} max(0, r^g_{upP}(G, A) - l) \tag{41}$$

The glucose and acetate uptake, and acetate secretion rates of the cleaner, are:

$$r^g_{upC}(G, A) = k_{\Delta PTS} \frac{G}{G + K_g}\frac{\Theta^n_a}{A^n + \Theta^n_a} \tag{42}$$

$$r^a_{upC}(G, A) = k_a \frac{A}{A + K_a}\frac{\Theta^m_g}{r^g_{upC}(G, A)^m + \Theta^m_g} + k_{Acs}\frac{A}{A + K_{Acs}} \tag{43}$$

$$r^a_{overC}(G, A) = k_{over} max(0, r^g_{upC}(G, A) - l) \tag{44}$$

| Parameter | Parameter Value | Description $[Units]$ |
|---|---|---|
| $D$ | $4.40 \times 10^{-1}$ | Dilution rate $[h^{-1}]$ |
| $G_{in}$ | $2.00 \times 10^{1}$ | Glucose inflow rate $[gh^{-1}]$ |
| $k_g$ | 1.53 | Glucose maximal uptake rate $[g\ gDW^{-1}h^{-1}]$ |
| $K_g$ | $9.00 \times 10^{-2}$ | Glucose half-maximal saturation $[gL^{-1}]$ |
| $\Theta_a$ | $5.20 \times 10^{-1}$ | Acetate inhibition constant $[gL^{-1}]$ |
| $n$ | 1.00 | Acetate inhibition strength |
| $k_{over}$ | $1.70 \times 10^{-1}$ | Acetate maximal overflow rate |
| $l$ | $7.00 \times 10^{-1}$ | Glucose uptake rate threshold for acetate overflow $[g\ gDW^{-1}h^{-1}]$ |
| $k_a$ | $9.70 \times 10^{-1}$ | Acetate maximal uptake rate $[g\ gDW^{-1}h^{-1}]$ |
| $K_a$ | $5.00 \times 10^{-1}$ | Acetate half-maximal saturation $[gL^{-1}]$ |
| $\Theta_g$ | $2.50 \times 10^{-1}$ | Carbon catabolite repression inhibition constant $[g\ gDW^{-1}h^{-1}]$ |
| $m$ | 1.00 | Carbon catabolite repression strength |
| $Y_g$ | $4.40 \times 10^{-1}$ | Biomass yield coefficient on glucose $[gDWg^{-1}]$ |
| $Y_a$ | $2.98 \times 10^{-1}$ | Biomass yield coefficient on acetate $[gDWg^{-1}]$ |
| $Y_h$ | $2.00 \times 10^{-1}$ | Heterologous protein yield constant |
| $k_{deg}$ | $4.4 \times 10^{-3}$ | Degradation rate $[h^{-1}]$ |
| $k_{\Delta PTS}$ | $3.80 \times 10^{-1}$ | Glucose maximal uptake rate for $\Delta PTS$ strain $[gDW^{-1}h^{-1}]$ |
| $k_{Acs}$ | 1.46 | ACS acetate maximal uptake rate $[gDW^{-1}h^{-1}]$ |
| $K_{Acs}$ | $1.20 \times 10^{-2}$ | ACS acetate half-maximal saturation $[gL^{-1}]$ |

Table : Case study ***EPHP***: Overview of the parameters in the mathematical model for an *Enhanced Production of Heterologous Proteins*, representing a *E. coli Synthetic Microbial Community*, as described by (Mauri *et al.*, 2020).

##### 6.9.2 Structural Identifiability Analysis (SIA)

In this case study, a structural identifiability analysis (SIA) could not be performed due to the discontinuity of the functions (equations 41, 44). These discontinuities present inherent challenges for performing identifiability analysis, regardless of whether symbolic or numerical methods are used. As such, the identifiability of the parameters in this model remains unresolved, highlighting the complexity of analyzing systems with piecewise or discontinuous dynamics.

##### 6.9.3 Practical Identifiability Analysis (PIA)

We addressed a parameter estimation problem involving all the observed states ($G$, $A$, $B_P$, $H$, $B_C$), with unknown initial conditions ($G_0$, $A_0$, $B_{P0}$, $H_0$, $B_{C0}$). To evaluate the performance and robustness of our estimation methods, we generated three sets of pseudo-experimental data, each incorporating different levels of Gaussian noise: 0%, 5%, and 10%. The generated data was homoscedastic and positive.

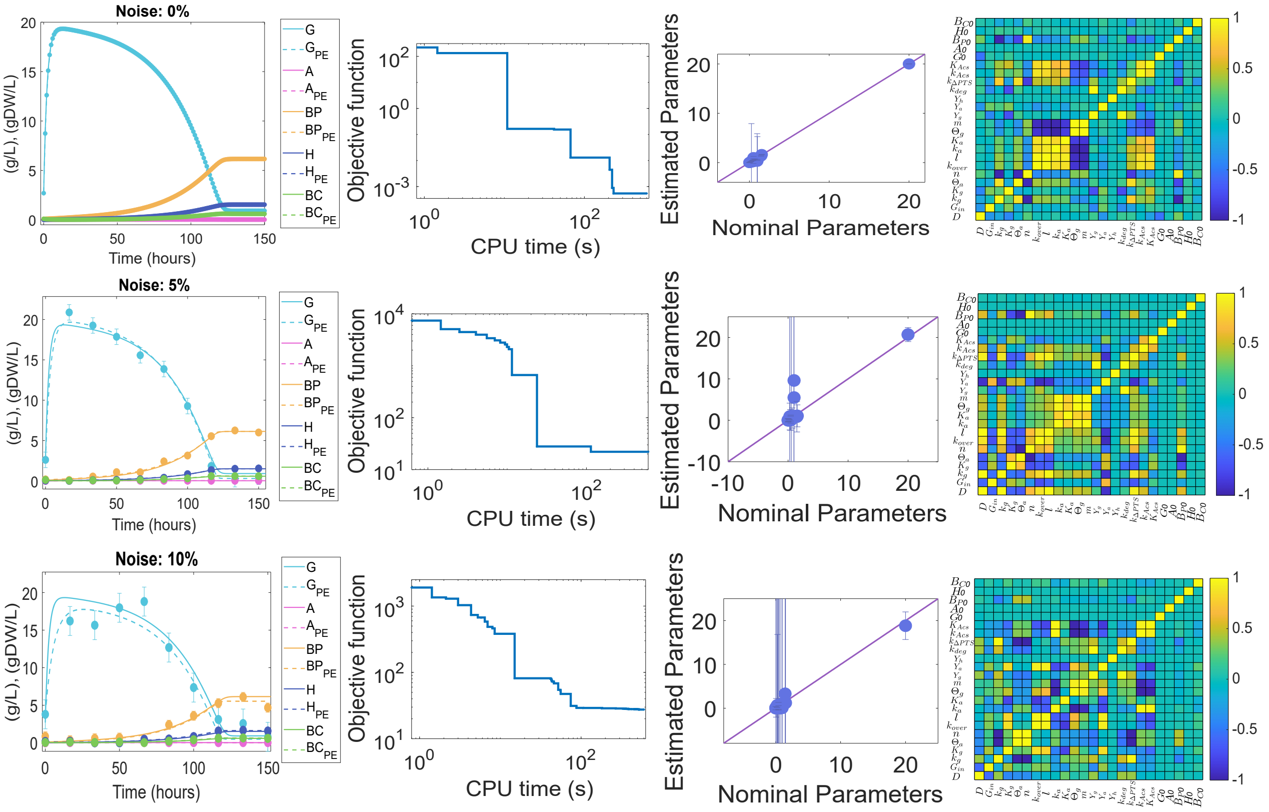

Figure : Case study ***EPHP***: Model calibration results for a synthetic microbial community of *Escherichia coli* (Mauri *et al.*, 2020). The first row displays results for the simulations with pseudo-experimental data and 0% noise, the second row shows results for 5% noise, and the third row corresponds to 10% noise. In each case, the simulation using the estimated parameters is compared to the simulation with nominal parameters, along with the convergence curve, the correlation matrix, and a comparison of nominal and estimated parameters.

After estimating the parameters, we conducted a sensitivity analysis to evaluate their influence on the population dynamics (Figure ).

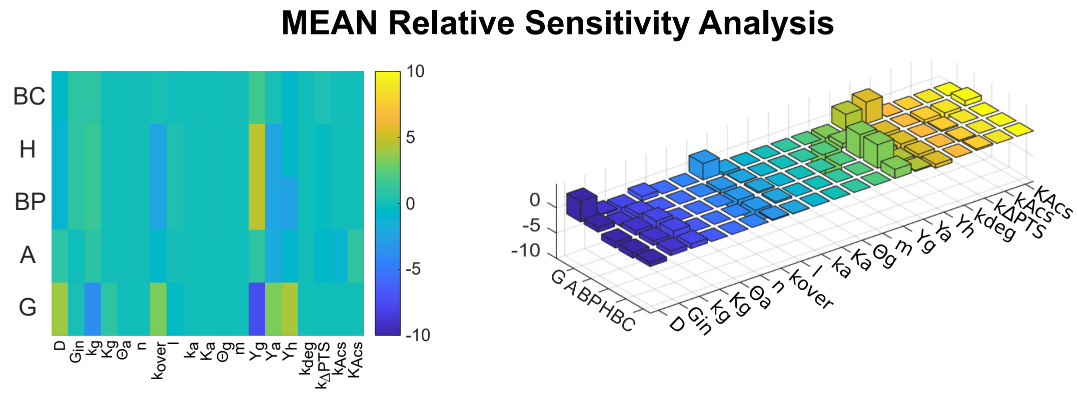

Figure : Case study ***EPHP***: Mean Sensitivity Analysis for the *Enhanced Production of Heterologous Proteins*. **Left**: Heatmap with the sensitivity of each observable (Y-axis) to each parameter (X-axis), highlighting how parameter variations affect the system's behavior. **Right**: Sensitivity analysis is presented in 3D.

| | **LB** | **UB** | **Nominal** | **Estimated** | **EA** | **ER (%)** |
|---|---|---|---|---|---|---|
| $D$ | $1 \times 10^{-2}$ | 1.00 | 0.44 | $4.24 \times 10^{-1}$ | $1.42 \times 10^{-3}$ | 3.64 |
| $G_{in}$ | 1.00 | $1 \times 10^{2}$ | $2.00 \times 10^{1}$ | $2.00 \times 10^{1}$ | $5.38 \times 10^{-3}$ | 0.00 |
| $k_g$ | $1 \times 10^{-1}$ | 10.00 | 1.53 | 1.63 | $4.90 \times 10^{-2}$ | 6.54 |
| $K_g$ | $1 \times 10^{-3}$ | $1 \times 10^{-1}$ | $9.00 \times 10^{-2}$ | $9.82 \times 10^{-2}$ | $1.45 \times 10^{-3}$ | 9.11 |
| $\Theta_a$ | $1 \times 10^{-2}$ | 1.00 | $5.20 \times 10^{-1}$ | 1.00 | $2.38 \times 10^{-1}$ | $9.23 \times 10^{1}$ |
| $n$ | $1 \times 10^{-2}$ | 1.00 | 1.00 | $5.17 \times 10^{-1}$ | $9.42 \times 10^{-2}$ | $4.83 \times 10^{1}$ |
| $k_{over}$ | $1 \times 10^{-2}$ | 1.00 | $1.70 \times 10^{-1}$ | $1.59 \times 10^{-1}$ | $8.90 \times 10^{-3}$ | 6.47 |
| $l$ | $1 \times 10^{-2}$ | 1.00 | $7.0 \times 10^{-1}$ | $6.78 \times 10^{-1}$ | $4.79 \times 10^{-2}$ | 3.14 |
| $k_a$ | $1 \times 10^{-2}$ | 1.00 | $9.70 \times 10^{-1}$ | $2.67 \times 10^{-1}$ | 5.60 | $7.25 \times 10^{1}$ |
| $K_a$ | $1 \times 10^{-2}$ | 1.00 | $5.0 \times 10^{-1}$ | $1.93 \times 10^{-1}$ | $3.61 \times 10^{-1}$ | $6.14 \times 10^{1}$ |
| $\Theta_g$ | $1 \times 10^{-1}$ | 10.00 | $2.50 \times 10^{-1}$ | $1.03 \times 10^{-1}$ | 7.81 | $5.88 \times 10^{1}$ |
| $m$ | $1 \times 10^{-2}$ | 1.00 | 1.00 | $4.95 \times 10^{-1}$ | 4.77 | $5.05 \times 10^{1}$ |
| $Y_g$ | $1 \times 10^{-2}$ | 1.00 | $4.40 \times 10^{-1}$ | $4.46 \times 10^{-1}$ | $3.56 \times 10^{-3}$ | 1.36 |
| $Y_a$ | $1 \times 10^{-2}$ | 1.00 | $2.98 \times 10^{-1}$ | $3.35 \times 10^{-1}$ | $4.34 \times 10^{-3}$ | $1.24 \times 10^{1}$ |
| $Y_h$ | $1 \times 10^{-2}$ | 1.00 | $2.00 \times 10^{-1}$ | $2.00 \times 10^{-1}$ | $5.57 \times 10^{-5}$ | 0.00 |
| $k_{deg}$ | $1 \times 10^{-4}$ | $1 \times 10^{-2}$ | $4.40 \times 10^{-3}$ | $9.96 \times 10^{-3}$ | $3.47 \times 10^{-3}$ | $1.26 \times 10^{2}$ |
| $k_{\Delta PTS}$ | $1 \times 10^{-2}$ | 1.00 | $3.80 \times 10^{-1}$ | $3.76 \times 10^{-1}$ | $9.80 \times 10^{-3}$ | 1.05 |
| $k_{Acs}$ | $1 \times 10^{-1}$ | $1.000 \times 10^{1}$ | 1.46 | 1.35 | $4.63 \times 10^{-2}$ | 7.53 |
| $K_{Acs}$ | $1 \times 10^{-3}$ | $1 \times 10^{-1}$ | $1.20 \times 10^{-2}$ | $1.27 \times 10^{-2}$ | $4.82 \times 10^{-4}$ | 5.83 |
| $G_0$ | 0.00 | 10.00 | 2.70 | 2.86 | $2.93 \times 10^{-2}$ | 5.93 |
| $A_0$ | 0.00 | 1.00 | 0.00 | $1.41 \times 10^{-6}$ | $5.79 \times 10^{-5}$ | – |
| $B_{P0}$ | 0.00 | 1.00 | $1.00 \times 10^{-1}$ | $7.86 \times 10^{-2}$ | $3.56 \times 10^{-3}$ | $2.14 \times 10^{1}$ |
| $H_0$ | 0.00 | 1.00 | 0.00 | $2.44 \times 10^{-3}$ | $2.00 \times 10^{-3}$ | – |
| $B_{C0}$ | 0.00 | 1.00 | $1.00 \times 10^{-1}$ | $9.85 \times 10^{-2}$ | $7.47 \times 10^{-4}$ | 1.50 |

Table : Case study ***EPHP***: Results for the *Enhanced Production of Heterologous Proteins* model with 0% homoscedastic Gaussian noise. The table represents the lower and upper bounds (**LB**, **UB**), nominal and estimated values, as well as the absolute and relative errors (**EA**, **ER**), for each estimated parameter.

| | **LB** | **UB** | **Nominal** | **Estimated** | **EA** | **ER (%)** |
|---|---|---|---|---|---|---|
| $D$ | $1 \times 10^{-2}$ | 1.00 | 0.44 | $2.94 \times 10^{-1}$ | $9.67 \times 10^{-1}$ | $3.32 \times 10^{1}$ |
| $G_{in}$ | 1.00 | $1 \times 10^{2}$ | 20.00 | $2.07 \times 10^{1}$ | 1.69 | $9.18 \times 10^{1}$ |
| $k_g$ | $1 \times 10^{-1}$ | 10.00 | 1.53 | 1.08 | 2.65 | $2.94 \times 10^{1}$ |
| $K_g$ | $1 \times 10^{-3}$ | $1 \times 10^{-1}$ | $9.00 \times 10^{-2}$ | $3.58 \times 10^{-2}$ | $2.43 \times 10^{-1}$ | $6.02 \times 10^{1}$ |
| $\Theta_a$ | $1 \times 10^{-2}$ | 1.00 | $5.20 \times 10^{-1}$ | $1.32 \times 10^{-1}$ | $1.27 \times 10^{2}$ | $7.46 \times 10^{1}$ |
| $n$ | $1 \times 10^{-2}$ | 1.00 | 1.00 | 5.49 | $4.43 \times 10^{3}$ | $4.49 \times 10^{2}$ |
| $k_{over}$ | $1 \times 10^{-2}$ | 1.00 | $1.70 \times 10^{-1}$ | $6.71 \times 10^{-2}$ | $1.96 \times 10^{-1}$ | $6.05 \times 10^{1}$ |
| $l$ | $1 \times 10^{-2}$ | 1.00 | $7.0 \times 10^{-1}$ | $5.42 \times 10^{-1}$ | 2.45 | $2.26 \times 10^{1}$ |
| $k_a$ | $1 \times 10^{-2}$ | 1.00 | $9.70 \times 10^{-1}$ | 1.00 | $1.59 \times 10^{11}$ | 3.09 |
| $K_a$ | $1 \times 10^{-2}$ | 1.00 | $5.0 \times 10^{-1}$ | 1.00 | $1.96 \times 10^{10}$ | $1.00 \times 10^{2}$ |
| $\Theta_g$ | $1 \times 10^{-1}$ | 10.00 | $2.50 \times 10^{-1}$ | $1.00 \times 10^{-2}$ | $1.61 \times 10^{8}$ | $9.6 \times 10^{1}$ |
| $m$ | $1 \times 10^{-2}$ | 1.00 | 1.00 | 9.62 | $1.38 \times 10^{11}$ | $8.62 \times 10^{2}$ |
| $Y_g$ | $1 \times 10^{-2}$ | 1.00 | $4.40 \times 10^{-1}$ | $4.06 \times 10^{-1}$ | $5.87 \times 10^{-1}$ | 7.73 |
| $Y_a$ | $1 \times 10^{-2}$ | 1.00 | $2.98 \times 10^{-1}$ | $9.13 \times 10^{-1}$ | 3.23 | $2.07 \times 10^{2}$ |
| $Y_h$ | $1 \times 10^{-2}$ | 1.00 | $2.00 \times 10^{-1}$ | $1.96 \times 10^{-1}$ | $1.23 \times 10^{-2}$ | 2.00 |
| $k_{deg}$ | $1 \times 10^{-4}$ | $1 \times 10^{-2}$ | $4.40 \times 10^{-3}$ | $2.22 \times 10^{-4}$ | $4.29 \times 10^{-1}$ | $9.50 \times 10^{1}$ |
| $k_{\Delta PTS}$ | $1 \times 10^{-2}$ | 1.00 | $3.80 \times 10^{-1}$ | $9.77 \times 10^{-2}$ | 2.50 | $7.43 \times 10^{1}$ |
| $k_{Acs}$ | $1 \times 10^{-1}$ | $1.00 \times 10^{1}$ | 1.46 | $8.79 \times 10^{-1}$ | 3.69 | $3.98 \times 10^{1}$ |
| $K_{Acs}$ | $1 \times 10^{-3}$ | $1 \times 10^{-1}$ | $1.20 \times 10^{-2}$ | $5.45 \times 10^{-2}$ | $3.77 \times 10^{-1}$ | $3.54 \times 10^{2}$ |
| $G_0$ | 0.00 | 10.00 | 2.70 | 2.61 | 1.89 | 3.33 |
| $A_0$ | 0.00 | 1.00 | 0.00 | $2.16 \times 10^{-3}$ | $3.33 \times 10^{-3}$ | – |
| $B_{P0}$ | 0.00 | 1.00 | $1.00 \times 10^{-1}$ | $1.63 \times 10^{-1}$ | $3.91 \times 10^{-1}$ | $6.30 \times 10^{1}$ |
| $H_0$ | 0.00 | 1.00 | 0.00 | $4.43 \times 10^{-21}$ | $1.51 \times 10^{-1}$ | – |
| $B_{C0}$ | 0.00 | 1.00 | $1.00 \times 10^{-1}$ | $1.22 \times 10^{-1}$ | $5.92 \times 10^{-2}$ | $2.20 \times 10^{1}$ |

Table : Case study ***EPHP***: Results for the *Enhanced Production of Heterologous Proteins* model with 5% homoscedastic Gaussian noise. The table displays the lower and upper bounds (**LB**, **UB**), nominal and estimated values, as well as the absolute and relative errors (**EA**, **ER**), for each estimated parameter.

| | LB | UB | Nominal | Estimated | EA | ER (%) |
|---|---|---|---|---|---|---|
| $D$ | $1 \times 10^{-2}$ | 1.00 | 0.44 | $1.76 \times 10^{-1}$ | $4.04 \times 10^{-1}$ | $6.00 \times 10^{1}$ |
| $G_{in}$ | 1.00 | $1 \times 10^{2}$ | 20.00 | $1.88 \times 10^{1}$ | 3.15 | 6.00 |
| $k_g$ | $1 \times 10^{-1}$ | 10.00 | 1.53 | 1.22 | $8.58 \times 10^{3}$ | $2.03 \times 10^{1}$ |
| $K_g$ | $1 \times 10^{-3}$ | $1 \times 10^{-1}$ | $9.00 \times 10^{-2}$ | $1.00 \times 10^{-1}$ | 2.12 | $1.11 \times 10^{1}$ |
| $\Theta_a$ | $1 \times 10^{-2}$ | 1.00 | $5.20 \times 10^{-1}$ | $2.47 \times 10^{-1}$ | $3.35 \times 10^{4}$ | $5.25 \times 10^{1}$ |
| $n$ | $1 \times 10^{-2}$ | 1.00 | 1.00 | $1.01 \times 10^{-1}$ | $8.94 \times 10^{2}$ | $8.99 \times 10^{1}$ |
| $k_{over}$ | $1 \times 10^{-2}$ | 1.00 | $1.70 \times 10^{-1}$ | $4.20 \times 10^{-1}$ | $1.38 \times 10^{2}$ | $1.47 \times 10^{2}$ |
| $l$ | $1 \times 10^{-2}$ | 1.00 | $7.0 \times 10^{-1}$ | $1.0047 \times 10^{-2}$ | $1.40 \times 10^{2}$ | $9.86 \times 10^{1}$ |
| $k_a$ | $1 \times 10^{-2}$ | 1.00 | $9.70 \times 10^{-1}$ | 1.00 | $3.0088 \times 10^{4}$ | 3.09 |
| $K_a$ | $1 \times 10^{-2}$ | 1.00 | $5.0 \times 10^{-1}$ | $5.92 \times 10^{-2}$ | $3.66 \times 10^{1}$ | $8.82 \times 10^{1}$ |
| $\Theta_g$ | $1 \times 10^{-1}$ | 10.00 | $2.50 \times 10^{-1}$ | $6.62 \times 10^{-1}$ | $3.98 \times 10^{5}$ | $1.65 \times 10^{2}$ |
| $m$ | $1 \times 10^{-2}$ | 1.00 | 1.00 | $1.00 \times 10^{-1}$ | $2.66 \times 10^{3}$ | $9.00 \times 10^{1}$ |
| $Y_g$ | $1 \times 10^{-2}$ | 1.00 | $4.40 \times 10^{-1}$ | $4.48 \times 10^{-1}$ | $8.80 \times 10^{-1}$ | 1.82 |
| $Y_a$ | $1 \times 10^{-2}$ | 1.00 | $2.98 \times 10^{-1}$ | $1.66 \times 10^{-1}$ | $1.67 \times 10^{1}$ | $4.43 \times 10^{1}$ |
| $Y_h$ | $1 \times 10^{-2}$ | 1.00 | $2.00 \times 10^{-1}$ | $2.26 \times 10^{-1}$ | $2.71 \times 10^{-2}$ | $1.30 \times 10^{1}$ |
| $k_{deg}$ | $1 \times 10^{-4}$ | $1 \times 10^{-2}$ | $4.40 \times 10^{-3}$ | $1.00 \times 10^{-2}$ | $3.76 \times 10^{-1}$ | $1.27 \times 10^{2}$ |
| $k_{\Delta PTS}$ | $1 \times 10^{-2}$ | 1.00 | $3.80 \times 10^{-1}$ | $2.88 \times 10^{-1}$ | $2.014 \times 10^{3}$ | $2.42 \times 10^{1}$ |
| $k_{Acs}$ | $1 \times 10^{-1}$ | $1.000 \times 10^{1}$ | 1.46 | 3.30 | $5.49 \times 10^{2}$ | $1.26 \times 10^{2}$ |
| $K_{Acs}$ | $1 \times 10^{-3}$ | $1 \times 10^{-1}$ | $1.20 \times 10^{-2}$ | $1.00 \times 10^{-1}$ | $3.01 \times 10^{1}$ | $7.33 \times 10^{2}$ |
| $G_0$ | 0.00 | 10.00 | 2.70 | 3.69 | 3.78 | $3.67 \times 10^{1}$ |
| $A_0$ | 0.00 | 1.00 | 0.00 | $2.95 \times 10^{-3}$ | $6.67 \times 10^{-3}$ | – |
| $B_{P0}$ | 0.00 | 1.00 | $1.00 \times 10^{-1}$ | $1.20 \times 10^{-1}$ | 1.08 | $2.00 \times 10^{1}$ |
| $H_0$ | 0.00 | 1.00 | 0.00 | $1.92 \times 10^{-1}$ | $3.02 \times 10^{-1}$ | – |
| $B_{C0}$ | 0.00 | 1.00 | $1.00 \times 10^{-1}$ | $3.09 \times 10^{-2}$ | $1.17 \times 10^{-1}$ | $6.91 \times 10^{1}$ |

Table : Case study ***EPHP***: Results for the *Enhanced Production of Heterologous Proteins* model with 10% homoscedastic Gaussian noise. The table displays the lower and upper bounds (**LB**, **UB**), nominal and estimated values, as well as the absolute and relative errors (**EA**, **ER**), for each estimated parameter.

##### 6.9.4 Estimation using multistart of local methods

To highlight the advantages of using a global optimization strategy (`eSS`) over a multistart approach with a local solver, we present below the results obtained for this case study. The histogram illustrates the distribution of objective function values obtained from multiple runs.

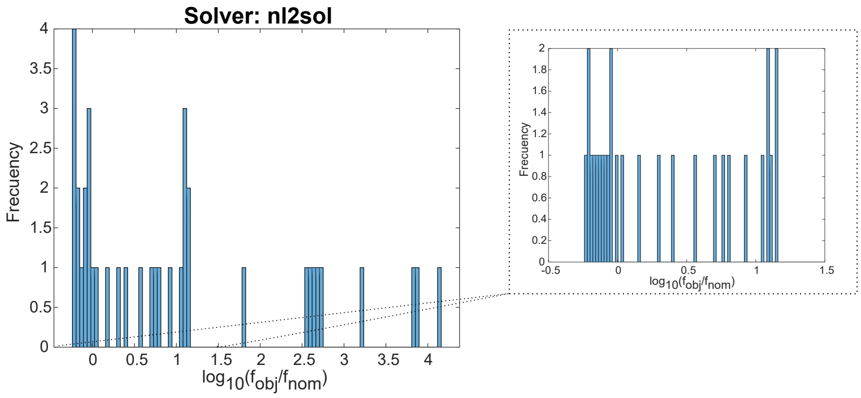

Figure : Case study ***EPHP***: Distribution of objective function values from 100 runs using the `nl2sol` local solver. The `nl2sol` solver converged to 100 optima out of 100 runs, 13 out of 100 runs resulted in a logarithm of the normalized objective function value ($log_{10}(f_{obj}/f_{nom})$) below 0, whereas 87 runs produced higher values.

#### 6.10 *MGLV*: *Generalized Lotka-Volterra* 12 species (Gut Microbiome)

##### 6.10.1 Model

The *Generalized Lotka-Volterra* (***GLV***) model is widely used for describing the dynamics of complex microbial communities, such as those found in the human gut microbiome. A synthetic ecology encompassing prevalent human-associated intestinal species: Bacteroides thetaiotaomicron ($BT$), Bacteroides ovatus ($BO$), Bacteroides uniformis ($BU$), Bacteroides vulgatus ($BV$), Blautia hydrogenotrophica ($BH$), Collinsella aerofaciens ($CA$), Clostridium hiranonis ($CH$), Desulfovibrio piger ($DP$), Eggerthella lenta ($EL$), Eubacterium rectale ($ER$), Faecalibacterium prausnitzii ($FP$), and Prevotella copri ($PC$), was designed to mirror the functional and phylogenetic diversity of the natural system, employing a system of 12 *Generalized Lotka-Volterra* equations. This model is based on a system of differential equations that captures the interactions between microbial species and the effects of external factors.

Specifically, the 12-species ***GLV*** model proposed by Venturelli *et al.* (2018) describes the rate of change in the relative abundance of each microbial species ($X_i$) as a function of its intrinsic growth rate ($\mu_i$) and its pairwise interactions with other species in the community ($\beta_{ij}$). The mathematical formulation of the model is expressed as:

$$\frac{dX_i}{dt} = \mu_i \cdot X_i + X_i \cdot \sum_{j=1}^{12} \beta_{ij} X_j \quad (i = 1, ..., 12) \tag{45}$$

This approach allows for the exploration of both the individual growth dynamics of each species and the cooperative or competitive interactions within the community.

##### 6.10.2 Structural Identifiability Analysis (SIA)

In this context, performing a structural identifiability analysis (SIA) is essential to determine whether the model's parameters can be uniquely estimated from the observed data. This step is crucial to ensuring that the inferences and predictions derived from the model are reliable and scientifically meaningful. In this case study, a fully observed model with unknown initial conditions was considered (Table ). `GenSSI2` was unable to perform the analysis due to memory limitations, resulting in an "out of memory" error. In contrast, `Structural Identifiability` tool successfully completed the analysis, delivering results 22 times faster than `SIAN`. This significant improvement in computational efficiency highlights the advantages of using `Structural Identifiability` for models with complex dynamics, where memory constraints may otherwise limit the applicability of other tools.

| | `GenSSI2` | | | | `SIAN` & `Structural Identifiability` | | | Time [min] | |
|---|---|---|---|---|---|---|---|---|---|
| Obs. | G | L | Non-Id | Time [min] | G | L | Non-Id | `SIAN` | `SI.jl` |
| $X_i$ $(i = 1, ..., 12)$ | - | - | - | ** | All | - | - | 335.20 | 15.16 |

Table : Case study ***MGLV***: Structural identifiability analysis (SIA) conducted using three tools, `GenSSI2` (**Left**), `Structural Identifiability` and `SIAN` (**Right**). The analysis was conducted for a fully observed system with unknown initial conditions. The parameters are classified as globally identifiable ("G"), locally identifiable ("L"), or non-identifiable ("Non-Id"). The entry *All* in the table indicates that all parameters in that category are identifiable as either globally, locally, or non-identifiable. A " - " indicates that no parameters are found in that category.

##### 6.10.3 Parameter bounds and stability

To ensure numerical stability and obtain physically meaningful parameter estimates, all parameters were constrained to be positive. This approach improves the numerical stability of the optimization process, as optimization algorithms generally perform better when the parameters are log-transformed, particularly when the parameters have a very large range.

Instead of directly optimizing the parameters, their logarithms were optimized, which facilitates the search for a minimum by transforming a potentially ill-conditioned problem into a more numerically stable one. To improve the numerical conditioning of the system, a constant value of 5.00 was added to all nominal parameters. This transformation resulted in the following reformulated system of ODEs:

$$\frac{dX_i}{dt} = X_i \left( (\mu_i - \alpha) + \sum_{j=1}^{12} (\beta_{ij} - \alpha) X_j \right) \text{where } \alpha = 5.00 \tag{46}$$

To enhance the robustness of the optimization process, the parameter estimation bounds were constrained. The upper bound ($UB$) for each parameter was set to 1.5 times its nominal value, while the lower bound ($LB$) was set to 0.5 times its nominal value.

This restriction was introduced to mitigate the issue of a limited number of valid solutions during the initial sampling phase of the optimization algorithm, which became particularly problematic when wider parameter bounds were considered. It was observed that expanding the bounds beyond this range resulted in an insufficient number of valid solutions to populate the initial population matrix. This behavior suggests that an excessively large search space can hinder the algorithm's ability to effectively explore the parameter space and identify feasible solutions within the allocated computational budget (1560 evaluations in this case).

##### 6.10.4 Practical Identifiability Analysis (PIA)

We addressed a parameter estimation problem for the *Human Gut Microbiome* model (***MGLV***) considering a fully observed scenario with known initial conditions (Figure ). To assess the robustness of our estimation methods, we generated three sets of pseudo-experimental data, each with different levels of Gaussian noise: 0%, 5%, and 10%. The generated data was homoscedastic. For the dataset with 0% noise, a weighted least squares approach was used as the cost function for parameter estimation. In contrast, for the 5% and 10% noise levels, a negative log-likelihood cost function was employed, as it is better suited for handling data with higher noise levels.

The impact of noise on parameter estimation was analyzed, with a particular focus on sign changes relative to the nominal values (see Figure ). The sign of these parameters is critical, as it determines whether microbial interactions are positive (mutualistic) or negative (competitive).

Notably, even in the absence of noise (0%), 48 interaction coefficients ($\beta_{ii}$, $\beta_{ij}$) exhibited sign changes compared to their nominal values (32.33% of all interactions). When the introduction of 5% noise, this number increased to 75 parameters (52.08%), with 10% noise to 77 parameters (53.47%). These findings highlight the sensitivity of parameter estimation to noise, emphasizing that higher noise levels can compromise the stability and reliability of the estimated parameters. This effect is particularly concerning in dynamic systems like the ***MGLV*** model, where precise parameter estimation is crucial for capturing the underlying biological interactions.

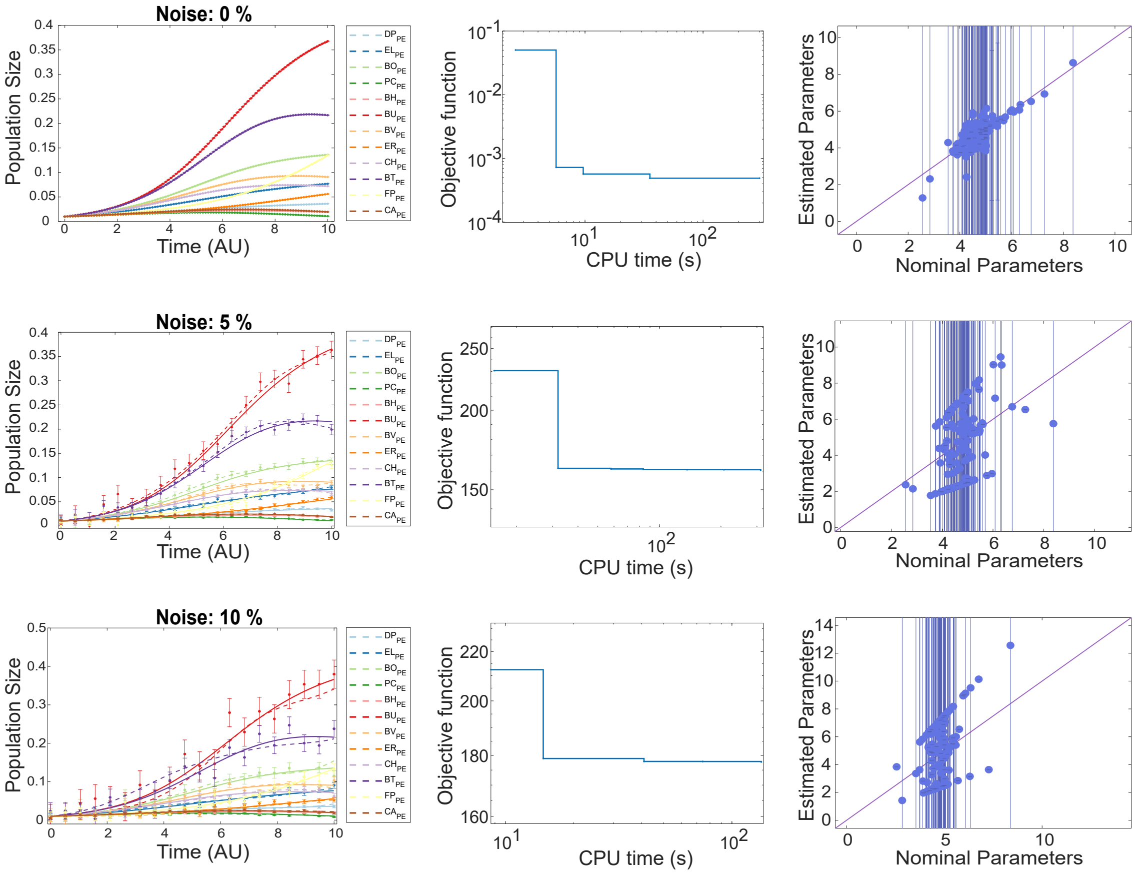

Figure : Case study ***MGLV***: Model calibration results for the *Human Gut Microbiome* model (Venturelli *et al.*, 2018). The first row displays results for the simulations with pseudo-experimental data with 0% noise, the second row shows results for 5% noise, and the third row corresponds to 10% noise. In each case, the simulation using the estimated parameters is compared to the simulation with nominal parameters, along with the convergence curve and a comparison of nominal and estimated parameters.

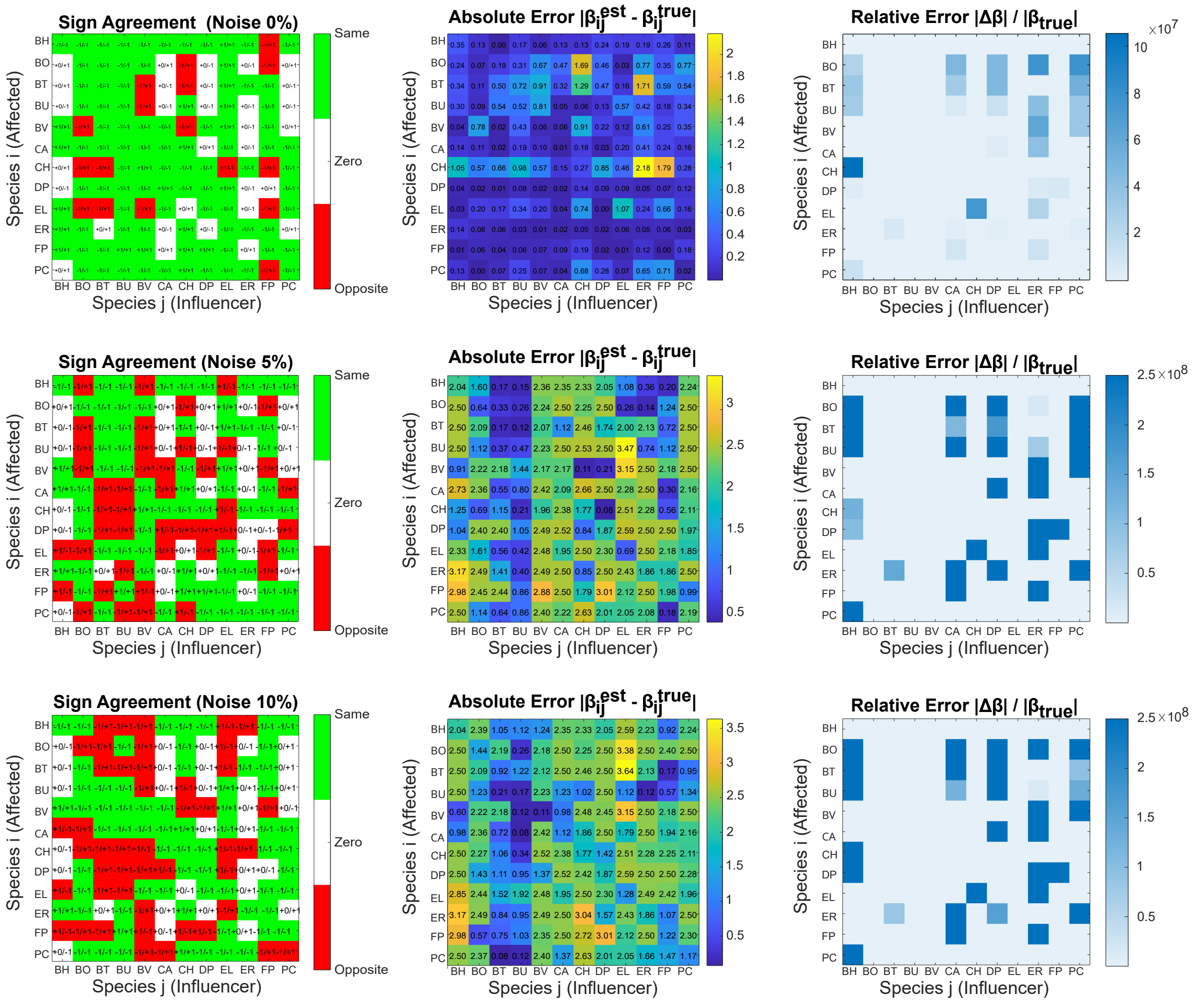

Figure : Case study ***MGLV***: Comparison of species interaction signs between estimated and nominal values $(\beta_{ii},\ \beta_{ij})$ in the *Human Gut Microbiome* model (Venturelli *et al.*, 2018). **Left:** Sign agreement. **Center:** Absolute error. **Right:** Relative error.

##### 6.10.5 optimization performance and challenges

In Figure , a comparison is presented based on 100 runs using the `eSS`, `nl2sol`, and `lsqnonlin` solvers. Notably, for the `nl2sol` and `lsqnonlin` solvers, the majority of runs failed to converge. Among them, `nl2sol` exhibited the worst performance, yielding only a single solution near its nominal value.

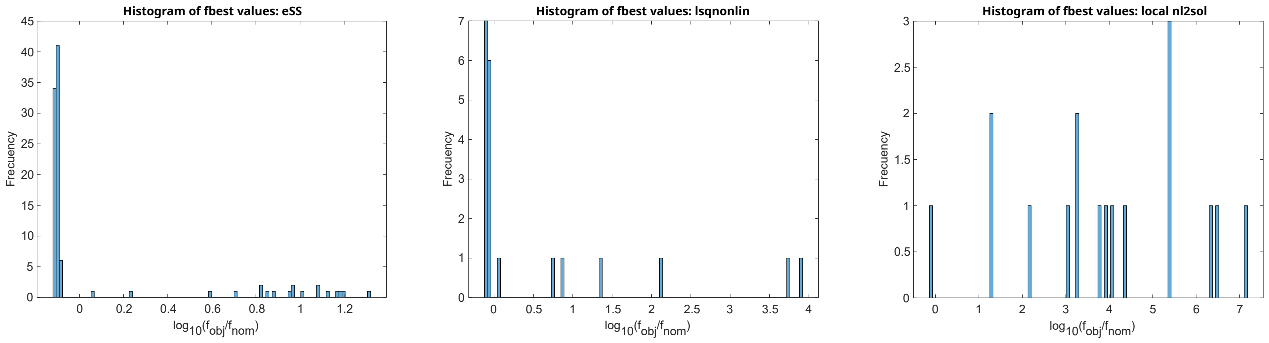

Figure : Case study ***MGLV***: Histograms showing the results of three multistart strategies with different solvers for the *Human Gut Microbiome* model, `Enhanced Scatter Search` (`eSS`) (**Left**), `lsqnonlin` (**Center**), and `nl2sol` (**Right**). The `eSS` solver converged to 100 optima out of 100 runs (81 resulted in a logarithm of the normalized objective function value ($log_{10}(f_{obj}/f_{nom})$) below 0, 19 above). In contrast, `lsqnonlin` and `nl2sol` converged to 20 (13 below 0, 7 above) and 17 optima (1 below 0, 16 above), respectively.

A post-analysis was conducted on the results from different multistarts runs to evaluate the fitting quality. The evaluation metrics included the best objective function value ($f_{best}$), root mean squared error ($RMSE$), and normalized root mean squared error ($NRMSE$). Figure  presents an example of overfitting and underfitting, while the corresponding results are summarized in Table .

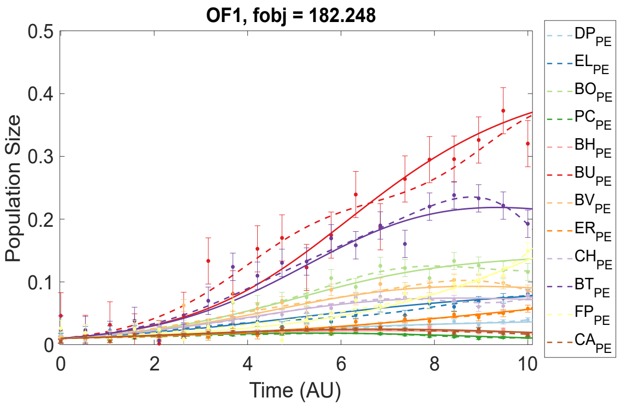

Figure : Case study ***MGLV***: Examples of overfitting (**Left**) and underfitting (**Right**) employing pseudo-experimental data with 10% Gaussian noise and the `eSS` solver. The solid line represents the simulation with nominal parameters. The objective function for the nominal values is: 237.925.

| Metric | OF1 | UF |
|---|---|---|
| $f_{best}$ | 182.2476 | 405.8853 |
| $RMSE$ | [0.0033, 0.0052, 0.0093, 0.0014, 0.0022, 0.0314, 0.0103, 0.0038, 0.0048, 0.0193, 0.0138, 0.0027] | [0.0035, 0.0063, 0.0400, 0.0017, 0.0023, 0.0380, 0.0117, 0.0039, 0.0062, 0.0303, 0.0151, 0.0027] |
| $NRMSE$ | [0.1006, 0.0734, 0.0717, 0.1267, 0.1311, 0.0847, 0.1080, 0.0752, 0.0662, 0.0834, 0.0976, 0.1268] | [0.1058, 0.0889, 0.3101, 0.1500, 0.1359, 0.1023, 0.1232, 0.0783, 0.0856, 0.1309, 0.1068, 0.1284] |

Table : Case study ***MGLV***: Post-analysis metrics comparing the overfitting (**OF1**) and underfitting (**UF**) of Figure . Results include the best objective function value ($f_{best}$), root-mean-square error ($RMSE$), and normalized root-mean-square error ($NRMSE$) for the two scenarios.

##### 6.10.6 Predictive power

As in the previous section of the ***GLV3*** model, we examine an overfitting scenario and evaluate its predictive accuracy under altered initial conditions. The selected overfitting for the study of the predictive power is shown in Figure .

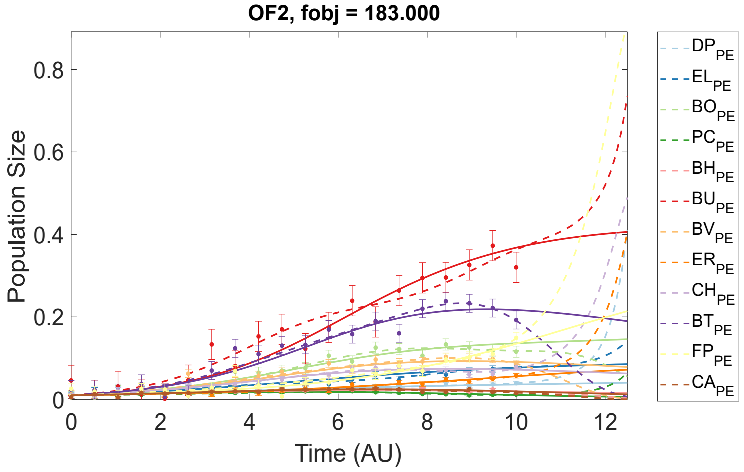

Figure : Case study ***MGLV***: Example of overfitting (**OF2**) (dashed lines) compared with the fit using nominal parameters (solid line). The post-analysis metrics are: $f_{best} = 183.0004$, $RMSE = [0.0033, 0.0052, 0.0092, 0.0014, 0.0022, 0.0313, 0.0103, 0.0038, 0.0049, 0.0196, 0.0138, 0.0027]$, $NRMSE = [0.1012, 0.0732, 0.0715, 0.1254, 0.1318, 0.0845, 0.1080, 0.0753, 0.0678, 0.0845, 0.0973, 0.1269]$.

The new set of initial conditions is: $DP_0 = 0.01$, $EL_0 = 0.02$, $BO_0 = 0.06$, $PC_0 = 0.01$, $BH_0 = 0.005$, $BU_0 = 0.005$, $BV_0 = 0.005$, $ER_0 = 0.001$, $CH_0 = 0.03$, $BT_0 = 0.001$, $FP_0 = 0.001$, $CA_0 = 0.001$. When applying the parameters from the overfitted scenario to these new initial conditions, the resulting fit is entirely inaccurate (Figure , Right). This underscores the limitations of overfitted models, as they fail to generalize to conditions outside the training dataset.

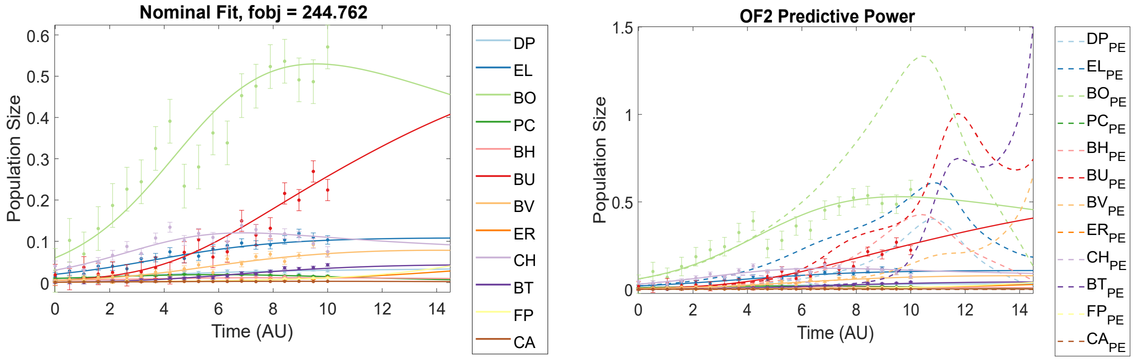

Figure : Case study ***MGLV***: Simulation employing nominal parameters with updated initial conditions (**Left**). Comparison of simulation results using parameters from overfitting scenario 2 (**OF2**) (dashed line) versus those obtained with nominal parameters (solid line), both under the updated initial conditions (**Right**).

### References

Alsolami, A. A. and El Hajji, M. (2023). Mathematical analysis of a bacterial competition in a continuous reactor in the presence of a virus. *Mathematics*, **11**(4).

Balsa-Canto, E., Alonso, A. A., and Banga, J. R. (2010). An iterative identification procedure for dynamic modeling of biochemical networks. *BMC systems biology*, **4**, 1–18.

Balsa-Canto, E., Henriques, D., Gábor, A., and Banga, J. R. (2016). AMIGO2, a toolbox for dynamic modeling, optimization and control in systems biology. *Bioinformatics*, **32**(21), 3357–3359.

Banga, J. R. and Balsa-Canto, E. (2008). Parameter estimation and optimal experimental design. *Essays in biochemistry*, **45**, 195–210.

Bellu, G., Saccomani, M. P., Audoly, S., and D'Angiò, L. (2007). Daisy: A new software tool to test global identifiability of biological and physiological systems. *Computer methods and programs in biomedicine*, **88**(1), 52–61.

Chiş, O., Banga, J. R., and Balsa-Canto, E. (2011). Genssi: a software toolbox for structural identifiability analysis of biological models. *Bioinformatics*, **27**(18), 2610–2611.

Chis, O.-T., Banga, J. R., and Balsa-Canto, E. (2011). Structural identifiability of systems biology models: a critical comparison of methods. *PloS one*, **6**(11), e27755.

Dimas Martins, A. and Gjini, E. (2020). Modeling competitive mixtures with the lotka-volterra framework for more complex fitness assessment between strains. *Frontiers in Microbiology*, **11**.

Dong, R., Goodbrake, C., Harrington, H. A., and Pogudin, G. (2023). Differential elimination for dynamical models via projections with applications to structural identifiability. *SIAM Journal on Applied Algebra and Geometry*, **7**(1), 194–235.

Egea, J. A., Balsa-Canto, E., García, M.-S. G., and Banga, J. R. (2009). Dynamic optimization of nonlinear processes with an enhanced scatter search method. *Industrial & Engineering Chemistry Research*, **48**(9), 4388–4401.

Fröhlich, F., Theis, F. J., and Hasenauer, J. (2014). Uncertainty analysis for non-identifiable dynamical systems: Profile likelihoods, bootstrapping and more. In P. Mendes, J. O. Dada, and K. Smallbone, editors, *Computational Methods in Systems Biology*, pages 61–72, Cham. Springer International Publishing.

Goldford, J. E., Lu, N., Bajić, D., Estrela, S., Tikhonov, M., Sanchez-Gorostiaga, A., Segrè, D., Mehta, P., and Sanchez, A. (2018). Emergent simplicity in microbial community assembly. *Science*, **361**(6401), 469–474.

Hofbauer, J., Hutson, V., and Jansen, W. (1987). Coexistence for systems governed by difference equations of lotka-volterra type. *Journal of mathematical biology*, **25**(5), 553—570.

Hong, H., Ovchinnikov, A., Pogudin, G., and Yap, C. (2019). SIAN: software for structural identifiability analysis of ODE models. *Bioinformatics*, **35**(16), 2873–2874.

Karlsson, J., Anguelova, M., and Jirstrand, M. (2012). An efficient method for structural identifiability analysis of large dynamic systems. *IFAC proceedings volumes*, **45**(16), 941–946.

Li, G., Leung, C.-Y., Wardi, Y., Debarbieux, L., and Weitz, J. S. (2020). Optimizing the timing and composition of therapeutic phage cocktails: A control-theoretic approach. *Bulletin of Mathematical Biology*, **82**.

Ligon, T. S., Fröhlich, F., Chiş, O. T., Banga, J. R., Balsa-Canto, E., and Hasenauer, J. (2018). Genssi 2.0: multi-experiment structural identifiability analysis of sbml models. *Bioinformatics*, **34**(8), 1421–1423.

Ljung, L. and Glad, T. (1994). On global identifiability for arbitrary model parametrizations. *automatica*, **30**(2), 265–276.

MacArthur, R. and Levins, R. (1967). The limiting similarity, convergence, and divergence of coexisting species. *The american naturalist*, **101**(921), 377–385.

Maes, K., Chatzis, M., and Lombaert, G. (2019). Observability of nonlinear systems with unmeasured inputs. *Mechanical Systems and Signal Processing*, **130**, 378–394.

Marsland, R., Cui, W., and Mehta, P. (2020). A minimal model for microbial biodiversity can reproduce experimentally observed ecological patterns. sci. rep. 10, 3308.

Massonis, G., Villaverde, A. F., and Banga, J. R. (2022). Improving dynamic predictions with ensembles of observable models. *Bioinformatics*, **39**(1), btac755.

Mauri, M., Gouzé, J.-L., de Jong, H., and Cinquemani, E. (2020). Enhanced production of heterologous proteins by a synthetic microbial community: Conditions and trade-offs. *PLOS Computational Biology*, **16**(4), 1–30.

McCaw, J. M., Arinaminpathy, N., Hurt, A. C., McVernon, J., and McLean, A. R. (2011). A mathematical framework for estimating pathogen transmission fitness and inoculum size using data from a competitive mixtures animal model. *PLoS Computational Biology*, **7**(4), e1002026.

Meshkat, N., Kuo, C. E.-z., and DiStefano III, J. (2014). On finding and using identifiable parameter combinations in nonlinear dynamic systems biology models and combos: a novel web implementation. *PloS one*, **9**(10), e110261.

Pohjanpalo, H. (1978). System identifiability based on the power series expansion of the solution. *Mathematical biosciences*, **41**(1-2), 21–33.

Remien, C. H., Eckwright, M. J., and Ridenhour, B. J. (2021). Structural identifiability of the generalized lotka–volterra model for microbiome studies. *Royal Society Open Science*, **8**(7), 201378.

Rey Barreiro, X. and Villaverde, A. F. (2023). Benchmarking tools for a priori identifiability analysis. *Bioinformatics*, **39**(2), btad065.

Rey Rostro, D. and Villaverde, A. (2022). Strikepy: Nonlinear observability analysis of inputs, states, and parameters in python. In *XLIII Jornadas de Automática*, pages 430–435. Universidade da Coruña. Servizo de Publicacións.

Roach, D. R., Leung, C. Y., Henry, M., Morello, E., Singh, D., Di Santo, J. P., Weitz, J. S., and Debarbieux, L. (2017). Synergy between the host immune system and bacteriophage is essential for successful phage therapy against an acute respiratory pathogen. *Cell host & microbe*, **22**(1), 38–47.

Sedoglavic, A. (2001). A probabilistic algorithm to test local algebraic observability in polynomial time. In *Proceedings of the 2001 international symposium on Symbolic and algebraic computation*, pages 309–317.

Shi, X. and Chatzis, M. (2022). An efficient algorithm to test the observability of rational nonlinear systems with unmeasured inputs. *Mechanical Systems and Signal Processing*, **165**, 108345.

Succurro, A. and Ebenhöh, O. (2018). Review and perspective on mathematical modeling of microbial ecosystems. *Biochem. Soc. Trans.*, **46**(2), 403–412.

Venturelli, O. S., Carr, A. V., Fisher, G., Hsu, R. H., Lau, R., Bowen, B. P., Hromada, S., Northen, T., and Arkin, A. P. (2018). Deciphering microbial interactions in synthetic human gut microbiome communities. *Molecular systems biology*, **14**(6), e8157.

Villaverde, A. F., Barreiro, A., and Papachristodoulou, A. (2016). Structural identifiability of dynamic systems biology models. *PLoS computational biology*, **12**(10), e1005153.

Villaverde, A. F., Pathirana, D., Fröhlich, F., Hasenauer, J., and Banga, J. R. (2022). A protocol for dynamic model calibration. *Briefings in bioinformatics*, **23**(1), bbab387.

Walter, E. and Lecourtier, Y. (1982). Global approaches to identifiability testing for linear and nonlinear state space models. *Mathematics and Computers in Simulation*, **24**(6), 472–482.
